# Anchoring the mean generation time in the SEIR to mitigate biases in ℜ_0_ estimates due to uncertainty in the distribution of the epidemiological delays

**DOI:** 10.1101/2023.01.13.523965

**Authors:** Jair Andrade, Jim Duggan

## Abstract

The basic reproduction number, *ℜ*_0_, is of paramount importance in the study of infectious disease dynamics. Primarily, *ℜ*_0_ serves as an indicator of the transmission potential of an emerging infectious disease and the effort required to control the invading pathogen. However, its estimates from compartmental models are strongly conditioned by assumptions in the model structure, such as the distributions of the latent and infectious periods (epidemiological delays). To further complicate matters, models with dissimilar delay structures produce equivalent incidence dynamics. Following a simulation study, we reveal that the nature of such equivalency stems from a linear relationship between *ℜ*_0_ and the mean generation time, along with adjustments to other parameters in the model. Leveraging this knowledge, we propose and successfully test an alternative parameterisation of the SEIR model that produces accurate *ℜ*_0_ estimates regardless of the distribution of the epidemiological delays, at the expense of biases in other quantities deemed of lesser importance. We further explore this approach’s robustness by testing various transmissibility levels, generation times, and data fidelity (overdispersion). Finally, we apply the proposed approach to data from the 1918 influenza pandemic. We anticipate that this work will mitigate biases in estimating *ℜ*_0_.

## 1 Introduction

The analysis of any infectious disease’s dynamics will inevitably lead to the *basic reproduction number* (*ℜ*_0_). Initially developed in the study of demographics [1], this quantity has been interpreted in the epidemiological context as the average number of secondary infections arising from the introduction of one infected individual into a totally susceptible population [2]. The usefulness and importance of *ℜ*_0_ lie primarily in its *threshold phenomenon* [3]. That is, a pathogen can invade a totally susceptible population only if *ℜ*_0_ *>* 1 [4]. Furthermore, the magnitude of *ℜ*_0_ gauges the transmission potential of an emerging infectious disease [3] and the effort required to control the invading pathogen [5]. Thus, accurate estimation of *ℜ*_0_ is crucial for understanding and managing infectious diseases.

Another reason for the popularity of *ℜ*_0_ is that one can estimate it from epidemiological data [6] using a number of methods. For diseases that allow the assumption of endemic equilibrium and homogeneous mixing, one can follow Mollison’s method [7] or Dietz’s approach [8]. The former requires prevalence data, whereas the latter leverages readily available information such as age at infection and average life expectancy. On the other hand, if an infection leads to either immunity or death in a closed population, seroprevalence studies can inform the fraction of the population that acquired the disease during an epidemic, i.e. the *final epidemic size*. In their seminal paper, Kermack and McKendrick [9,10] formulated a relationship between the final epidemic size and *ℜ*_0_, from which the latter can be calculated. Unlike the previous methods, which require the epidemic to reach a steady state, *ℜ*_0_ may be determined from the intrinsic growth rate of the infected population [3,5] using incidence data of the early stages of the epidemic, as long as the growth of new cases exhibits pure exponential behaviour. Alternatively, we can employ the entire report of daily case notifications if *ℜ*_0_ is formulated as a function of a compartmental model’s parameters [11–13]. These models can be stochastic [14], semi-deterministic [15,16] or deterministic [9,17].

These compartmental models are said to be *mechanistic* [18], namely, structures based on a scientific understanding of infectious disease dynamics [19]. The relevance of that mechanistic property lies in the role of the model. Rather than being a merely mathematical artefact to produce a desired output, the model also embeds a dynamic hypothesis of the underlying process that generates the observed data. Hence, the parameters, states and interactions that comprise a particular formulation represent their counterparts in the real world. If the model accurately captures the properties of the actual phenomenon, finding an adequate *configuration* (assign values to parameters) should yield a behaviour over time of infections that resembles the observed trajectory. The values of such parameters can be obtained from individual-level observations [4] or via statistical inference [20–22], a process also known as *trajectory matching* or *model fitting*.

Furthermore, matching simulated and observed behaviour can be regarded as a validation test on the dynamic hypothesis that links structure to behaviour [23]. Nevertheless, one should understand this validation step as a falsification test [24]. That is, if the model fails to reproduce the observed behaviour, it can certainly be rejected. On the contrary, obtaining an accurate match (or fit) does not immediately validate the dynamic hypothesis inasmuch as there may be other competing hypotheses that fit the data equally well. Indeed, this circumstance impacts the estimation of *ℜ*_0_ from compartmental models (and the intrinsic growth rate method), where different assumptions can yield accurate fits [25]. However, estimates vary according to the specific assumptions embedded in each fitting model [3,26].

For instance, the choice of the distributions of the latent and infectious periods (epidemiological delays) in the deterministic Susceptible-Exposed-Infectious-Recovered (*SEIR*) framework plays an essential role in the inference of *ℜ*_0_ [25]. Briefly put, misspecifying the structure of such delays leads to biases in the estimates. That is, a systematic difference between true and estimated parameters. Although there are techniques [27,28] to construct models with realistic distributions, modellers do not know exactly which distribution to incorporate in their formulation. In view of this drawback, Wearing and colleagues [25] fitted various *SEIR* models (with different delay distributions) to a single incidence dataset to select the best structure based on a goodness-of-fit measure. Nevertheless, the results appear inconclusive. Notwithstanding that Krylova & Earn [29] assume their validity, no further research establishes the reliability of such an approach. This assessment immediately warrants the need for the work presented here: a systematic study oriented to determine whether it is possible to infer *ℜ*_0_ accurately from *SEIR* models fitted to incidence data in light of the uncertainty in the distributions of the epidemiological delays. We describe the steps of this study in the sections below. All the analysis is performed in R. The code is freely available at https://github.com/jandraor/delays.

## 2. Data Generating Process

### 2.1 The system (latent) component

In order to undertake a systematic study, experimenters must have access to a sizeable set of observations. In this case, multiple time series of daily case notifications of a particular disease under various conditions. Equally important, such conditions need to be known *a priori*. To meet these conditions, we leverage the mechanistic property of compartmental models and employ the *SEIR* framework as a synthetic data generator [30,31]. This framework has been widely applied to studying various infectious diseases, such as measles [29,32,33], COVID-19 [16,34,35], and influenza [12,17,22,36–38]. In this work, we restrict our attention to the simplest version of this family of models. The rationale for this decision is straightforward; conceptual models entail efficiency inasmuch as they facilitate the understanding and identification of the underlying causes of a particular result. Moreover, it is often the case that principles that stem from basic models apply to more elaborated extensions.

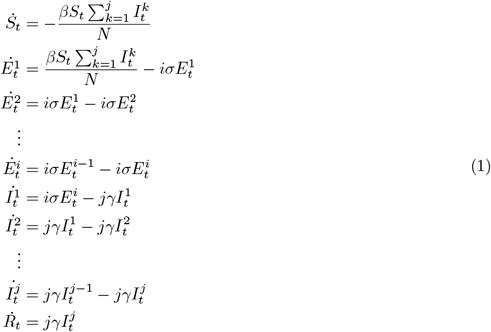

Specifically, the *SEIR* (Eq 1) stratifies individuals as susceptible (*S*_*t*_), exposed (*E*_*t*_), infectious (*I*_*t*_), and recovered (*R*_*t*_) and describes the transitions between states (*S*_*t*_ *→ E*_*t*_ *→ I*_*t*_ *→ R*_*t*_) in terms of differential equations. Susceptible individuals acquire infection, *S*_*t*_ *→ E*_*t*_, through contact with infectious individuals, where the number of contacts is independent of the population size (*N*). Formally, one refers to this assumption as the frequency-dependent (or mass action) transmission: *βS*_*t*_*I*_*t*_*/N*. Here, *β* corresponds to the effective contact rate or transmission parameter. The movement of individuals from the class *E*_*t*_ to class *R*_*t*_ is modelled using a well-known mathematical procedure [39] to achieve realistic distributions [40,41] of the time that individuals spend in states *E*_*t*_ and *I*_*t*_, otherwise known as the latent and infectious periods, respectively. Such a procedure corresponds to the subdivision of a class into stages arranged in series. For instance, one can divide the exposed class into *i* stages. Newly infected individuals enter the first exposed stage, 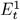, pass through each in turn and become infectious upon leaving the *ith* stage 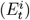. The progression between stages is assumed to occur at a constant per-capita rate (*iσ*), leading to an exponential waiting time with mean 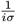 in each stage [11]. This formulation implies that the lapse between infection and becoming infectious is described by the sum of *i* independent exponential random variables with equal rates, a convolution resulting in a gamma-distributed random variable [42]. Therefore, the subdivision of the exposed class into various stages is equivalent to formulate the latent period in terms of a gamma distribution with mean *σ*^*−*1^ and shape *i*. Similarly, one can divide the infectious class into *j* stages to formulating a gamma-distributed infectious period.

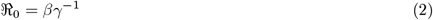

Overall, we refer to Eq 1 as the *SE*^*i*^*I*^*j*^*R* framework. Notice that the standard *SEIR* corresponds to the *SE*^1^*I*^1^*R* instance. Moreover, as the parameter *i* increases, the distribution becomes more closely centred on its mean (tighter), to the extent that if *i → ∞*, the variance is removed. That is, in the limit, all individuals have the same latent period. An equivalent argument applies to the infectious period. No less important, as indicated by [11], irrespective of the values of *i* and *j* that the *SE*^*i*^*I*^*j*^*R* may take, the basic reproduction number depends exclusively on the transmission rate and the mean infectious period (Eq 2). Furthermore, it is noteworthy to mention that subdividing a class is a mathematical device that allows the incorporation of additional distributions in a system of differential equations, and the number of stages may not correspond to biological features of the infection process [32]. Lastly, we assume that the disease leads to permanent immunity and that the outbreak’s time scale is much faster than the characteristic times for demographic processes (births and deaths), therefore their effects are not included. This last assumption implies that the population remains constant over the simulation period.

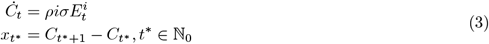

Subsequently, we define the link between the *SE*^*i*^*I*^*j*^*R* and incidence data (Eq 3). Based on the literature [15,16,22,36], we posit that incidence (*Ċ*) is proportional to the rate at which individuals become infectious 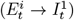. Such proportional effect or reporting rate (*ρ*) stems from the fact that individuals experience various degrees of symptom severity [43]. In particular, individuals with low severity levels (asymptomatic and mild symptoms) may not seek health care attention, resulting in case reports that most likely miss a significant fraction of infected individuals. As opposed to the continuous nature of differential equation models, case notifications occur at discrete times. To reconcile this tension, we define the report of new cases (*x*_*t\**_) as the change in the total number of cases (*C*_*t*_) in one-day intervals.

Furthermore, we tailor the synthetic data generator towards influenza given that this virus causes unpredictable but recurring pandemics that can have significant global consequences [44]. As a matter of fact, there have been four influenza pandemics over the past 100 years, including the H1N1 pandemic in 1918, with 50 estimated million deaths [45]. Adapting the *SE*^*i*^*I*^*j*^*R* framework to this choice involves the selection of plausible parameter values or ground truths [46]. For simplicity, we restrict the synthetic data generator to eight instances: *i* = *{*1, 3*} × j* = *{*1, 2, 3, 4*}*. These instances share constants *σ, γ, β, ρ*, and *N*, which are configured identically. In particular, we configure parameters *σ* and *γ* from the assumed values (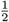 for both) in the Cumberland case study [12,22]. Following this choice, we select a value of *β* that yields a b^2^asic reproduction number (*2*.*5*) within a plausible range (*2-4*) of pandemic influenza [47]. Regarding *ρ*, we choose a value (*0*.*75*) consistent with reported estimates in the literature [12,22]. The remaining constant, *N*, has only a scaling effect, and any particular value (*10,000* in this case) does not alter the model dynamics provided that *N* = *S*_0_ + *E*_0_ + *I*_0_ + *R*_0_, where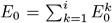 and 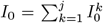. In relation to initial conditions, we assume that a patient zero triggers the outbreak of a novel influenza pathogen. In mathematical terms, *S*_0_ = *N -* 1 and 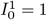. The remaining initial conditions of the within-host profile are set to zero.

Having delimited the *SE*^*i*^*I*^*j*^*R* framework and configured its instances, we run simulations (Fig 1) that illustrate the impact of the delay structure on the incidence dynamics. In agreement with the literature [4,25], note in Fig 1 that if we fix the latent period distribution (*i*) and vary that for the infectious period (*j*), incidence reports that stem from more tightly distributed infectious periods (larger *j*) reach the incidence peak earlier and end more abruptly. This difference in behaviour over time occurs despite the fact that these instances share identical *ℜ*_0_ and equal average latent and infectious periods. On the other hand, if we fix the infectious period (compare two lines of the same colour across panels), decreasing the latent period’s variance (increasing *i* from *1* to *3*) produces the opposite effect. Namely, tighter latent period distributions (larger *i*) push forward the peak time and extend the outbreak’s duration.

**Figure 1:**
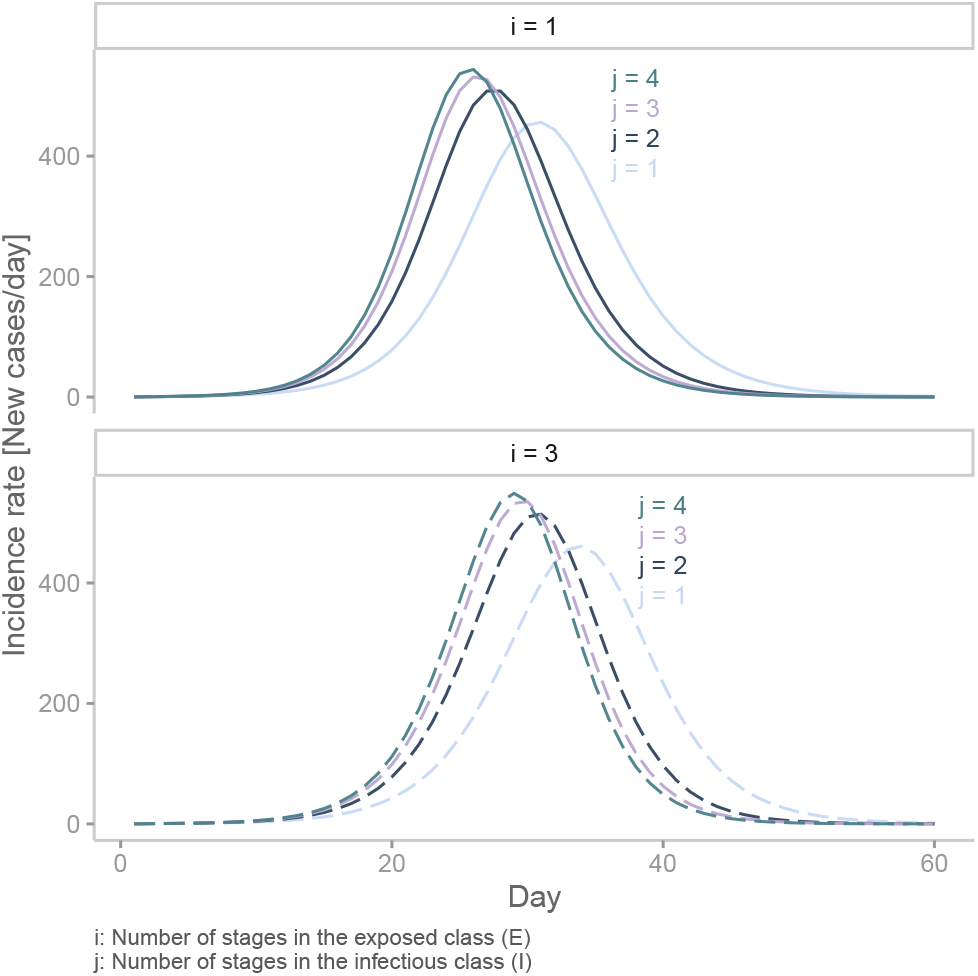
Incidence reports generated by various instances of the *SE*^*i*^*I*^*j*^*R* framework. In this plot, we present two distributions of the latent period and four distributions of the infectious period. The colour of a line corresponds to a particular value of *j* (infectious period distribution). Solid lines indicate that the incidence report stems from an SEIR model with an exponentially-distributed latent period (*i* = 1). Dashed lines indicate that the incidence report stems from an SEIR model with a gamma-distributed latent period (*i* = 3).

### 2.2 Measurement component

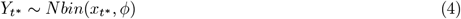

Borrowing terminology from the state-space literature [18,48], one can frame the output produced by the *SE*^*i*^*I*^*j*^*R* framework as predictions obtained from a system or *latent* component. In practice, though, continous and smooth predictions from ODE models differ from noisy and discrete incidence reports collected by public health surveillance. Moreover, given that a system component is merely a partial representation of a more complex reality, some elements are necessarily omitted. Consequently, it is required to equip the data generating process with a structure that accounts for the discrepancies between model prediction and actual data. We refer to this structure as the measurement component. In epidemiology, one can formulate the measurement of new infections via the *Negative Binomial* distribution, considering that this function does not tie the observation mean to the variance, offering the flexibility to account for overdispersion [19]. Accordingly, we define the observation of new cases (Y_*t\**_) in terms of a Negative Binomial distribution (Eq 4) specified by location (mean) and diffusion parameters. The former corresponds to the predicted incidence by the system component (*x*_*t\**_), whereas the latter (*ϕ*) modulates the concentration of measurements. Note that the inverse of the concentration parameter (*ϕ*^*−*1^) represents overdispersion inasmuch as an increase in its magnitude leads to greater diffusion in the data.

Defining a measurement component completes the formulation of the data generating process. Consequently, we draw samples from Eq 4 using statistical simulation (*rnbinom in R*). For each *SE*^*i*^*I*^*j*^*R* instance, we generate *40 noisy* time series. We perform this process for two levels (*high* and *low*) of data fidelity, a feature measured by *ϕ*^*−*1^. High-fidelity data (*ϕ*^*−*1^ = 0) implies that the measurement component applies only a slight distortion on the original signal (incidence). Notice that this configuration of the Negative Binomial (with no overdispersion) is equivalent to the *Poisson* distribution. Conversely, a positive value (overdispersion) of *ϕ*^*−*1^ (such as *1/3*) distorts the original signal to such an extent that one cannot easily discern the underlying incidence dynamics (low-fidelity data). We generated a total of *320* incidence reports, of which Fig 2 presents a sample of four representative reports. The reader can find the complete details in the electronic supplementary material S1. To facilitate the communication of results, we introduce the notation *D*^*ij*^, which indicates the origin of a given set of time series. For example, *D*^14^ indicates that the observed incidence was obtained from the *SE*^1^*I*^4^*R* instance.

**Figure 2:**
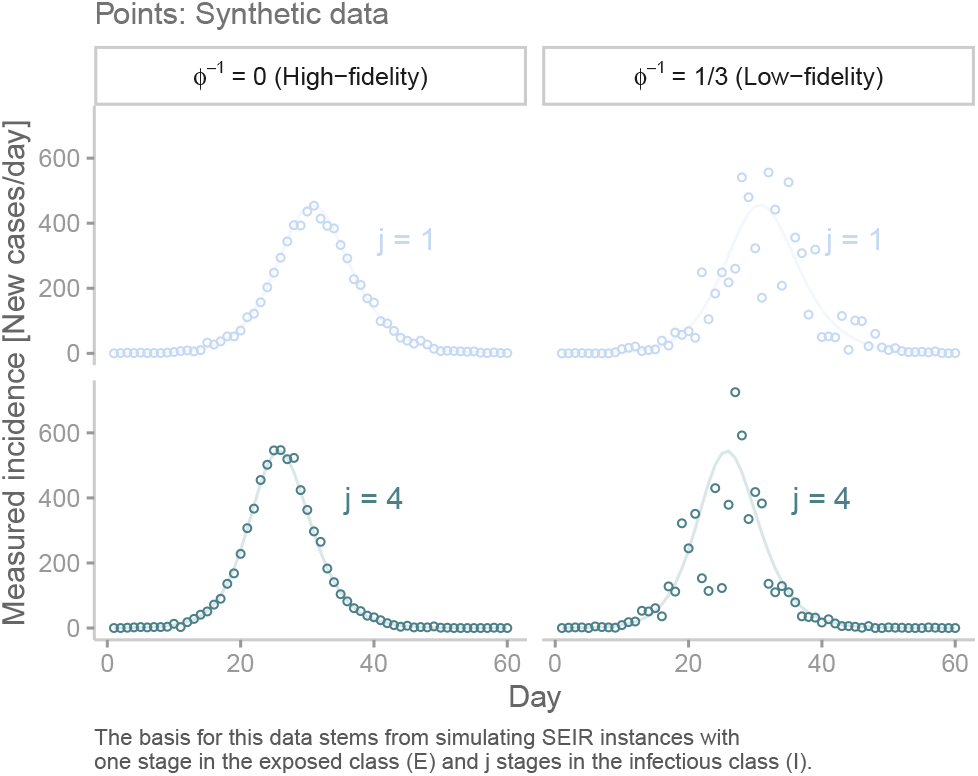
Sample of synthetic data. This plot shows four representative incidence reports (dots) obtained from the simulation of two *SE*^1^*I*^*j*^*R* instances (lines). To obtain each report, we sample from the negative binomial distribution.

## 3. Inference

The synthetic incidence reports described in the previous section allow us to assess the performance of various candidate models in recovering ground truths, particularly *ℜ*_0_, our quantity of interest. Specifically, we fit model candidates to incidence data following a *Bayesian* approach [22,49]. That is, each candidate’s unknown parameters are treated as random variables, which describe the knowledge (or uncertainty) about their actual values [50], expressed in terms of a probability distribution. This distribution is updated in light of new information summarised by a likelihood function. This function evaluates the compatibility between a given incidence report and multiple configurations of a model candidate [51]. Such updating process yields the target or posterior distribution, an information device whereby we derive answers for our inferential questions. We approximate the posterior distribution via sampling using Hamiltonian Monte Carlo or HMC [52], an algorithm successfully employed to perform statistical inference from epidemiological models [16,17,22,53,54]. This algorithm is provided by the statistical package *Stan* [55].

### 3.1 Three unknowns (traditional): *β, ρ, I*_0_

For simplicity, we initially restrict the inference analysis to *D*^1*j*^ high-fidelity observations. To fit each incidence report, we postulate four instances, *j* = *{*1, 2, 3, 4*}*, from the *SE*^1^*I*^*j*^*R* framework, which share identical mean latent and infectious periods. We refer to the approach of fixing the means of the epidemiological delays to values obtained from the literature, regardless of their distribution, as the *traditional* parameterisation. Moreover, it is assumed that the measurement component is fully known. Consequently, discrepancies between estimated and actual values are ascribed to misspecification in the infectious period distribution. To avoid confusion between the origin of data and the fitting model, we denote the latter as *M*^*ij*^. As a consequence, this design requires the estimation of *320* posterior distributions. Given this process’s computational burden, we limit the number of random variables in each model to three: the transmission rate (*β*), the reporting rate (*ρ*) and the initial number of infected individuals in stage one 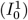. The remaining parameters and initial conditions are considered to be known, i.e. they are fixed to their actual values. Based on this setup, we fit each candidate to a given dataset using HMC sampling, with *four* Markov chains and *1000* iterations (plus *1000* for warm-up) each, checking for convergence and effective sample sizes. The complete set of results can be found in the electronic supplementary material *S2 § 1*.

The results presented in Fig 3 replicate a finding previously reported in the literature [25,56]: the existence of a subtle yet fundamental interaction between the assumed model structure and estimated *ℜ*_0_. Misspecifying the infectious period distribution with a tighter distribution (higher *j*) generates lower *ℜ*_0_ estimates (Fig 3A). Furthermore, regardless of the assumed distribution of the infectious period, all candidate models fit the data equally well. To emphasise the importance and implications of this observation, we compare inferred and actual latent incidences in Fig 3B. Recall that fitting a candidate model to a given incidence (*y*_*t*_) produces a set of samples that describes the posterior distribution. Then, we use those samples to simulate the candidate’s system component, thereby generating inferred latent incidences (lines in Fig 3B). Then, those lines are compared to *x*_*t\**_, the true latent incidence (Fig 1). Notice that by definition, we do not have access to *x*_*t\**_ in practical applications, but by virtue of this simulation study, such an impediment is overcome. The comparison reveals a symmetry shared among the candidate models. That is, any of these formulations can match the true latent incidence provided that *β, ρ* and 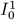 are configured appropriately. It is important to remark that this symmetry is restricted to the latent incidence and does not extend to the dynamics of other states. For instance, candidates with different delay distributions that yield equivalent incidences will not reach the same long-term equilibrium, given the differences in their *ℜ*_0_.

**Figure 3:**
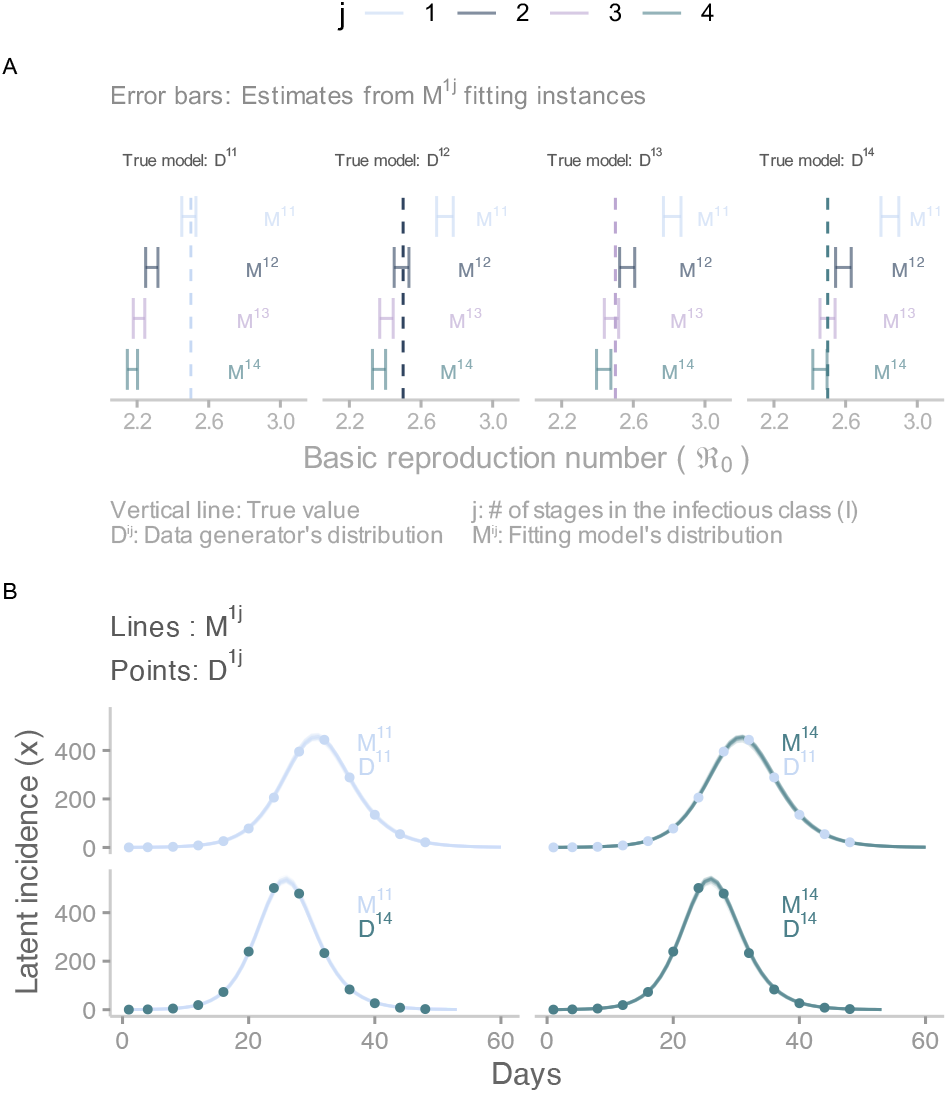
Inference results obtained from the three-unknown parameterisation. This plot shows the results of fitting model candidates to incidence reports. A) Comparison of estimates for the basic reproduction number obtained from fitting four candidate models to four incidence reports. Error bars correspond to 95% credible intervals, and the vertical line denotes the true value. B) Comparison between inferred incidence (lines) obtained from two candidate models fitted to two incidence reports (dots). Twenty time series represent inferred incidence. Given the high-fidelity data, all inferred incidences are nearly identical, giving the impression of only one line in each panel.

Logically, such symmetry should render the approach of comparing fit scores impractical. A fit score, such as the Maximum Likelihood Estimate (MLE), measures the consistency between a dataset and the output generated by a model. Since candidates produce equivalent output, differences among MLEs will solely reflect the stochasticity (noise) of the measurement component. We empirically verify this conjecture by selecting the candidate with the largest MLE for each incidence report (see the electronic supplementary material *S2 § 1*.*3*.*2*). We observe that *M* ^11^ candidates attain the largest MLE in only *12* out of *20* times when matching *D*^11^ incidence reports. Even worse, *M* ^13^ instances are always outperformed in fitting *D*^13^ datasets. Overall, no candidate passes the 60% mark. Similarly, the *mean absolute scaled error* (MASE), a metric specifically designed for evaluating the accuracy of time-series forecasts [57], indicates that candidates produce virtually identical scores when fitting any given incidence report. In light of this evidence, one can safely conclude that score comparison is not a reliable approach to determining the correct distribution of epidemiological delays from incidence data. To further complicate matters, information criteria (such as *AIC* and *BIC*) and cross-validation methods cannot assist in this task, considering that the evaluated structures produce equivalent output and share an equal number of unknown parameters.

### 3.2 Four unknowns: *β, ρ, I*_0_, *γ*

The reason for such inherent symmetry is the *generation time*, the time between the infection of a primary case and one of its secondary cases [58]. This quantity’s shape, in tandem with *ℜ*_0_, determines the initial dynamics of an infectious disease [5]. Interestingly, these elements also characterise long-term behaviour. Krylova & Earn [29] found that *SEIR* models that account for demographic processes with different delay distributions produce equivalent dynamics of epidemiological transitions (e.g. from annual to biennial epidemic cycles) if they share identical *ℜ*_0_ and mean generation time (*τ*). An analytical expression for this last quantity can obtained using the method described by Svensson [58]. In particular, for the *SE*^*i*^*I*^*j*^*R* framework, *τ* can be expressed as a function of the average delays (*σ*^*−*1^, *γ*^*−*1^) and the infectious period distribution (*j*).

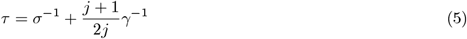

In this analysis, we have, until now, fixed the mean generation time on each candidate model by excluding *σ* and *γ* from the inference process. Taking note of the effects on short and long-term dynamics that produce the interaction between *τ* and *ℜ*_0_, we now promote *γ* to the category of *estimated parameter* in order to explore the impact of a variable mean generation time. The reason for choosing *γ* as the extra parameter is based on the fact that it interacts with both quantities of interest (Eqs 2 and 5). This choice implies the need for estimating four parameters per model instance. To do so, we follow the approach described in the previous section. The reader can find the full set of results in the electronic supplementary material *S2 § 2*. Unsurprisingly, given the extra degree of freedom, all candidates fit any of the incidence data equally well. In this design, though, the match between synthetic data and fitting model’s output is achieved at the expense of less precision, although greater accuracy. *Precision* refers to the width of uncertainty intervals, and *accuracy* to whether the interval captures the actual value. To illustrate this phenomenon, in Fig 4, we present the results of fitting four candidates models (*M* ^1*j*^) to four incidence reports that stem from different distributions of the infectious period (*D*^1*j*^). Here, we see that the range of *ℜ*_0_ widened (Fig 4A) compared to that presented in the previous section (Fig 3A).

**Figure 4:**
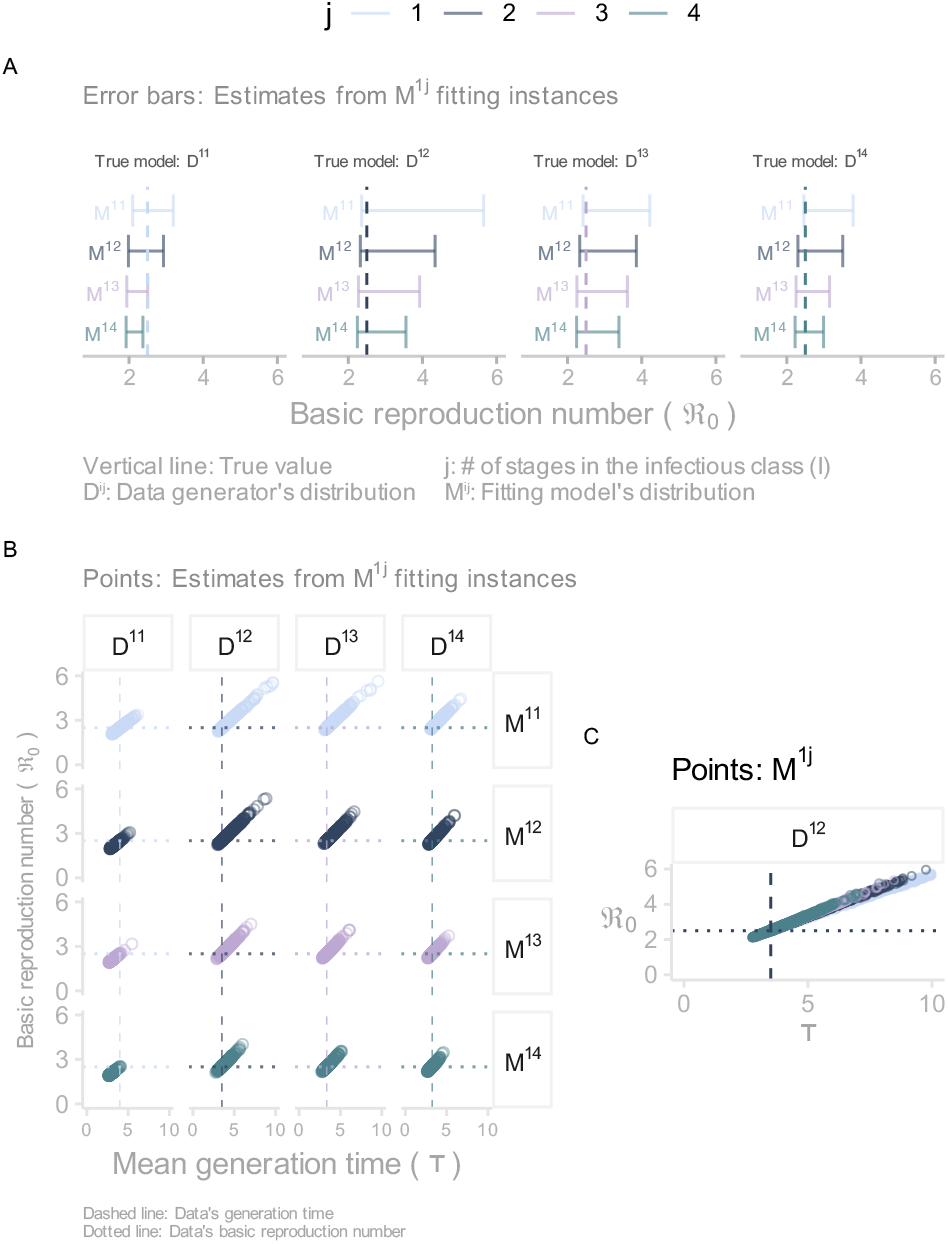
Inference results obtained from the four-unknown parameterisation. A) Comparison of estimates for the basic reproduction number obtained from fitting four candidate models to four incidence reports. Error bars correspond to 95% credible intervals, and the vertical line denotes the true value. B) Linear relationship between the basic reproduction number and the mean generation time estimated from posterior distributions obtained from fitting four candidate models to four incidence reports. These distributions are represented via samples. From each sample, we compute the predicted *ℜ*_0_ and *τ* (dots). C) This plot collapses the second column in B into a single panel.

Undoubtedly, the primary insight from allowing *γ* to vary is the unravelled interaction between *ℜ*_0_ and *τ*. We visualise this interaction by plugging samples of *β* and *γ* into Eqs 2 and 5 to obtain an approximation of the expected values of *ℜ*_0_ and *τ*. When these two quantities are displayed on a scatter plot (Fig 4B), a linear relationship appears, regardless of the data’s origin or the fitting model’s structure. The interpretation of such linear association indicates that for a given fitting model, infinite pairs of *ℜ*_0_ and *τ* yield equivalent incidence dynamics. However, in virtue of their linear relationship, each value of *τ* corresponds to exactly one value of *ℜ*_0_.

### 3.3 Three unknowns (alternative):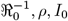, *ρ, I*_0_

More importantly, the linear relationships shown in Fig 4B reveal an intriguing insight. Notice that irrespective of the structure (*M* ^1*j*^) fitting data of any origin (*D*^1*j*^), the true values of *ℜ*_0_ and *τ* as a pair (the intersection between the dotted and dashed lines) are subsumed into any of the linear associations. This observation implies that the true *ℜ*_0_ can correspond only to the right *τ*. Therefore, it could be possible to accurately estimate *ℜ*_0_ from a model whose mean generation time is fixed to the true underlying value, but the shape of the epidemiological delays may differ from that of the data generating process. To test this hypothesis, we reformulate the *SE*^*i*^*I*^*j*^*R* framework so that *τ* becomes a parameter of every model instance. Consequently, we combine Eqs 2 and 5 into 6, which expresses *β* as a dependent variable of four parameters: *j, σ, τ*, and *ℜ*_0_.

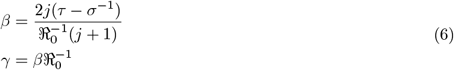

Parameter *j* is based on the fitting model’s structure, whereas *σ* and *τ* are fixed to the true values that produced the incidence reports. For instance, a *D*^12^ report stems from a structure whose *σ* and *τ* are equal to 0.5 and 3.5 (applying Eq 5), respectively. Therefore, an *M* ^14^ candidate fitting this report has *j, σ*, and *τ* fixed to 4, 0.5, and 3.5, respectively. An immediate consequence of this procedure is the need to constrain *γ* in order to maintain logical consistency. Accordingly, we define *γ* as a function of *β* and *ℜ*_0_ (6). This approach is analogous to fixing *γ* to an arbitrary value that yields the desired *τ*. Such a value may not correspond to that of the data generating process. Lastly, the remaining parameter, *ℜ*_0_, is subject to inference. We opt to estimate its inverse for a practical reason. Taking into account the threshold phenomenon and the fact that all incidence reports exhibit outbreak-like behaviour, any estimated value of *ℜ*_0_ must fall within the interval (1, *∞*). It then logically follows that its inverse 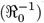 spans over the range (0, 1). This transformation permits the inference algorithm to operate in a much smaller parameter space, which enhances sampling efficiency.

We subsequently incorporate the redefined components (*β* and *γ*) into the *SE*^*i*^*I*^*j*^*R* framework to produce an alternative set of four candidate models with three unknowns: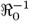, *ρ* and 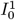. Similarly as before, we estimate the posterior distribution for each candidate fitted to an incidence report. The reader can find the complete set results in the electronic supplementary material *S2 §3*. These results once more highlight the intrinsic symmetry of SEIR formulations. Specifically, provided there is an adequate configuration, any candidate structure can accurately match the observed incidence despite differences in the infectious period distribution. Nevertheless, this alternative parameterisation exhibits a distinctive and crucial feature: the estimation of *ℜ*_0_ is less sensitive to the assumed distribution of the infectious delay. To support this claim, we present in Fig 5 the results of fitting the four alternative candidates to four incidence reports of dissimilar origin. Here, it can be seen that all candidates recover (via 95 % credible intervals) the underlying true *ℜ*_0_, notwithstanding the origin of the data or the fitting model.

**Figure 5:**
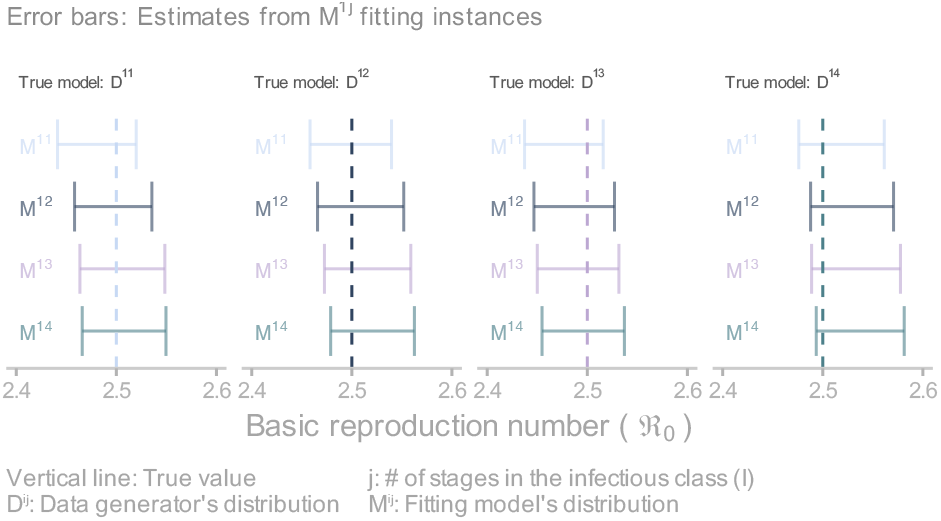
Inference results obtained from the three-unknown alternative parameterisation. This plot compares estimates for the basic reproduction number obtained from fitting four candidate models to four incidence reports. Error bars correspond to 95% credible intervals, and the vertical line denotes the true value.

Recovering the underlying *ℜ*_0_ is not exclusive to this sample of four datasets but is generalised across the *80* high-fidelity *D*^1*j*^ datasets. To summarise this insight, we borrow a concept from the *frequentist* tradition. Such a concept known as *coverage* [59] means that if one collects a large number of samples from the same process and constructs the corresponding confidence intervals, then a certain percentage of the intervals will contain or cover the true parameter. This percentage is given by the confidence level. For instance, if one fits a model to 100 datasets and estimates an equal number of confidence intervals at the 95% significance level, then 95 of those intervals will cover the true value. Admittedly, it is implicitly assumed that our 95% credible intervals (obtained from posterior distributions) are proportional to 95% confidence intervals. Indeed, estimated intervals for *ℜ*_0_ and *ρ* conform to this concept (see electronic supplementary material *S2 §3*.*3*), where minor deviations are justified by the fact that coverage is defined asymptotically (infinite measurements). However, asymptotics does not account for the large deviance observed in the estimates of *I*_0_. We explain this inconsistency in the section below where *I*_0_ becomes more prominent.

To conclude this section, we report the analysis of the low-fidelity datasets (right column in Fig 2). The reader can find the results in the electronic supplementary material *S3*. Overall, we obtain similar insights in comparison to those derived from the high-fidelity datasets. In the absence of structural differences, it is unsurprising that the effect of larger noise in the signal (overdispersion) results in greater uncertainty in parameter estimates. This decrease in precision (wider credible intervals) can obscure or accentuate features of the inference process. On the one hand, overdispersion masks biases in estimates. For instance, noisier measurements cause *I*_0_ estimates from the *alternative* parameterisation to conform to the expected coverage, which should not occur based on the results obtained from the high-fidelity datasets. On the other hand, overdispersion exacerbates identifiability issues. Under the *four-unknown* parameterisation, some *ℜ*_0_ estimates reach values up to 40. This result is a reminder that choosing an adequate number of unknowns is not a trivial decision. Setting more unknowns than the data can tolerate renders models unidentifiable. In this context, unidentifiability occurs because the incidence data does not provide enough information to update the prior distribution of *γ*. As discussed above, many values of *γ* are consistent with the observed incidence, an insight that holds for both levels of data fidelity. Finally, we note that overdispersion estimates are robust to the choice of the infectious period distribution.

### 3.4 Misspecifying the latent period distribution

Thus far, we have conducted the inference process assuming that the latent period distribution (*i*) is known. Lifting this constraint would strain our computational resources, producing a four-fold increase in the pool of candidates fitting a single report (assuming *i, j ∈ {*1, 2, 3, 4*}*). Instead of undertaking such costly exploration, one could leverage the fact that the mean generation time depends solely on the mean latent period rather than its particular distribution (Eq 5). To test this idea, we compare the estimates obtained from candidate models with the *right* and *wrong* latent period distribution. We illustrate this process with the *80 D*^3*j*^ low-fidelity (*ϕ*^*−*1^ = 1*/*3) datasets. For each dataset, we fit eight candidates *M* ^*ij*^ from the traditional three-unknown parameterisation, where the latent period distribution can take the wrong (*i* = 1) and the right (*i* = 3) values, and the infectious period distribution varies as before, namely, *j ∈ {*1, 2, 3, 4*}*. The reader can find the complete results in the electronic supplementary material *S5*.

To facilitate the presentation of the results, we first focus on candidates *M* ^13^ and *M* ^33^ fitting one *D*^33^ incidence report. Fig 6A shows that both models predict similar, although not identical, latent incidence dynamics. Further inspection reveals that the slight difference in the predicted incidence due to dissimilar latent period distributions does not lead to variation in *ℜ*_0_ estimates. To corroborate this assessment, we expand the analysis to the eight candidates matching the same incidence report. The right-hand side of Fig 6B shows that *ℜ*_0_ estimates are sensitive to variation in the structure of the infectious period but are indifferent to the latent period distribution. In compliance with the literature, the more dispersed latent period (*i* = 1) leads to an earlier incidence peak compared to the tighter distribution (*i* = 3) in the context of identical *ℜ*_0_.

**Figure 6:**
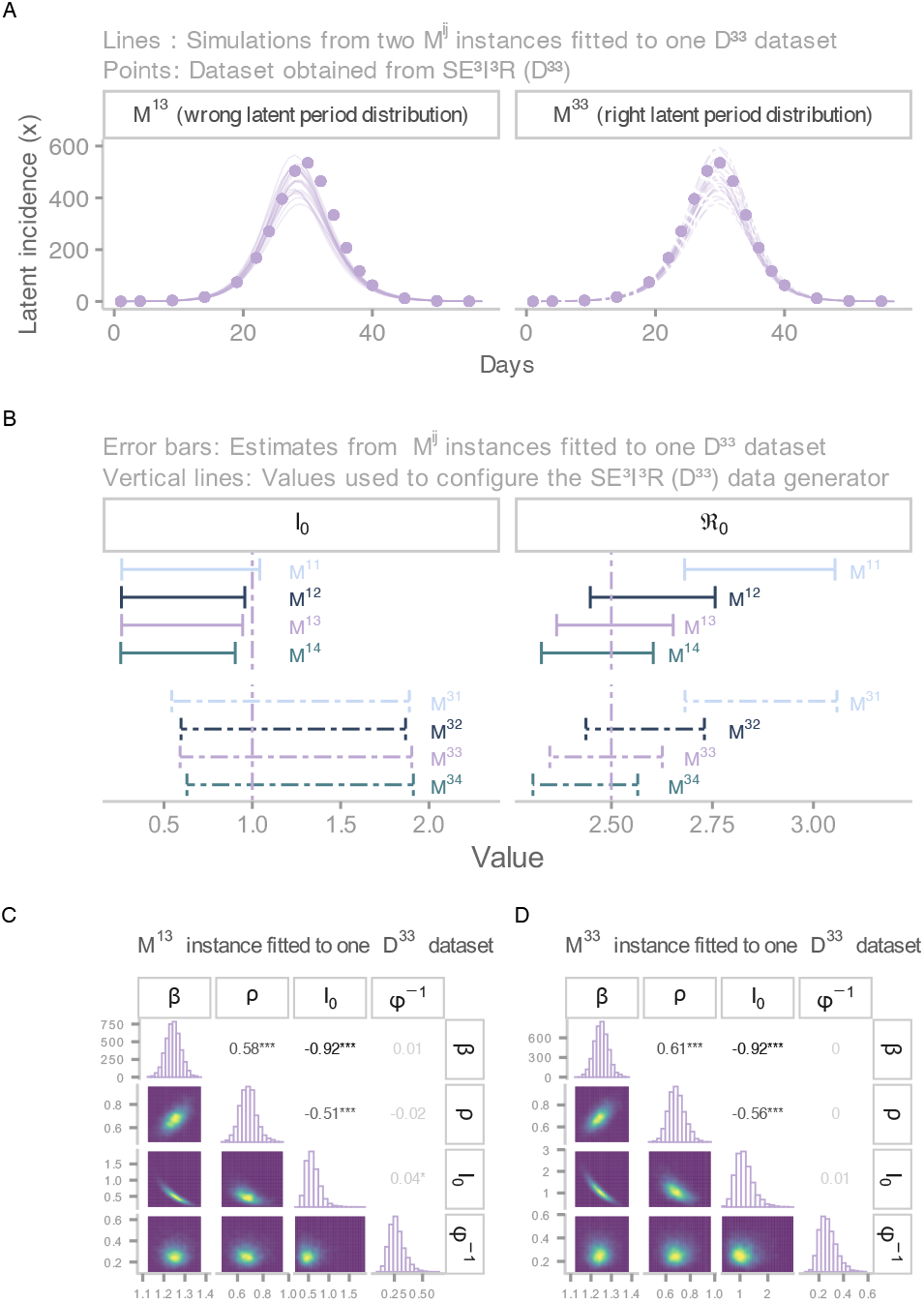
Comparing estimates from candidates models with the wrong and correct distributions of the latent period. These models stem from the three-unknown parameterisation A) Incidence fit from two candidate models matching a *D*^33^ incidence report. Candidate on the left has an exponientially-distributed (wrong) latent period, whereas candidate on the right has Gamma-distributed latent period (correct). B) Comparison of estimates of the initial number of infectious individuals at stage 1 and the basic reproduction number by fitting model. Error bars correspond to 95% credible intervals, and vertical lines denotes the true value. C) Joint posterior distribution from an *M* ^13^ candidate fitting a *D*^33^ report. The diagonal shows posterior marginal distributions. In the lower triangular part, each possible pairwise conditional distribution is displayed, whereas the upper triangular part presents the correlation among parameters. C) Joint posterior distribution from *M* ^33^ candidate fitting a *D*^33^ report.

Nevertheless, the mechanism that enables models with heterogeneous distributions to produce analogous incidence dynamics remains unexplained. The left-hand side of Fig 6B, which displays *I*_0_ estimates, provides the first hint. This plot shows that instances with the wrong latent period distribution (*M* ^1*j*^) systematically underestimate (via 95% credible intervals) the actual value (vertical line). To explain this phenomenon, we draw on a broader view of the posterior distribution. It is commonplace to restrict inference analyses to one parameter at a time (i.e. marginal distributions), neglecting the information provided by the full posterior distribution. To redress this shortcoming, we visualise the full distribution via pair plots. Specifically, Fig 6C corresponds to the summary of the posterior distribution obtained from fitting *M* ^13^ to one *D*^33^ incidence report. The upper triangular elements of this plot indicate that the three estimated parameters are strongly correlated. Especially *β* and *I*_0_, or more compellingly, *ℜ*_0_ and *I*_0_. Recall that the basic reproduction number is directly proportional to *β*. Therefore, although the mean generation time determines which *ℜ*_0_ corresponds to the observed incidence, *I*_0_ (and *ρ* to a lesser extent) regulates the flexibility of *ℜ*_0_ to reach such a desired value. Interestingly, *I*_0_ provides such a degree of flexibility that unrealistic adjustments in its estimates allow us to equate dissimilar model structures. Notice that the only discernible difference between Figs 6C and 6D (*M* ^33^ fitted to *D*^33^) is seen in the marginal distributions of *I*_0_. In fact, this phenomenon explains the *failure* of the alternative parameterisation to recover the true value of *I*_0_.

In view of these symmetries, it is not unreasonable to expect that candidates from the four-unknown and the alternative parameterisations, too, are indifferent to the latent period distribution once *I*_0_ and *ρ* correct for any misspecification. To verify this premise, we fit the parameterisations mentioned above to the *D*^3*j*^ low-fidelity incidence reports. As anticipated, the inference results indicate that the four-unknown parameterisation (S5 § 2) uncover the linear association between *τ* and *ρ* due to the unidentifiability of *γ*. Likewise, the alternative parameterisation (S5 § 3) recovers the true *ℜ*_0_ irrespective of the formulation of the epidemiological delays. Furthermore, we replicate these results using the *D*^3*j*^ high-fidelity datasets (see electronic supplementary information S4).

### 3.5 Sensitivity analysis

So far, model candidates have been amalgamated with the appropriate measurement component. In this section, we explore the implications that can arise from ignoring overdispersion. That is, equipping model candidates with a Poisson measurement component. We perform such exploration by inferring *ℜ*_0_ from *M* ^1*j*^ candidates (alternative parameterisation) fitted to the *D*^3*j*^ low-fidelity datasets discussed in the previous section. As expected, the results indicate that employing the Poisson distribution (see S5 § 4) leads to overconfident (too precise) and biased (inaccurate) estimates in the context of overdispersion. We observe these features with narrow uncertainty intervals that do not cover the true value. This result implies that the wrong choice of the measurement component can offset any gains in accuracy due to the alternative parameterisation.

On the other hand, the synthetic data used for the analysis presented in the previous sections stems from models configured to identical *ℜ*_0_ and similar mean generation times (variation due to the infectious period distribution). Naturally, one wonders whether the usefulness of the alternative parameterisation holds in other conditions. To answer this question, we repeat the workflow described in this paper for additional scenarios of *τ* and *ℜ*_0_. For simplicity, we restrict this sensitivity analysis to datasets derived from models with an exponentially-distributed latent period (*D*^1*j*^). Additionally, we equip the fitting candidates with the appropriate measurement component. The complete set of results is presented in the electronic supplementary information S6. We present these results in terms of scenarios (Table 1). For instance, the base case scenario, *Scenario 1*, corresponds to data generated from *SE*^*i*^*I*^*j*^*R* configured to *ℜ*_0_ = 2.5 and *τ*_*e*_ = 4 (results presented in Section 3.3), where *τ*_*e*_ serves as a scenario identifier and denotes the mean generation time obtained from an exponentially-distributed infectious period (*j* = 1).

**Table 1:** Scenarios

| Scenario | $\mathcal{R}_0$ | $\tau_e$ | $\mathcal{R}_0$ recovered? (High-fidelity) | $\mathcal{R}_0$ recovered? (Low-fidelity) |
| --- | --- | --- | --- | --- |
| 1 | 2.5 | 4 | Yes | Yes |
| 2 | 2.5 | 8 | Yes | Yes |
| 3 | 2.5 | 13 | Yes | Yes |
| 4 | 9.0 | 4 | No | Yes |
| 5 | 15.0 | 4 | No | Yes |

For *Scenario 2*, we increase the reference mean generation time (*τ*_*e*_ = 8), while keeping *ℜ*_0_ at 2.5. First, we focus on the high-fidelity datasets. Overall, the greater the divergence between the fitting model’s infectious period distribution and the distribution that generated the data, the greater the loss in accuracy. Namely, lower coverage. To provide an example, the 95% credible intervals constructed from *M* ^14^ candidates fitting *D*^11^ incidence reports only attain coverage of 30% for *ℜ*_0_. Closer inspection, though, reveals that such accuracy loss is more statistical than practical. To support this statement, we calculate the average relative difference between the actual and estimated *ℜ*_0_, finding that misspecification of the infectious period distribution leads to a maximum average relative error of 2%. In contrast, we would obtain discrepancies up to 15 % if we adopted the traditional approach. Simply put, it is costlier to misspecify the mean generation time than the mean infectious period. Furthermore, such slight differences in the alternative parameterisation are erased by overdispersion. That is, overdispersion masks minor misspecification in the process component. Moreover, in *Scenario 3* (*ℜ*_0_ = 2.5, *τ*_*e*_ = 13), we observe that further increasing of the mean generation time does not lead to significant drops in the coverage of *ℜ*_0_ under both levels of data fidelity. In a nutshell, it is reasonable to suggest that the alternative parameterisation is robust to various levels of the mean generation time.

Conversely, we cannot maintain the same assertion for various values of *ℜ*_0_. Indeed, Fig 4C provided the first hint. This plot shows that the straight lines do not overlap as *ℜ*_0_ reaches relatively high values. Consequently, in scenarios *4* (*ℜ*_0_ = 9, *τ*_*e*_ = 4) and *5* (*ℜ*_0_ = 17, *τ*_*e*_ = 4), we test the implications of larger transmissibility levels. The results indicate that as we increase the underlying *ℜ*_0_ for generating the data, the equivalency among fitting models dissipates and misspecification in the infectious period distribution leads to biased estimates of *ℜ*_0_. The size of such bias is proportional to the misspecification of the infectious period and the underlying *ℜ*_0_. This feature is primarily seen in the estimates derived from high-fidelity datasets, where coverage levels are low, and the average relative error between actual and estimated values cannot be overlooked. However, when we examine the posterior distributions obtained from fitting the low-fidelity data, it is seen that, once again, overdispersion masks misspecification in the process component, as evidenced by the high coverage levels. This is not to say that overdispersion is a desired feature in the data, but rather to emphasise that its presence hinders the attainment of precise estimates. Undoubtedly, having this understanding is of practical importance, given that it allows us to discern the necessary effort in data collection and model improvement.

## 4. Application to Influenza A

Leveraging the knowledge gained from the synthetic data, the last step in this work consists of exploiting the relationship between the basic reproduction number and the mean generation time to update the *ℜ*_0_ estimate of an outbreak of the 1918 influenza pandemic. The reader can find the full set of results in the electronic supplementary information S7. In particular, we focus on an outbreak that occurred in the city of Cumberland (Maryland) during the autumn of 1918, for which the U.S. Public Health Service organised special surveys [60] to determine the proportion of the population infected. Previous studies [12,22] employed the default heuristic of adopting an *SEIR* with exponentially-distributed epidemiological delays whose means were configured to values reported in the literature. Moreover, in these studies, the *SEIR* was coupled with the Poisson distribution resulting in a 95 % CI [2.5–2.6] for *ℜ*_0_. However, adopting a more realistic measurement component, such as the Negative Binomial distribution, produces lower and wider estimates: 95% CI [2.2, 2.4]. Further, if we jettison the assumption of an exponentially-distributed infectious period for a more realistic distribution, such as the gamma distribution, we even obtain lower estimates. For instance, a gamma-distributed infectious period with four stages (*SEI*^4^*R*) returns a 95% CI of [2.0, 2.2]. As noted earlier, the estimates obtained from this default heuristic or traditional approach are sensitive to the uncertainty in the infectious period distribution. On the contrary, when we fix the mean generation time in the *SEI*^*j*^*R* (alternative parameterisation) to a value (2.85 days) obtained from the literature [5,61], we derive nearly identical *ℜ*_0_ estimates (95% CI [2.0, 2.1]) regardless of the infectious period distribution (Fig 7). Notice that this estimate is similar to that obtained from the *SEI*^4^*R*, bolstering the fact that the actual infectious period is far from being exponentially distributed.

**Figure 7:**
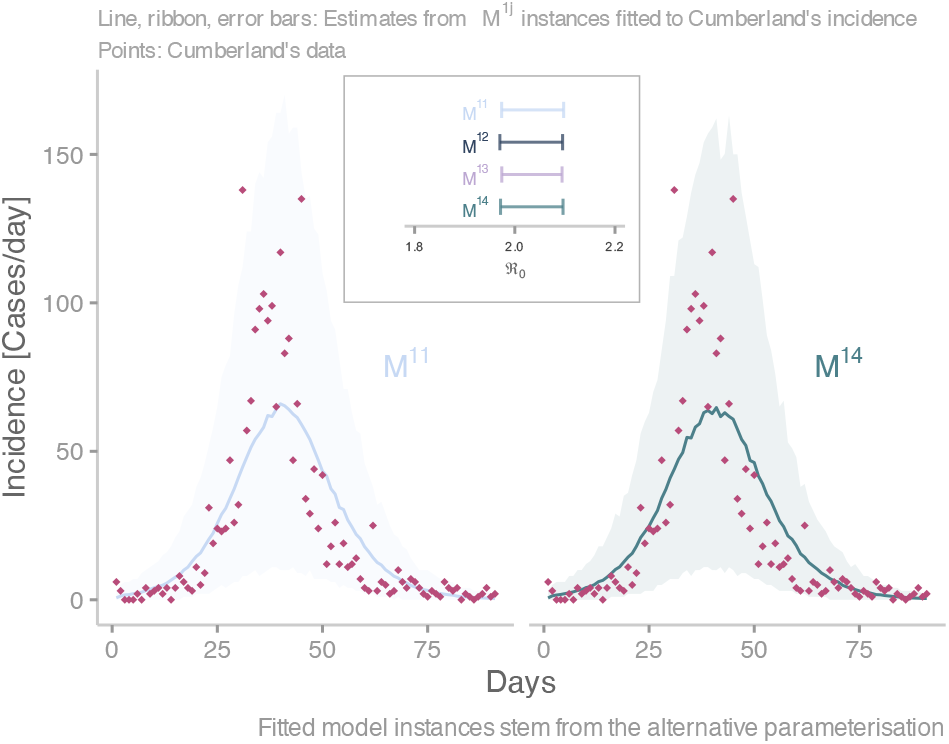
Application of the three-unknown alternative parameterisation. This plot shows the estimates obtained from fitting four candidates (*M* ^1*j*^) to the daily number of influenza cases (rhombi) detected by the U.S. Public Health Service in Cumberland (Maryland) during the 1918 influenza pandemic, from 22 September 1918 to 30 November 1918. Ribbons correspond to 95% credible intervals of the predicted measured incidence by two candidate models. The solid line denotes the median. The box inside the plot shows the estimates for the basic reproduction number by fitting model. Error bars correspond to 95% credible intervals.

## 5. Conclusion

The misspecification of various assumptions within the *SEIR* framework can negatively impact the estimation of *ℜ*_0_. In recognition of this risk, we ran a simulation study comprised of approximately 1000 synthetic datasets and 8000 model fits, whereby we identified the relative influence of some of those assumptions. Specifically, we found that fixing the mean generation time to a reliable estimate is of paramount importance. In contrast, one can be more lenient on the specification of the latent period distribution and the mean infectious period provided that other estimated parameters (*I*_0_ and *ρ*) redress the misspecification. We leveraged this knowledge to formulate an alternative parameterisation that is more robust to the uncertainty of the epidemiological delays. However, there is a caveat with this alternative formulation. Although it exploits a local symmetry (incidence dynamics) of the *SEIR* framework, such symmetry does not extend to the other states of the system. Therefore, the usefulness of the alternative parameterisation is confined to the estimation of *ℜ*_0_, and it is not a substitute for other kinds of analyses. For instance, if, on the contrary, our variable of interest were *I*_0_, we would obtain unreliable estimates. Furthermore, the alternative *SEIR* with exponentially-distributed delays will be as overoptimistic as its traditional counterpart in predicting the critical vaccination proportion or the effectiveness of an imperfect VIH treatment in the context of within-host dynamics [56]. Therefore, the alternative parameterisation is a mitigation strategy in the absence of complete information. Furthermore, its usefulness is abated by highly transmissible pathogens (Section 3.5). Nevertheless, biases in the estimates due to large *ℜ*_0_ are only detected with high-fidelity data. That is, data with little or no overdispersion.

Despite the significant computational effort of simulation analyses, a single study cannot offer overarching statements. Further work is required to test the validity of these insights in stricter or more elaborated contexts. For instance, we assumed the complete availability of the incidence time-series throughout this study. This assumption restricts the validity of the approach to retrospective analyses. However, other situations exist where only fewer incidence measurements are available to the modellers, such as the early phase of a pandemic response. Hence, it remains to be seen the effect of various levels of data availability on the performance of the suggested approach. Furthermore, for simplicity, we ignored age-related effects in the dynamics of the infectious disease as well as process stochasticity (demographic and environmental) and time-varying contact rates. We expect that future research builds on the findings provided by this study and addresses the aforementioned challenges to construct ever more reliable inference approaches.

## Acknowledgement

The project has received funding from the European Union’s Horizon 2020 research and innovation programme under grant agreement No 883285. The material presented and views expressed here are the responsibility of the author(s) only. The EU Commission takes no responsibility for any use made of the information set out. The funders had no role in study design, data collection and analysis, decision to publish, or preparation of the manuscript.

