## Supplementary material for "Anchoring the mean generation time in the SEIR to mitigate biases in ℜ_0_ estimates due to uncertainty in the distribution of the epidemiological delays": S1.html


# S1

This electronic supplementary material aims to document in a
reproducible manner the generation of the *instruments* employed
to carry out the inference study presented in the main text. By
instruments, we refer to the synthetic incidence reports and the models
used to fit them. Both instruments are the product of a common data
generating process.

### 1 Synthetic incidence

#### 1.1 System (latent) component

The first step for generating synthetic data consists of simulating
each of the \(SE^iI^jR\) instances. As
mentioned in the main text, we restrict the number of instances to
eight. Two for the latent period, \(i
=\{1,3\}\), and four, \(j
=\{1,2,3,4\}\), for the infectious period. Moreover, these
instances are configured to plausible parameter values and initial
conditions.

##### 1.1.1 Parameter values

Irrespective of the distribution of the latent or the infectious
period, all of the instances are configured to the values below. Notice
that the choice of these values implies that all instances share an
identical \(\Re\_0\) value (2.5).

Table 1. Constants

| Parameter | Value |
| --- | --- |
| $$\beta$$ | 1.25 |
| $$\sigma$$ | 0.50 |
| $$\gamma$$ | 0.50 |
| $$\rho$$ | 0.75 |
| $$N$$ | 10000.00 |

##### 1.1.2 Initial conditions

Regarding the initial conditions of the eight instances, all states
set to zero except for the number of susceptible individuals (\(S\)) and infectious individuals in the
first stage (\(I^1\)).

Table 2. Inits

| State | Init value |
| --- | --- |
| $$S$$ | 9999 |
| $$I^1$$ | 1 |

##### 1.1.3 Latent incidence

By latent incidence, we mean the *smooth* incidence predicted
by an ODE model. The plot below shows the latent incidence predicted by
the eight instances of the \(SE^iI^jR\)
framework.

Fig 1. Incidence obtained from the ODE instances

#### 1.2 Observational component

Subsequently, for each latent incidence, we generate \(40\) incidence reports using the Negative
Binomial distribution. Specifically, we produce *two* sets of 20
reports. The first set’s noise level is equivalent to that of the
Poisson distribution (no overdispersion). Namely, the concentration
parameter (\(\phi^{-1}\)) is set to
zero. We refer to these sets as *high-fidelity*. For the other
sets, we add overdispersion (\(\phi^{-1} =
1/3\)) and identified them as *low-fidelity*.

##### 1.2.1 High-fidelity (Poisson) \(D^{1j}\)

Fig 2. Simulated incidence reports. Measurement noise from the Poisson
distribution was added to the smooth trajectories obtained from SEIR
instances with an exponential-distributed latent period.

##### 1.2.2 Low-fidelity (overdispersed) \(D^{1j}\)

Fig 3. Simulated incidence reports. Measurement noise from the negative
binomial distribution with overdispersion was added to the smooth
trajectories obtained from SEIR instances with an
exponential-distributed latent period.

##### 1.2.3 High-fidelity \(D^{3j}\)

Fig 4. Simulated incidence reports. Measurement noise from the Poisson
distribution was added to the smooth trajectories obtained from SEIR
instances with an gamma-distributed latent period.

##### 1.2.4 Low-fidelity \(D^{3j}\)

Fig 5. Simulated incidence reports. Measurement noise from the negative
binomial distribution with overdispersion was added to the smooth
trajectories obtained from SEIR instances with an gamma-distributed
latent period.

### 2 Inference files

One can employ mechanistic models as inference tools to fit data in
order to estimate unknown quantities. In this work, we perform the
inference process through the Statistical software *Stan*. This
software requires users to specify the instructions for the sampling
process in Stan’s own language. In the sections below, we show examples
of such language for the three parameterisations described in the main
text.

#### 2.1 Three-unknown parameterisation (traditional)

##### 2.1.1 Example 1

Here is an example of an \(M^{11}\)
candidate coupled with a Poisson measurement model.

```
## functions {
##   vector X_model(real time, vector y, array[] real params) {
##     vector[5] dydt;
##     real S_to_E;
##     real E1_to_I1;
##     real I_to_R;
##     real C_in;
##     S_to_E = params[1]*y[1]*y[3]/params[3];
##     E1_to_I1 = params[4]*y[2];
##     I_to_R = 1*params[5]*y[3];
##     C_in = params[2]*E1_to_I1;
##     dydt[1] = -S_to_E;
##     dydt[2] = S_to_E-E1_to_I1;
##     dydt[3] = E1_to_I1-I_to_R;
##     dydt[4] = I_to_R;
##     dydt[5] = C_in;
##     return dydt;
##   }
## }
## data {
##   int<lower = 1> n_obs;
##   int<lower = 1> n_params;
##   int<lower = 1> n_difeq;
##   array[n_obs] int y;
##   real t0;
##   array[n_obs] real ts;
##   real N;
##   real par_sigma;
##   real par_gamma;
##   real xi;
## }
## parameters {
##   real<lower = 0> par_beta;
##   real<lower = 0, upper = 1> par_rho;
##   real<lower = 0> I0;
## }
## transformed parameters{
##   array[n_obs] vector[n_difeq] x; // Output from the ODE solver
##   array[n_params] real params;
##   vector[n_difeq] x0; // init values
##   array[n_obs] real delta_x_1;
##   x0[1] = N * (1 -  xi) - I0; // S
##   x0[2] = 0; // E1
##   x0[3] = I0; // I1
##   x0[4] = xi * N; // R
##   x0[5] = I0; // C
##   params[1] = par_beta;
##   params[2] = par_rho;
##   params[3] = N;
##   params[4] = par_sigma;
##   params[5] = par_gamma;
##   x = ode_rk45(X_model, x0, t0, ts, params);
##   delta_x_1[1] =  x[1, 5] - x0[5] + 1e-5;
##   for (i in 1:n_obs-1) {
##     delta_x_1[i + 1] = x[i + 1, 5] - x[i, 5] + 1e-5;
##   }
## }
## model {
##   par_beta ~ lognormal(0, 1);
##   par_rho ~ beta(2, 2);
##   I0 ~ lognormal(0, 1);
##   y ~ poisson(delta_x_1);
## }
## generated quantities {
##   real log_lik;
##   array[n_obs] int sim_y;
##   log_lik = poisson_lpmf(y | delta_x_1);
##   sim_y = poisson_rng(delta_x_1);
## }
```

##### 2.1.2 Example 2

Here is an example of an \(M^{32}\)
candidate coupled with a Negative Binomial measurement model.

```
## functions {
##   vector X_model(real time, vector y, array[] real params) {
##     vector[9] dydt;
##     real S_to_E;
##     real E1_to_E2;
##     real I1_to_I2;
##     real I2_to_I3;
##     real I3_to_R;
##     real E2_to_E3;
##     real E3_to_I1;
##     real C_in;
##     S_to_E = params[1]*y[1]*(y[3]+y[6]+y[7])/params[3];
##     E1_to_E2 = 3*params[4]*y[2];
##     I1_to_I2 = 3*params[5]*y[3];
##     I2_to_I3 = 3*params[5]*y[6];
##     I3_to_R = 3*params[5]*y[7];
##     E2_to_E3 = 3*params[4]*y[8];
##     E3_to_I1 = 3*params[4]*y[9];
##     C_in = params[2]*E3_to_I1;
##     dydt[1] = -S_to_E;
##     dydt[2] = S_to_E-E1_to_E2;
##     dydt[3] = E3_to_I1-I1_to_I2;
##     dydt[4] = I3_to_R;
##     dydt[5] = C_in;
##     dydt[6] = I1_to_I2-I2_to_I3;
##     dydt[7] = I2_to_I3-I3_to_R;
##     dydt[8] = E1_to_E2-E2_to_E3;
##     dydt[9] = E2_to_E3-E3_to_I1;
##     return dydt;
##   }
## }
## data {
##   int<lower = 1> n_obs;
##   int<lower = 1> n_params;
##   int<lower = 1> n_difeq;
##   array[n_obs] int y;
##   real t0;
##   array[n_obs] real ts;
##   real N;
##   real par_sigma;
##   real par_gamma;
##   real xi;
## }
## parameters {
##   real<lower = 0> par_beta;
##   real<lower = 0, upper = 1> par_rho;
##   real<lower = 0> I0;
##   real<lower = 0> inv_phi;
## }
## transformed parameters{
##   array[n_obs] vector[n_difeq] x; // Output from the ODE solver
##   array[n_params] real params;
##   vector[n_difeq] x0; // init values
##   array[n_obs] real delta_x_1;
##   real phi;
##   phi = 1 / inv_phi;
##   x0[1] = N * (1 -  xi) - I0; // S
##   x0[2] = 0; // E1
##   x0[3] = I0; // I1
##   x0[4] = xi * N; // R
##   x0[5] = I0; // C
##   x0[6] = 0; // I2
##   x0[7] = 0; // I3
##   x0[8] = 0; // E2
##   x0[9] = 0; // E3
##   params[1] = par_beta;
##   params[2] = par_rho;
##   params[3] = N;
##   params[4] = par_sigma;
##   params[5] = par_gamma;
##   x = ode_rk45(X_model, x0, t0, ts, params);
##   delta_x_1[1] =  x[1, 5] - x0[5] + 1e-5;
##   for (i in 1:n_obs-1) {
##     delta_x_1[i + 1] = x[i + 1, 5] - x[i, 5] + 1e-5;
##   }
## }
## model {
##   par_beta ~ lognormal(0, 1);
##   par_rho ~ beta(2, 2);
##   I0 ~ lognormal(0, 1);
##   inv_phi ~ exponential(5);
##   y ~ neg_binomial_2(delta_x_1, phi);
## }
## generated quantities {
##   real log_lik;
##   array[n_obs] int sim_y;
##   log_lik = neg_binomial_2_lpmf(y | delta_x_1, phi);
##   sim_y = neg_binomial_2_rng(delta_x_1, phi);
## }
```

#### 2.2 Four-unknown parameterisation

##### 2.2.1 Example 1

Here is an example of an \(M^{14}\)
candidate coupled with a Poisson measurement model.

```
## functions {
##   vector X_model(real time, vector y, array[] real params) {
##     vector[8] dydt;
##     real S_to_E;
##     real E1_to_I1;
##     real I1_to_I2;
##     real C_in;
##     real I2_to_I3;
##     real I3_to_I4;
##     real I4_to_R;
##     S_to_E = params[1]*y[1]*(y[3]+y[6]+y[7]+y[8])/params[4];
##     E1_to_I1 = params[5]*y[2];
##     I1_to_I2 = 4*params[3]*y[3];
##     C_in = params[2]*E1_to_I1;
##     I2_to_I3 = 4*params[3]*y[6];
##     I3_to_I4 = 4*params[3]*y[7];
##     I4_to_R = 4*params[3]*y[8];
##     dydt[1] = -S_to_E;
##     dydt[2] = S_to_E-E1_to_I1;
##     dydt[3] = E1_to_I1-I1_to_I2;
##     dydt[4] = I4_to_R;
##     dydt[5] = C_in;
##     dydt[6] = I1_to_I2-I2_to_I3;
##     dydt[7] = I2_to_I3-I3_to_I4;
##     dydt[8] = I3_to_I4-I4_to_R;
##     return dydt;
##   }
## }
## data {
##   int<lower = 1> n_obs;
##   int<lower = 1> n_params;
##   int<lower = 1> n_difeq;
##   array[n_obs] int y;
##   real t0;
##   array[n_obs] real ts;
##   real N;
##   real par_sigma;
##   real xi;
## }
## parameters {
##   real<lower = 0> par_beta;
##   real<lower = 0, upper = 1> par_rho;
##   real<lower = 0> I0;
##   real<lower = 0, upper = 1> par_gamma;
## }
## transformed parameters{
##   array[n_obs] vector[n_difeq] x; // Output from the ODE solver
##   array[n_params] real params;
##   vector[n_difeq] x0; // init values
##   array[n_obs] real delta_x_1;
##   x0[1] = N * (1 -  xi) - I0; // S
##   x0[2] = 0; // E1
##   x0[3] = I0; // I1
##   x0[4] = xi * N; // R
##   x0[5] = I0; // C
##   x0[6] = 0; // I2
##   x0[7] = 0; // I3
##   x0[8] = 0; // I4
##   params[1] = par_beta;
##   params[2] = par_rho;
##   params[3] = par_gamma;
##   params[4] = N;
##   params[5] = par_sigma;
##   x = ode_rk45(X_model, x0, t0, ts, params);
##   delta_x_1[1] =  x[1, 5] - x0[5] + 1e-5;
##   for (i in 1:n_obs-1) {
##     delta_x_1[i + 1] = x[i + 1, 5] - x[i, 5] + 1e-5;
##   }
## }
## model {
##   par_beta ~ lognormal(0, 1);
##   par_rho ~ beta(2, 2);
##   I0 ~ lognormal(0, 1);
##   par_gamma ~ beta(2, 2);
##   y ~ poisson(delta_x_1);
## }
## generated quantities {
##   real log_lik;
##   array[n_obs] int sim_y;
##   log_lik = poisson_lpmf(y | delta_x_1);
##   sim_y = poisson_rng(delta_x_1);
## }
```

##### 2.2.2 Example 2

Here is an example of an \(M^{13}\)
candidate coupled with a Negative Binomial measurement model.

```
## functions {
##   vector X_model(real time, vector y, array[] real params) {
##     vector[7] dydt;
##     real S_to_E;
##     real E1_to_I1;
##     real I1_to_I2;
##     real C_in;
##     real I2_to_I3;
##     real I3_to_R;
##     S_to_E = params[1]*y[1]*(y[3]+y[6]+y[7])/params[4];
##     E1_to_I1 = params[5]*y[2];
##     I1_to_I2 = 3*params[3]*y[3];
##     C_in = params[2]*E1_to_I1;
##     I2_to_I3 = 3*params[3]*y[6];
##     I3_to_R = 3*params[3]*y[7];
##     dydt[1] = -S_to_E;
##     dydt[2] = S_to_E-E1_to_I1;
##     dydt[3] = E1_to_I1-I1_to_I2;
##     dydt[4] = I3_to_R;
##     dydt[5] = C_in;
##     dydt[6] = I1_to_I2-I2_to_I3;
##     dydt[7] = I2_to_I3-I3_to_R;
##     return dydt;
##   }
## }
## data {
##   int<lower = 1> n_obs;
##   int<lower = 1> n_params;
##   int<lower = 1> n_difeq;
##   array[n_obs] int y;
##   real t0;
##   array[n_obs] real ts;
##   real N;
##   real par_sigma;
##   real xi;
## }
## parameters {
##   real<lower = 0> par_beta;
##   real<lower = 0, upper = 1> par_rho;
##   real<lower = 0> I0;
##   real<lower = 0, upper = 1> par_gamma;
##   real<lower = 0> inv_phi;
## }
## transformed parameters{
##   array[n_obs] vector[n_difeq] x; // Output from the ODE solver
##   array[n_params] real params;
##   vector[n_difeq] x0; // init values
##   array[n_obs] real delta_x_1;
##   real phi;
##   phi = 1 / inv_phi;
##   x0[1] = N * (1 -  xi) - I0; // S
##   x0[2] = 0; // E1
##   x0[3] = I0; // I1
##   x0[4] = xi * N; // R
##   x0[5] = I0; // C
##   x0[6] = 0; // I2
##   x0[7] = 0; // I3
##   params[1] = par_beta;
##   params[2] = par_rho;
##   params[3] = par_gamma;
##   params[4] = N;
##   params[5] = par_sigma;
##   x = ode_rk45(X_model, x0, t0, ts, params);
##   delta_x_1[1] =  x[1, 5] - x0[5] + 1e-5;
##   for (i in 1:n_obs-1) {
##     delta_x_1[i + 1] = x[i + 1, 5] - x[i, 5] + 1e-5;
##   }
## }
## model {
##   par_beta ~ lognormal(0, 1);
##   par_rho ~ beta(2, 2);
##   I0 ~ lognormal(0, 1);
##   par_gamma ~ beta(2, 2);
##   inv_phi ~ exponential(5);
##   y ~ neg_binomial_2(delta_x_1, phi);
## }
## generated quantities {
##   real log_lik;
##   array[n_obs] int sim_y;
##   log_lik = neg_binomial_2_lpmf(y | delta_x_1, phi);
##   sim_y = neg_binomial_2_rng(delta_x_1, phi);
## }
```

#### 2.3 Three-unknown parameterisation (Alternative)

##### 2.3.1 Example 1

Here is an example of an \(M^{11}\)
candidate coupled with a Poisson measurement model.

```
## functions {
##   vector X_model(real time, vector y, array[] real params) {
##     vector[5] dydt;
##     real E1_to_I1;
##     real C_in;
##     real aux_j;
##     real aux_tau;
##     real var_beta;
##     real var_gamma;
##     real S_to_E;
##     real I_to_R;
##     E1_to_I1 = params[5]*y[2];
##     C_in = params[2]*E1_to_I1;
##     aux_j = (1+1)/(2.0*1);
##     aux_tau = params[4]-(1/params[5]);
##     var_beta = (1/params[1])*(aux_j/aux_tau);
##     var_gamma = var_beta*params[1];
##     S_to_E = var_beta*y[1]*y[3]/params[3];
##     I_to_R = 1*var_gamma*y[3];
##     dydt[1] = -S_to_E;
##     dydt[2] = S_to_E-E1_to_I1;
##     dydt[3] = E1_to_I1-I_to_R;
##     dydt[4] = I_to_R;
##     dydt[5] = C_in;
##     return dydt;
##   }
## }
## data {
##   int<lower = 1> n_obs;
##   int<lower = 1> n_params;
##   int<lower = 1> n_difeq;
##   array[n_obs] int y;
##   real t0;
##   array[n_obs] real ts;
##   real N;
##   real par_tau;
##   real par_sigma;
##   real xi;
## }
## parameters {
##   real<lower = 0, upper = 1> par_inv_R0;
##   real<lower = 0, upper = 1> par_rho;
##   real<lower = 0> I0;
## }
## transformed parameters{
##   array[n_obs] vector[n_difeq] x; // Output from the ODE solver
##   array[n_params] real params;
##   vector[n_difeq] x0; // init values
##   array[n_obs] real delta_x_1;
##   x0[1] = N * (1 -  xi) - I0; // S
##   x0[2] = 0; // E1
##   x0[3] = I0; // I1
##   x0[4] = xi * N; // R
##   x0[5] = I0; // C
##   params[1] = par_inv_R0;
##   params[2] = par_rho;
##   params[3] = N;
##   params[4] = par_tau;
##   params[5] = par_sigma;
##   x = ode_rk45(X_model, x0, t0, ts, params);
##   delta_x_1[1] =  x[1, 5] - x0[5] + 1e-5;
##   for (i in 1:n_obs-1) {
##     delta_x_1[i + 1] = x[i + 1, 5] - x[i, 5] + 1e-5;
##   }
## }
## model {
##   par_inv_R0 ~ beta(2, 2);
##   par_rho ~ beta(2, 2);
##   I0 ~ lognormal(0, 1);
##   y ~ poisson(delta_x_1);
## }
## generated quantities {
##   real log_lik;
##   array[n_obs] int sim_y;
##   log_lik = poisson_lpmf(y | delta_x_1);
##   sim_y = poisson_rng(delta_x_1);
## }
```

##### 2.3.2 Example 2

Here is an example of an \(M^{12}\)
candidate coupled with a Negative Binomial measurement model.

```
## functions {
##   vector X_model(real time, vector y, array[] real params) {
##     vector[6] dydt;
##     real E1_to_I1;
##     real C_in;
##     real aux_j;
##     real aux_tau;
##     real var_beta;
##     real var_gamma;
##     real I2_to_R;
##     real S_to_E;
##     real I1_to_I2;
##     E1_to_I1 = params[5]*y[2];
##     C_in = params[2]*E1_to_I1;
##     aux_j = (2+1)/(2.0*2);
##     aux_tau = params[4]-(1/params[5]);
##     var_beta = (1/params[1])*(aux_j/aux_tau);
##     var_gamma = var_beta*params[1];
##     I2_to_R = 2*var_gamma*y[6];
##     S_to_E = var_beta*y[1]*(y[3]+y[6])/params[3];
##     I1_to_I2 = 2*var_gamma*y[3];
##     dydt[1] = -S_to_E;
##     dydt[2] = S_to_E-E1_to_I1;
##     dydt[3] = E1_to_I1-I1_to_I2;
##     dydt[4] = I2_to_R;
##     dydt[5] = C_in;
##     dydt[6] = I1_to_I2-I2_to_R;
##     return dydt;
##   }
## }
## data {
##   int<lower = 1> n_obs;
##   int<lower = 1> n_params;
##   int<lower = 1> n_difeq;
##   array[n_obs] int y;
##   real t0;
##   array[n_obs] real ts;
##   real N;
##   real par_tau;
##   real par_sigma;
##   real xi;
## }
## parameters {
##   real<lower = 0, upper = 1> par_inv_R0;
##   real<lower = 0, upper = 1> par_rho;
##   real<lower = 0> I0;
##   real<lower = 0> inv_phi;
## }
## transformed parameters{
##   array[n_obs] vector[n_difeq] x; // Output from the ODE solver
##   array[n_params] real params;
##   vector[n_difeq] x0; // init values
##   array[n_obs] real delta_x_1;
##   real phi;
##   phi = 1 / inv_phi;
##   x0[1] = N * (1 -  xi) - I0; // S
##   x0[2] = 0; // E1
##   x0[3] = I0; // I1
##   x0[4] = xi * N; // R
##   x0[5] = I0; // C
##   x0[6] = 0; // I2
##   params[1] = par_inv_R0;
##   params[2] = par_rho;
##   params[3] = N;
##   params[4] = par_tau;
##   params[5] = par_sigma;
##   x = ode_rk45(X_model, x0, t0, ts, params);
##   delta_x_1[1] =  x[1, 5] - x0[5] + 1e-5;
##   for (i in 1:n_obs-1) {
##     delta_x_1[i + 1] = x[i + 1, 5] - x[i, 5] + 1e-5;
##   }
## }
## model {
##   par_inv_R0 ~ beta(2, 2);
##   par_rho ~ beta(2, 2);
##   I0 ~ lognormal(0, 1);
##   inv_phi ~ exponential(5);
##   y ~ neg_binomial_2(delta_x_1, phi);
## }
## generated quantities {
##   real log_lik;
##   array[n_obs] int sim_y;
##   log_lik = neg_binomial_2_lpmf(y | delta_x_1, phi);
##   sim_y = neg_binomial_2_rng(delta_x_1, phi);
## }
```
