## Supplementary material for "Anchoring the mean generation time in the SEIR to mitigate biases in ℜ_0_ estimates due to uncertainty in the distribution of the epidemiological delays": S2.html


# S2

This appendix illustrates the process of fitting various
parameterisations of the \(SE^1I^jR\)
model (\(M^{1j}\)) to
**high-fidelity \(D^{1j}\)** incidence reports. We
mean by *parameterisation* the decision of categorising model
parameters as either *unknown* or *assumed*. For the
unknown parameters, we construct prior distributions, which will be
eventually updated in light of the data via HMC sampling, resulting in a
posterior distribution. On the other hand, assumed parameters are fixed
at their true values.

### 1 Three-unknown parameterisation (traditional)

For candidates from this parameterisation, we assume that two
parameters and one initial condition are unknown: \(\beta\), \(\rho\), and \(I^1\_0\). Each candidate is coupled with the
appropriate likelihood function. That is, the *Poisson*
distribution.

#### 1.1 Prior distributions

The prior distributions shown below apply for all candidates from
this parameterisation.

##### 1.1.1 Effective contact rate (\(\beta\))

Fig 1. Histogram from samples obtained from the contact rate’s prior
distribution

##### 1.1.2 Reporting rate (\(\rho\))

Fig 2. Histogram from samples obtained from the reporting rate’s prior
distribution

##### 1.1.3 Initial number of infectious individuals (\(I\_0\))

Fig 3. Histogram from samples obtained from the prior distribution of
the initial number of infectious individuals.

#### 1.2 Posterior distributions

We stratify inference results by the infectious period distribution
that produced the observed incidences. For instance, \(D\_{11}\) implies that candidate models
fitted incidence reports that stem from the \(SE^1I^1R\).

##### 1.2.1 Fitting \(D^{11}\)

###### 1.2.1.1 Incidence fit

The figure below compares actual (points) and simulated (lines)
latent incidence by model candidate.

Fig 4. Posterior predictive checks against latent incidence. Dots denote
synthetic data while lines indicate simulations from candidates’ ODE
structure. We configure these structures using samples from the
posterior distribution.

To persuade the reader that all models fit the data equally well, we
draw on the mean absolute scaled error (MASE). This quantity is a
measure of the accuracy of forecasts, well-suited for time-series.
Therefore, we employ the MASE to compare each of the 4000 simulated
latent incidences against its true counterpart (\(x\)) and observed incidence (\(y\)). By simulated latent incidences, we
refer to the process of plugging the samples of a posterior distribution
into an ODE model to obtain incidence trajectories via simulation. We
summarise the results via histograms in the plot below. The left column
(of panels) contains the comparison between simulated and actual latent
incidences. The right column of panels displays the comparison between
simulated latent incidence and the observed incidence. Overall, there is
no noticeable variation in the histograms as the fitting model changes
(increasing \(j\)).

Fig 5. Fit scores by candidate model and type of data. \(x\) denotes latent incidence, whereas \(y\) indicates observed incidence. Vertical
line denotes the mean.

###### 1.2.1.2 Joint posterior distribution

In this plot, we show an example of a joint posterior distribution.
Specifically, this posterior distribution was derived from fitting the
\(M^{11}\) candidate to one \(D^{11}\) incidence report.

Fig 6. Posterior distribution. The diagonal shows the posterior marginal
distributions (histograms). In the lower triangular part, the joint
posterior distribution of each possible combination of two parameters is
displayed (heatmaps). The upper triangular part shows the correlation
among parameters.

###### 1.2.1.3 Marginal posterior distributions

In this section, we present a comparison between estimated marginal
posterior distributions (error bars) and actual values (vertical lines).
Such distributions are the result of fitting all four candidates to four
\(D^{11}\) incidence reports. The value
in the middle of the error bars indicates the relative error between the
actual value and the mean of the marginal posterior distribution.

###### 1.2.1.3.1 Basic reproduction number (\(\Re\_0\))

Fig 7. Estimates of the basic reproduction number by dataset and model
candidate. Error bars denote 95% credible intervals. The value in the
middle of the bars indicates the distance between the marginal
posterior’s mean and the true value (vertical line).

###### 1.2.1.3.2 Initial number of infectious individuals (\(I\_0\))

Fig 8. Estimates of the initial number of infectious individuals by
dataset and model candidate. Error bars denote 95% credible intervals.
The value in the middle of the bars indicates the distance between the
marginal posterior’s mean and the true value (vertical line).

###### 1.2.1.3.3 Reporting rate (\(\rho\))

Fig 9. Estimates of the reporting rate by dataset and model candidate.
Error bars denote 95% credible intervals. The value in the middle of the
bars indicates the distance between the marginal posterior’s mean and
the true value (vertical line).

##### 1.2.2 Fitting \(D^{12}\)

###### 1.2.2.1 Incidence fit

Fig 10. Posterior predictive checks against latent incidence. Dots
denote synthetic data while lines indicate simulations from candidates’
ODE structure. We configure these structures using samples from the
posterior distribution.

Fig 11. Fit scores by candidate model and type of data. \(x\) denotes latent incidence, whereas \(y\) indicates observed incidence. Vertical
line denotes the mean.

###### 1.2.2.2 Joint posterior distribution

In this plot, we show an example of a joint posterior distribution.
Specifically, this posterior distribution was derived from fitting the
\(M^{12}\) candidate to one \(D^{12}\) incidence report.

Fig 12. Posterior distribution. The diagonal shows the posterior
marginal distributions (histograms). In the lower triangular part, the
joint posterior distribution of each possible combination of two
parameters is displayed (heatmaps). The upper triangular part shows the
correlation among parameters.

###### 1.2.2.3 Marginal posterior distributions

###### 1.2.2.3.1 Basic reproduction number (\(\Re\_0\))

Fig 13. Estimates of the basic reproduction number by dataset and model
candidate. Error bars denote 95% credible intervals. The value in the
middle of the bars indicates the distance between the marginal
posterior’s mean and the true value (vertical line).

###### 1.2.2.3.2 Initial number of infectious individuals (\(I\_0\))

Fig 14. Estimates of the initial number of infectious individuals by
dataset and model candidate. Error bars denote 95% credible intervals.
The value in the middle of the bars indicates the distance between the
marginal posterior’s mean and the true value (vertical line).

###### 1.2.2.3.3 Reporting rate (\(\rho\))

Fig 15. Estimates of the reporting rate by dataset and model candidate.
Error bars denote 95% credible intervals. The value in the middle of the
bars indicates the distance between the marginal posterior’s mean and
the true value (vertical line).

##### 1.2.3 Fitting \(D^{13}\)

###### 1.2.3.1 Incidence fit

Fig 16. Posterior predictive checks against latent incidence. Dots
denote synthetic data while lines indicate simulations from candidates’
ODE structure. We configure these structures using samples from the
posterior distribution.

Fig 17. Fit scores by candidate model and type of data. \(x\) denotes latent incidence, whereas \(y\) indicates observed incidence. Vertical
line denotes the mean.

###### 1.2.3.2 Joint posterior distribution

In this plot, we show an example of a joint posterior distribution.
Specifically, this posterior distribution was derived from fitting the
\(M^{13}\) candidate to one \(D^{13}\) incidence report.

Fig 18. Posterior distribution. The diagonal shows the posterior
marginal distributions (histograms). In the lower triangular part, the
joint posterior distribution of each possible combination of two
parameters is displayed (heatmaps). The upper triangular part shows the
correlation among parameters.

###### 1.2.3.3 Marginal posterior distributions

###### 1.2.3.3.1 Basic reproduction number (\(\Re\_0\))

Fig 19. Estimates of the basic reproduction number by dataset and model
candidate. Error bars denote 95% credible intervals. The value in the
middle of the bars indicates the distance between the marginal
posterior’s mean and the true value (vertical line).

###### 1.2.3.3.2 Initial number of infectious individuals (\(I\_0\))

Fig 20. Estimates of the initial number of infectious individuals by
dataset and model candidate. Error bars denote 95% credible intervals.
The value in the middle of the bars indicates the distance between the
marginal posterior’s mean and the true value (vertical line).

###### 1.2.3.3.3 Reporting rate (\(\rho\))

Fig 21. Estimates of the reporting rate by dataset and model candidate.
Error bars denote 95% credible intervals. The value in the middle of the
bars indicates the distance between the marginal posterior’s mean and
the true value (vertical line).

##### 1.2.4 Fitting \(D^{14}\)

###### 1.2.4.1 Incidence fit

Fig 22. Posterior predictive checks against latent incidence. Dots
denote synthetic data while lines indicate simulations from candidates’
ODE structure. We configure these structures using samples from the
posterior distribution.

Fig 23. Fit scores by candidate model and type of data. \(x\) denotes latent incidence, whereas \(y\) indicates observed incidence. Vertical
line denotes the mean.

###### 1.2.4.2 Joint posterior distribution

In this plot, we show an example of a joint posterior distribution.
Specifically, this posterior distribution was derived from fitting the
\(M^{14}\) candidate to one \(D^{14}\) incidence report.

Fig 24. Posterior distribution. The diagonal shows the posterior
marginal distributions (histograms). In the lower triangular part, the
joint posterior distribution of each possible combination of two
parameters is displayed (heatmaps). The upper triangular part shows the
correlation among parameters.

###### 1.2.4.3 Marginal posterior distributions

###### 1.2.4.3.1 Basic reproduction number (\(\Re\_0\))

Fig 25. Estimates of the basic reproduction number by dataset and model
candidate. Error bars denote 95% credible intervals. The value in the
middle of the bars indicates the distance between the marginal
posterior’s mean and the true value (vertical line).

###### 1.2.4.3.2 Initial number of infectious individuals (\(I\_0\))

Fig 26. Estimates of the initial number of infectious individuals by
dataset and model candidate. Error bars denote 95% credible intervals.
The value in the middle of the bars indicates the distance between the
marginal posterior’s mean and the true value (vertical line).

###### 1.2.4.3.3 Reporting rate (\(\rho\))

Fig 27. Estimates of the reporting rate by dataset and model candidate.
Error bars denote 95% credible intervals. The value in the middle of the
bars indicates the distance between the marginal posterior’s mean and
the true value (vertical line).

#### 1.3 Summary

##### 1.3.1 Coverage

Table 1. Coverage table

| D^ij^ | M^ij^ | ℜ_0_ | I_0_ | ρ |
| --- | --- | --- | --- | --- |
| 11 | 11 | 95% | 95% | 100% |
| 11 | 12 | 0% | 95% | 30% |
| 11 | 13 | 0% | 95% | 0% |
| 11 | 14 | 0% | 90% | 0% |
| 12 | 11 | 0% | 90% | 15% |
| 12 | 12 | 90% | 95% | 100% |
| 12 | 13 | 5% | 95% | 95% |
| 12 | 14 | 0% | 95% | 75% |
| 13 | 11 | 0% | 85% | 5% |
| 13 | 12 | 0% | 90% | 95% |
| 13 | 13 | 95% | 100% | 100% |
| 13 | 14 | 35% | 100% | 100% |
| 14 | 11 | 0% | 85% | 0% |
| 14 | 12 | 0% | 90% | 60% |
| 14 | 13 | 25% | 95% | 90% |
| 14 | 14 | 85% | 95% | 95% |

##### 1.3.2 MLE criterion

Each bar (column) in the chart below represents the number of times
that particular model candidate attained the largest likelihood score
for a given set of incidence reports. Recall that each set of reports
comprises 20 time-series. For instance, the first column (left to right)
in the first panel (top-left) indicates that \(M^{11}\) outperforms its competitors 12 out
of 20 times (60 %) in fitting \(D^{11}\) incidence reports.

Fig 28. Score summary

### 2 Four-unknown parameterisation

In this parameterisation, for each candidate, we assume one
additional unknown parameter: the recovery rate (\(\gamma\)). Furthermore, all candidates are
coupled with the Poisson distribution.

#### 2.1 Prior distributions

Prior distributions for \(\beta\),
\(\rho\), and \(I^1\_0\) are identical to those described here.

##### 2.1.1 Recovery rate (\(\gamma\))

This prior distribution assumes that the magnitude of the mean
infectious period (\(\gamma^{-1}\)) is
at least one.

Fig 29. Histogram from samples obtained from the recovery rate’s prior
distribution

#### 2.2 Posterior distributions

##### 2.2.1 Fitting \(D^{11}\)

###### 2.2.1.1 Incidence fit

Fig 30. Posterior predictive checks against latent incidence. Dots
denote synthetic data while lines indicate simulations from candidates’
ODE structure. We configure these structures using samples from the
posterior distribution.

Fig 31. Fit scores by candidate model and type of data. \(x\) denotes latent incidence, whereas \(y\) indicates observed incidence. Vertical
line denotes the mean.

###### 2.2.1.2 Joint posterior distribution

Fig 32. Posterior distribution. The diagonal shows the posterior
marginal distributions (histograms). In the lower triangular part, the
joint posterior distribution of each possible combination of two
parameters is displayed (heatmaps). The upper triangular part shows the
correlation among parameters.

###### 2.2.1.3 Marginal posterior distributions

###### 2.2.1.3.1 Basic reproduction number (\(\Re\_0\))

Fig 33. Estimates of the basic reproduction number by dataset and model
candidate. Error bars denote 95% credible intervals. The value in the
middle of the bars indicates the distance between the marginal
posterior’s mean and the true value (vertical line).

###### 2.2.1.3.2 Reporting rate (\(\rho\))

Fig 34. Estimates of the reporting rate by dataset and model candidate.
Error bars denote 95% credible intervals. The value in the middle of the
bars indicates the distance between the marginal posterior’s mean and
the true value (vertical line).

###### 2.2.1.3.3 Initial number of infectious individuals (\(I\_0\))

Fig 35. Estimates of the initial number of infectious individuals by
dataset and model candidate. Error bars denote 95% credible intervals.
The value in the middle of the bars indicates the distance between the
marginal posterior’s mean and the true value (vertical line).

###### 2.2.1.3.4 Recovery rate (\(\gamma\))

For this parameter, we opt for a different visualisation.
Specifically, for each model candidate fitting a particular \(D^{11}\) incidence report, we compare the
marginal posterior distribution against the prior distribution via
histograms. These plots show that the data does little to inform the
prior distribution as it can be seen that both distributions agree to a
large extent along the parameter space. For this reason, we claim that
the four-unknown parameterisation is **unidentifiable**.
This statement is also supported by the ridge created by \(\beta\) and \(\gamma\) observed here.

Fig 36. Comparison between the recovery rate’s prior distribution (grey
histograms) against four marginal posterior distributions obtained from
each candidate (coloured histograms).

##### 2.2.2 Fitting \(D^{12}\)

###### 2.2.2.1 Incidence fit

Fig 37. Posterior predictive checks against latent incidence. Dots
denote synthetic data while lines indicate simulations from candidates’
ODE structure. We configure these structures using samples from the
posterior distribution.

Fig 38. Fit scores by candidate model and type of data. \(x\) denotes latent incidence, whereas \(y\) indicates observed incidence. Vertical
line denotes the mean.

###### 2.2.2.2 Joint posterior distribution

Fig 39. Posterior distribution. The diagonal shows the posterior
marginal distributions (histograms). In the lower triangular part, the
joint posterior distribution of each possible combination of two
parameters is displayed (heatmaps). The upper triangular part shows the
correlation among parameters.

###### 2.2.2.3 Marginal posterior distributions

###### 2.2.2.3.1 Basic reproduction number (\(\Re\_0\))

Fig 40. Estimates of the basic reproduction number by dataset and model
candidate. Error bars denote 95% credible intervals. The value in the
middle of the bars indicates the distance between the marginal
posterior’s mean and the true value (vertical line).

###### 2.2.2.3.2 Reporting rate (\(\rho\))

Fig 41. Estimates of the reporting rate by dataset and model candidate.
Error bars denote 95% credible intervals. The value in the middle of the
bars indicates the distance between the marginal posterior’s mean and
the true value (vertical line).

###### 2.2.2.3.3 Initial number of infectious individuals (\(I\_0\))

Fig 42. Estimates of the initial number of infectious individuals by
dataset and model candidate. Error bars denote 95% credible intervals.
The value in the middle of the bars indicates the distance between the
marginal posterior’s mean and the true value (vertical line).

###### 2.2.2.3.4 Recovery rate (\(\gamma\))

Fig 43. Comparison between the recovery rate’s prior distribution (grey
histograms) against four marginal posterior distributions obtained from
each candidate (coloured histograms).

##### 2.2.3 Fitting \(D^{13}\)

###### 2.2.3.1 Incidence fit

Fig 44. Posterior predictive checks against latent incidence. Dots
denote synthetic data while lines indicate simulations from candidates’
ODE structure. We configure these structures using samples from the
posterior distribution.

Fig 45. Fit scores by candidate model and type of data. \(x\) denotes latent incidence, whereas \(y\) indicates observed incidence. Vertical
line denotes the mean.

###### 2.2.3.2 Joint posterior distribution

Fig 46. Posterior distribution. The diagonal shows the posterior
marginal distributions (histograms). In the lower triangular part, the
joint posterior distribution of each possible combination of two
parameters is displayed (heatmaps). The upper triangular part shows the
correlation among parameters.

###### 2.2.3.3 Marginal posterior distributions

###### 2.2.3.3.1 Basic reproduction number (\(\Re\_0\))

Fig 47. Estimates of the basic reproduction number by dataset and model
candidate. Error bars denote 95% credible intervals. The value in the
middle of the bars indicates the distance between the marginal
posterior’s mean and the true value (vertical line).

###### 2.2.3.3.2 Reporting rate (\(\rho\))

Fig 48. Estimates of the reporting rate by dataset and model candidate.
Error bars denote 95% credible intervals. The value in the middle of the
bars indicates the distance between the marginal posterior’s mean and
the true value (vertical line).

###### 2.2.3.3.3 Initial number of infectious individuals (\(I\_0\))

Fig 49. Estimates of the initial number of infectious individuals by
dataset and model candidate. Error bars denote 95% credible intervals.
The value in the middle of the bars indicates the distance between the
marginal posterior’s mean and the true value (vertical line).

###### 2.2.3.3.4 Recovery rate (\(\gamma\))

Fig 50. Comparison between the recovery rate’s prior distribution (grey
histograms) against four marginal posterior distributions obtained from
each candidate (coloured histograms).

##### 2.2.4 Fitting \(D^{14}\)

###### 2.2.4.1 Incidence fit

Fig 51. Posterior predictive checks against latent incidence. Dots
denote synthetic data while lines indicate simulations from candidates’
ODE structure. We configure these structures using samples from the
posterior distribution.

Fig 52. Fit scores by candidate model and type of data. \(x\) denotes latent incidence, whereas \(y\) indicates observed incidence. Vertical
line denotes the mean.

###### 2.2.4.2 Joint posterior distribution

Fig 53. Posterior distribution. The diagonal shows the posterior
marginal distributions (histograms). In the lower triangular part, the
joint posterior distribution of each possible combination of two
parameters is displayed (heatmaps). The upper triangular part shows the
correlation among parameters.

###### 2.2.4.3 Marginal posterior distributions

###### 2.2.4.3.1 Basic reproduction number (\(\Re\_0\))

Fig 54. Estimates of the basic reproduction number by dataset and model
candidate. Error bars denote 95% credible intervals. The value in the
middle of the bars indicates the distance between the marginal
posterior’s mean and the true value (vertical line).

###### 2.2.4.3.2 Reporting rate (\(\rho\))

Fig 55. Estimates of the reporting rate by dataset and model candidate.
Error bars denote 95% credible intervals. The value in the middle of the
bars indicates the distance between the marginal posterior’s mean and
the true value (vertical line).

###### 2.2.4.3.3 Initial number of infectious individuals (\(I\_0\))

Fig 56. Estimates of the initial number of infectious individuals by
dataset and model candidate. Error bars denote 95% credible intervals.
The value in the middle of the bars indicates the distance between the
marginal posterior’s mean and the true value (vertical line).

###### 2.2.4.3.4 Recovery rate (\(\gamma\))

Fig 57. Comparison between the recovery rate’s prior distribution (grey
histograms) against four marginal posterior distributions obtained from
each candidate (coloured histograms).

#### 2.3 Summary

##### 2.3.1 Coverage

Table 2. Coverage table

| D^ij^ | M^ij^ | ℜ_0_ | I_0_ | γ^-1^ | ρ |
| --- | --- | --- | --- | --- | --- |
| 11 | 11 | 100% | 100% | 100% | 100% |
| 11 | 12 | 95% | 100% | 100% | 100% |
| 11 | 13 | 65% | 100% | 100% | 70% |
| 11 | 14 | 50% | 95% | 100% | 60% |
| 12 | 11 | 95% | 100% | 100% | 95% |
| 12 | 12 | 100% | 100% | 100% | 100% |
| 12 | 13 | 95% | 100% | 100% | 95% |
| 12 | 14 | 95% | 100% | 100% | 90% |
| 13 | 11 | 95% | 95% | 100% | 95% |
| 13 | 12 | 95% | 95% | 100% | 95% |
| 13 | 13 | 100% | 95% | 100% | 100% |
| 13 | 14 | 100% | 95% | 100% | 100% |
| 14 | 11 | 70% | 95% | 95% | 80% |
| 14 | 12 | 95% | 90% | 100% | 90% |
| 14 | 13 | 100% | 100% | 100% | 95% |
| 14 | 14 | 100% | 100% | 100% | 100% |

##### 2.3.2 \(\Re\_0\) vs \(\tau\)

Fig 58. Scatterplot of the interaction between the mean generation time
and the basic reproduction number.

### 3 Three-unknown parameterisation (Alternative)

The *alternative parameterisation* refers to the algebraic
manipulation of the \(SE^iI^jR\)
framework so as to obtain a set equations wherein the basic reproduction
number (\(\Re\_0\)) and the mean
generation time (\(\tau\)) are explicit
parameters of the model. In doing so, \(\beta\) and \(\gamma\) become functions of other
variables rather than parameters. In these parameterisation, we assume
the basic reproduction number’s inverse, the reporting rate (\(\rho\)) and initial number of infectious
individuals (\(I^1\_0\)) as unknowns.
The remaining parameters are fixed to their true values.

\[\begin{equation}
\begin{aligned}
\beta &= \frac{2 j (\tau - \sigma^{-1})}{\Re\_0^{-1} (j +
1)}\\
\gamma &= \beta \Re\_0^{-1}
\end{aligned}
\end{equation}\]

\[\begin{equation}
\begin{aligned}
\dot{S\_t} &= -\frac{\beta S\_t \sum^j\_{k=1} I^k\_t}{N}\\
\dot{E^1\_t} &= \frac{\beta S\_t \sum^j\_{k=1} I^k\_t}{N} - i\sigma
E^1\_t \\
\dot{E^2\_t} &= i\sigma E^1\_t - i\sigma E^2\_t \\
\vdots \\
\dot{E^i\_t} &= i \sigma E^{i - 1}\_t - i \sigma E^i\_t \\
\dot{I^1\_t} &= i \sigma E^{i}\_t - \gamma I^1\_t \\
\dot{I^2\_t} &= j \gamma I^{1}\_t - \gamma I^2\_t \\
\vdots \\
\dot{I^j\_t} &= j \gamma I^{j - 1}\_t - j \gamma I^{j}\_t \\
\dot{R\_t} &= j \gamma I^j\_t
\end{aligned}
\end{equation}\]

#### 3.1 Prior distributions

The prior distributions shown below apply for all candidates from
this parameterisation. Prior distributions for \(\rho\), and \(I^1\_0\) are identical to those described here.

##### 3.1.1 The inverse of the basic reproduction number (\(\Re\_0^{-1}\))

Since model candidates are fitting outbreak-like incidence data, it
is warranted to assume that the basic reproduction number is higher than
one (\(\Re\_0 > 1\)). This
observation implies that the magnitude of its inverse (\(\Re\_0^{-1}\)) between 0 and 1.
Consequently, we formulate a prior that is consistent with such a
constraint.

Fig 59. Histogram from samples obtained from the prior distribution of
the basic reproduction number’s inverse.

#### 3.2 Posterior distributions

##### 3.2.1 Fitting \(D^{11}\)

###### 3.2.1.1 Incidence fit

Fig 60. Posterior predictive checks against latent incidence. Dots
denote synthetic data while lines indicate simulations from candidates’
ODE structure. We configure these structures using samples from the
posterior distribution.

Fig 61. Fit scores by candidate model and type of data. \(x\) denotes latent incidence, whereas \(y\) indicates observed incidence. Vertical
line denotes the mean.

###### 3.2.1.2 Joint posterior distribution

Fig 62. Posterior distribution. The diagonal shows the posterior
marginal distributions (histograms). In the lower triangular part, the
joint posterior distribution of each possible combination of two
parameters is displayed (heatmaps). The upper triangular part shows the
correlation among parameters.

###### 3.2.1.3 Marginal posterior distributions

###### 3.2.1.3.1 Basic reproduction number (\(\Re\_0\))

Fig 63. Estimates of the basic reproduction number by dataset and model
candidate. Error bars denote 95% credible intervals. The value in the
middle of the bars indicates the distance between the marginal
posterior’s mean and the true value (vertical line).

###### 3.2.1.3.2 Initial number of infectious individuals (\(I\_0\))

Fig 64. Estimates of the initial number of infectious individuals by
dataset and model candidate. Error bars denote 95% credible intervals.
The value in the middle of the bars indicates the distance between the
marginal posterior’s mean and the true value (vertical line).

##### 3.2.2 Fitting \(D^{12}\)

###### 3.2.2.1 Incidence fit

Fig 65. Posterior predictive checks against latent incidence. Dots
denote synthetic data while lines indicate simulations from candidates’
ODE structure. We configure these structures using samples from the
posterior distribution.

Fig 66. Fit scores by candidate model and type of data. \(x\) denotes latent incidence, whereas \(y\) indicates observed incidence. Vertical
line denotes the mean.

###### 3.2.2.2 Joint posterior distribution

Fig 67. Posterior distribution. The diagonal shows the posterior
marginal distributions (histograms). In the lower triangular part, the
joint posterior distribution of each possible combination of two
parameters is displayed (heatmaps). The upper triangular part shows the
correlation among parameters.

###### 3.2.2.3 Marginal posterior distributions

###### 3.2.2.3.1 Basic reproduction number (\(\Re\_0\))

Fig 68. Estimates of the basic reproduction number by dataset and model
candidate. Error bars denote 95% credible intervals. The value in the
middle of the bars indicates the distance between the marginal
posterior’s mean and the true value (vertical line).

###### 3.2.2.3.2 Initial number of infectious individuals (\(I\_0\))

Fig 69. Estimates of the initial number of infectious individuals by
dataset and model candidate. Error bars denote 95% credible intervals.
The value in the middle of the bars indicates the distance between the
marginal posterior’s mean and the true value (vertical line).

##### 3.2.3 Fitting \(D^{13}\)

###### 3.2.3.1 Incidence fit

Fig 70. Posterior predictive checks against latent incidence. Dots
denote synthetic data while lines indicate simulations from candidates’
ODE structure. We configure these structures using samples from the
posterior distribution.

Fig 71. Fit scores by candidate model and type of data. \(x\) denotes latent incidence, whereas \(y\) indicates observed incidence. Vertical
line denotes the mean.

###### 3.2.3.2 Joint posterior distribution

Fig 72. Posterior distribution. The diagonal shows the posterior
marginal distributions (histograms). In the lower triangular part, the
joint posterior distribution of each possible combination of two
parameters is displayed (heatmaps). The upper triangular part shows the
correlation among parameters.

###### 3.2.3.3 Marginal posterior distributions

###### 3.2.3.3.1 Basic reproduction number (\(\Re\_0\))

Fig 73. Estimates of the basic reproduction number by dataset and model
candidate. Error bars denote 95% credible intervals. The value in the
middle of the bars indicates the distance between the marginal
posterior’s mean and the true value (vertical line).

###### 3.2.3.3.2 Initial number of infectious individuals (\(I\_0\))

Fig 74. Estimates of the initial number of infectious individuals by
dataset and model candidate. Error bars denote 95% credible intervals.
The value in the middle of the bars indicates the distance between the
marginal posterior’s mean and the true value (vertical line).

##### 3.2.4 Fitting \(D^{14}\)

###### 3.2.4.1 Incidence fit

Fig 75. Posterior predictive checks against latent incidence. Dots
denote synthetic data while lines indicate simulations from candidates’
ODE structure. We configure these structures using samples from the
posterior distribution.

Fig 76. Fit scores by candidate model and type of data. \(x\) denotes latent incidence, whereas \(y\) indicates observed incidence. Vertical
line denotes the mean.

###### 3.2.4.2 Joint posterior distribution

Fig 77. Posterior distribution. The diagonal shows the posterior
marginal distributions (histograms). In the lower triangular part, the
joint posterior distribution of each possible combination of two
parameters is displayed (heatmaps). The upper triangular part shows the
correlation among parameters.

###### 3.2.4.3 Marginal posterior distributions

###### 3.2.4.3.1 Basic reproduction number (\(\Re\_0\))

Fig 78. Estimates of the basic reproduction number by dataset and model
candidate. Error bars denote 95% credible intervals. The value in the
middle of the bars indicates the distance between the marginal
posterior’s mean and the true value (vertical line).

###### 3.2.4.3.2 Initial number of infectious individuals (\(I\_0\))

Fig 79. Estimates of the initial number of infectious individuals by
dataset and model candidate. Error bars denote 95% credible intervals.
The value in the middle of the bars indicates the distance between the
marginal posterior’s mean and the true value (vertical line).

#### 3.3 Summary

##### 3.3.1 Coverage

Table 3. Coverage table

| D^ij^ | M^ij^ | ℜ_0_ | I_0_ | ρ |
| --- | --- | --- | --- | --- |
| 11 | 11 | 95% | 95% | 100% |
| 11 | 12 | 95% | 45% | 95% |
| 11 | 13 | 90% | 10% | 95% |
| 11 | 14 | 85% | 5% | 95% |
| 12 | 11 | 90% | 65% | 100% |
| 12 | 12 | 90% | 95% | 100% |
| 12 | 13 | 95% | 85% | 100% |
| 12 | 14 | 95% | 75% | 100% |
| 13 | 11 | 75% | 50% | 100% |
| 13 | 12 | 95% | 95% | 100% |
| 13 | 13 | 95% | 95% | 100% |
| 13 | 14 | 100% | 90% | 100% |
| 14 | 11 | 90% | 20% | 100% |
| 14 | 12 | 90% | 60% | 100% |
| 14 | 13 | 85% | 80% | 100% |
| 14 | 14 | 85% | 95% | 95% |
