## Supplementary material for "Anchoring the mean generation time in the SEIR to mitigate biases in ℜ_0_ estimates due to uncertainty in the distribution of the epidemiological delays": S3.html


# S3

This appendix illustrates the process of fitting various
parameterisations of the \(SE^1I^jR\)
model (\(M^{1j}\)) to
**low-fidelity \(D^{1j}\)** incidence reports. We
mean by *parameterisation* the decision of categorising
parameters as either *unknown* or *assumed*. For the
unknown parameters, we construct prior distributions, which will be
eventually updated in light of the data via HMC sampling, resulting in a
posterior distribution. On the other hand, assumed parameters are fixed
at their true values.

### 1 Three-unknown

For candidates from this parameterisation, we assume that two
parameters and one initial condition are unknown: \(\beta\), \(\rho\), and \(I^1\_0\). Each candidate is coupled with the
appropriate likelihood function. In this case, given that the data is
overdispersed, such likelihood function corresponds to the Negative
Binomial distribution. The selection of this function entails the
estimation of an additional parameter, the degree of concentration
(\(\phi^{-1}\)) around the incidence’s
expected value.

#### 1.1 Prior distributions

The prior distributions shown below apply for all candidates from
this parameterisation.

##### 1.1.1 Effective contact rate (\(\beta\))

Fig 1. Histogram from samples obtained from the contact rate’s prior
distribution

##### 1.1.2 Reporting rate (\(\rho\))

Fig 2. Histogram from samples obtained from the reporting rate’s prior
distribution

##### 1.1.3 Initial number of infectious individuals (\(I\_0\))

Fig 3. Histogram from samples obtained from the prior distribution of
the initial number of infectious individuals.

##### 1.1.4 Overdispersion parameter (\(\phi^{-1}\))

Fig 4. Histogram from samples obtained from the overdispersion
parameter’s prior distribution.

#### 1.2 Posterior distributions

We stratify inference results by the infectious period distribution
that produced the observed incidences. For instance, \(D\_{11}\) implies that candidate models
fitted incidence reports that stem from the \(SE^1I^1R\).

###### 1.2.1.3.4 Overdispersion parameter (\(\phi^{-1}\))

Fig 11. Estimates of overdispersion by dataset and model candidate.
Error bars denote 95% credible intervals. The value in the middle of the
bars indicates the distance between the marginal posterior’s mean and
the true value (vertical line).

##### 1.2.2 Fitting \(D^{12}\)

###### 1.2.2.1 Incidence fit

Fig 12. Posterior predictive checks against latent incidence. Dots
denote synthetic data while lines indicate simulations from candidates’
ODE structure. We configure these structures using samples from the
posterior distribution.

Fig 13. Fit scores by candidate model and type of data. \(x\) denotes latent incidence, whereas \(y\) indicates observed incidence. Vertical
line denotes the mean.

###### 1.2.2.2 Joint posterior distribution

In this plot, we show an example of a joint posterior distribution.
Specifically, this posterior distribution was derived from fitting the
\(M^{12}\) candidate to one \(D^{12}\) incidence report.

Fig 14. Posterior distribution. The diagonal shows the posterior
marginal distributions (histograms). In the lower triangular part, the
joint posterior distribution of each possible combination of two
parameters is displayed (heatmaps). The upper triangular part shows the
correlation among parameters.

###### 1.2.2.3 Marginal posterior distributions

###### 1.2.2.3.1 Basic reproduction number (\(\Re\_0\))

Fig 15. Estimates of the basic reproduction number by dataset and model
candidate. Error bars denote 95% credible intervals. The value in the
middle of the bars indicates the distance between the marginal
posterior’s mean and the true value (vertical line).

###### 1.2.2.3.2 Reporting rate (\(\rho\))

Fig 16. Estimates of the reporting rate by dataset and model candidate.
Error bars denote 95% credible intervals. The value in the middle of the
bars indicates the distance between the marginal posterior’s mean and
the true value (vertical line).

###### 1.2.2.3.3 Initial number of infectious individuals (\(I\_0\))

Fig 17. Estimates of the initial number of infectious individuals by
dataset and model candidate. Error bars denote 95% credible intervals.
The value in the middle of the bars indicates the distance between the
marginal posterior’s mean and the true value (vertical line).

###### 1.2.2.3.4 Overdispersion parameter (\(\phi^{-1}\))

Fig 18. Estimates of overdispersion by dataset and model candidate.
Error bars denote 95% credible intervals. The value in the middle of the
bars indicates the distance between the marginal posterior’s mean and
the true value (vertical line).

##### 1.2.3 Fitting \(D^{13}\)

###### 1.2.3.1 Incidence fit

Fig 19. Posterior predictive checks against latent incidence. Dots
denote synthetic data while lines indicate simulations from candidates’
ODE structure. We configure these structures using samples from the
posterior distribution.

Fig 20. Fit scores by candidate model and type of data. \(x\) denotes latent incidence, whereas \(y\) indicates observed incidence. Vertical
line denotes the mean.

###### 1.2.3.2 Joint posterior distribution

In this plot, we show an example of a joint posterior distribution.
Specifically, this posterior distribution was derived from fitting the
\(M^{13}\) candidate to one \(D^{13}\) incidence report.

Fig 21. Posterior distribution. The diagonal shows the posterior
marginal distributions (histograms). In the lower triangular part, the
joint posterior distribution of each possible combination of two
parameters is displayed (heatmaps). The upper triangular part shows the
correlation among parameters.

###### 1.2.3.3 Marginal posterior distributions

###### 1.2.3.3.1 Basic reproduction number (\(\Re\_0\))

Fig 22. Estimates of the basic reproduction number by dataset and model
candidate. Error bars denote 95% credible intervals. The value in the
middle of the bars indicates the distance between the marginal
posterior’s mean and the true value (vertical line).

###### 1.2.3.3.2 Reporting rate (\(\rho\))

Fig 23. Estimates of the reporting rate by dataset and model candidate.
Error bars denote 95% credible intervals. The value in the middle of the
bars indicates the distance between the marginal posterior’s mean and
the true value (vertical line).

###### 1.2.3.3.3 Initial number of infectious individuals (\(I\_0\))

Fig 24. Estimates of the initial number of infectious individuals by
dataset and model candidate. Error bars denote 95% credible intervals.
The value in the middle of the bars indicates the distance between the
marginal posterior’s mean and the true value (vertical line).

###### 1.2.3.3.4 Overdispersion parameter (\(\phi^{-1}\))

Fig 25. Estimates of overdispersion by dataset and model candidate.
Error bars denote 95% credible intervals. The value in the middle of the
bars indicates the distance between the marginal posterior’s mean and
the true value (vertical line).

##### 1.2.4 Fitting \(D^{14}\)

###### 1.2.4.1 Incidence fit

Fig 26. Posterior predictive checks against latent incidence. Dots
denote synthetic data while lines indicate simulations from candidates’
ODE structure. We configure these structures using samples from the
posterior distribution.

Fig 27. Fit scores by candidate model and type of data. \(x\) denotes latent incidence, whereas \(y\) indicates observed incidence. Vertical
line denotes the mean.

###### 1.2.4.2 Joint posterior distribution

In this plot, we show an example of a joint posterior distribution.
Specifically, this posterior distribution was derived from fitting the
\(M^{14}\) candidate to one \(D^{14}\) incidence report.

Fig 28. Posterior distribution. The diagonal shows the posterior
marginal distributions (histograms). In the lower triangular part, the
joint posterior distribution of each possible combination of two
parameters is displayed (heatmaps). The upper triangular part shows the
correlation among parameters.

###### 1.2.4.3 Marginal posterior distributions

###### 1.2.4.3.1 Basic reproduction number (\(\Re\_0\))

Fig 29. Estimates of the basic reproduction number by dataset and model
candidate. Error bars denote 95% credible intervals. The value in the
middle of the bars indicates the distance between the marginal
posterior’s mean and the true value (vertical line).

###### 1.2.4.3.2 Reporting rate (\(\rho\))

Fig 30. Estimates of the reporting rate by dataset and model candidate.
Error bars denote 95% credible intervals. The value in the middle of the
bars indicates the distance between the marginal posterior’s mean and
the true value (vertical line).

###### 1.2.4.3.3 Initial number of infectious individuals (\(I\_0\))

Fig 31. Estimates of the initial number of infectious individuals by
dataset and model candidate. Error bars denote 95% credible intervals.
The value in the middle of the bars indicates the distance between the
marginal posterior’s mean and the true value (vertical line).

###### 1.2.4.3.4 Overdispersion parameter (\(\phi^{-1}\))

Fig 32. Estimates of overdispersion by dataset and model candidate.
Error bars denote 95% credible intervals. The value in the middle of the
bars indicates the distance between the marginal posterior’s mean and
the true value (vertical line).

#### 1.3 Summary

##### 1.3.1 Coverage

Table 1. Coverage table

| D^ij^ | M^ij^ | ℜ_0_ | I_0_ | ρ | φ^-1^ |
| --- | --- | --- | --- | --- | --- |
| 11 | 11 | 90% | 100% | 90% | 100% |
| 11 | 12 | 5% | 100% | 85% | 100% |
| 11 | 13 | 0% | 100% | 80% | 100% |
| 11 | 14 | 0% | 95% | 80% | 100% |
| 12 | 11 | 30% | 90% | 100% | 100% |
| 12 | 12 | 90% | 90% | 100% | 100% |
| 12 | 13 | 65% | 90% | 100% | 100% |
| 12 | 14 | 35% | 95% | 100% | 100% |
| 13 | 11 | 0% | 95% | 95% | 95% |
| 13 | 12 | 80% | 95% | 95% | 95% |
| 13 | 13 | 95% | 95% | 90% | 95% |
| 13 | 14 | 90% | 95% | 90% | 95% |
| 14 | 11 | 0% | 95% | 95% | 95% |
| 14 | 12 | 55% | 95% | 90% | 95% |
| 14 | 13 | 90% | 100% | 90% | 95% |
| 14 | 14 | 100% | 100% | 90% | 95% |

##### 1.3.2 MLE criterion

Each bar (column) in the chart below represents the number of times
that particular model candidate attained the largest likelihood score
for a given set of incidence reports. Recall that each set of reports
comprises 20 time-series. For instance, the first column (left to right)
in the first panel (top-left) indicates that \(M^{11}\) outperforms its competitors 10 out
of 20 times (50 %) in fitting \(D^{11}\) incidence reports.

Fig 33. Score summary

### 2 Four-unknown parameterisation

In this parameterisation, for each candidate, we assume one
additional unknown parameter: the recovery rate (\(\gamma\)). Furthermore, all candidates are
coupled with the Negative Binomial distribution.

#### 2.1 Prior distributions

Prior distributions for \(\beta\),
\(\rho\), \(I^1\_0\) and \(\phi^{-1}\) are identical to those
described here.

##### 2.1.1 Recovery rate (\(\gamma\))

This prior distribution assumes that the magnitude of the mean
infectious period (\(\gamma^{-1}\)) is
at least one.

#### 2.3 Summary

##### 2.3.1 Coverage

Table 2. Coverage table

| D^ij^ | M^ij^ | ℜ_0_ | I_0_ | γ^-1^ | ρ | φ^-1^ |
| --- | --- | --- | --- | --- | --- | --- |
| 11 | 11 | 100% | 100% | 100% | 95% | 100% |
| 11 | 12 | 100% | 100% | 100% | 90% | 100% |
| 11 | 13 | 100% | 100% | 100% | 90% | 100% |
| 11 | 14 | 95% | 100% | 100% | 90% | 100% |
| 12 | 11 | 100% | 95% | 100% | 100% | 100% |
| 12 | 12 | 100% | 100% | 100% | 100% | 100% |
| 12 | 13 | 100% | 100% | 100% | 100% | 100% |
| 12 | 14 | 100% | 100% | 100% | 100% | 100% |
| 13 | 11 | 95% | 95% | 100% | 90% | 95% |
| 13 | 12 | 100% | 95% | 100% | 95% | 95% |
| 13 | 13 | 100% | 100% | 100% | 95% | 95% |
| 13 | 14 | 100% | 100% | 100% | 95% | 95% |
| 14 | 11 | 95% | 100% | 100% | 95% | 95% |
| 14 | 12 | 100% | 100% | 100% | 95% | 90% |
| 14 | 13 | 100% | 100% | 100% | 90% | 95% |
| 14 | 14 | 100% | 100% | 100% | 90% | 95% |

##### 2.3.2 \(R\_0\) vs \(\tau\)

Fig 63. Scatterplot of the interaction between the mean generation time
and the basic reproduction number.

#### 3.3 Summary

##### 3.3.1 Coverage

Table 3. Coverage table

| D^ij^ | M^ij^ | ℜ_0_ | I_0_ | ρ | φ^-1^ |
| --- | --- | --- | --- | --- | --- |
| 11 | 11 | 95% | 100% | 80% | 100% |
| 11 | 12 | 95% | 85% | 85% | 100% |
| 11 | 13 | 95% | 80% | 85% | 100% |
| 11 | 14 | 95% | 75% | 80% | 100% |
| 12 | 11 | 90% | 100% | 100% | 100% |
| 12 | 12 | 90% | 90% | 100% | 100% |
| 12 | 13 | 90% | 90% | 100% | 100% |
| 12 | 14 | 90% | 85% | 100% | 100% |
| 13 | 11 | 90% | 95% | 95% | 95% |
| 13 | 12 | 95% | 95% | 95% | 95% |
| 13 | 13 | 95% | 95% | 95% | 95% |
| 13 | 14 | 95% | 95% | 95% | 90% |
| 14 | 11 | 90% | 90% | 90% | 90% |
| 14 | 12 | 95% | 100% | 90% | 95% |
| 14 | 13 | 95% | 100% | 90% | 90% |
| 14 | 14 | 100% | 100% | 90% | 95% |
