## Supplementary material for "Anchoring the mean generation time in the SEIR to mitigate biases in ℜ_0_ estimates due to uncertainty in the distribution of the epidemiological delays": S4.html


# S4

This appendix illustrates the process of fitting various
parameterisations of the \(SE^iI^jR\)
model (\(M^{1j}\) & \(M^{3j}\)) to **high-fidelity \(D^{3j}\)** incidence reports. We
mean by *parameterisation* the decision of categorising
parameters as either *unknown* or *assumed*. For the
unknown parameters, we construct prior distributions, which will be
eventually updated in light of the data via HMC sampling, resulting in a
posterior distribution. On the other hand, assumed parameters are fixed
at their true values.

### 1 Three-unknown parameterisation (traditional)

For candidates from this parameterisation, we assume that two
parameters and one initial condition are unknown: \(\beta\), \(\rho\), and \(I^1\_0\). Each candidate is coupled with the
Poisson distribution.

#### 1.1 Prior distributions

The prior distributions shown below apply for all candidates from
this parameterisation.

##### 1.1.1 Effective contact rate (\(\beta\))

Fig 1. Histogram from samples obtained from the contact rate’s prior
distribution

##### 1.1.2 Reporting rate (\(\rho\))

Fig 2. Histogram from samples obtained from the reporting rate’s prior
distribution

##### 1.1.3 Initial number of infectious individuals (\(I\_0\))

Fig 3. Histogram from samples obtained from the prior distribution of
the initial number of infectious individuals.

#### 1.2 Posterior distributions

We stratify inference results by the distribution of the infectious
period that produced the observed incidences. For instance, \(D\_{31}\) implies that the fitted incidence
stems from an \(SE^3I^1R\) with an
gamma-distributed (shape = 3) latent period and an
exponential-distributed infectious period.

##### 1.2.1 Fitting \(D^{31}\)

###### 1.2.1.1 Incidence fit

The figure below compares actual (points) and simulated (lines)
latent incidence by model candidate.

Fig 4. Posterior predictive checks against latent incidence. Dots denote
synthetic data while lines indicate simulations from candidates’ ODE
structure. We configure these structures using samples from the
posterior distribution.

We draw on the mean absolute scaled error (MASE) to compare incidence
fits. This quantity is a measure of the accuracy of forecasts,
well-suited for time-series. Therefore, we employ the MASE to compare
each of the 4000 simulated latent incidences against its true
counterpart (\(x\)) and observed
incidence (\(y\)). By simulated latent
incidences, we refer to the process of plugging the samples of a
posterior distribution into an ODE model to obtain incidence
trajectories via simulation. We summarise the results via histograms in
the plot below. The left column (of panels) contains the comparison
between simulated and actual latent incidences. The right column of
panels displays the comparison between simulated latent incidence and
the observed incidence. Overall, there is no noticeable difference in
the histograms as the distribution of the infectious period in the
fitting model varies (increasing \(j\)). In stark contrast, assuming the wrong
distribution of the latent period leads to suboptimal incidence fits.
Notice that assuming a gamma-distributed latent period with shape 3
(\(i = 3\)) produces better fits (lower
MASE) than a model with the same distribution of the infectious period,
but with an exponentially-distributed latent period (\(i = 1\)).

Fig 5. Fit scores by candidate model and type of data. \(x\) denotes latent incidence, whereas \(y\) indicates observed incidence. Vertical
line denotes the mean.

###### 1.2.1.2 Joint posterior distribution (right latent period)

In this plot, we show an example of a joint posterior distribution.
Specifically, this posterior distribution was derived from fitting the
\(M^{31}\) candidate to one \(D^{31}\) incidence report.

Fig 6. Posterior distribution. The diagonal shows the posterior marginal
distributions (histograms). In the lower triangular part, the joint
posterior distribution of each possible combination of two parameters is
displayed (heatmaps). The upper triangular part shows the correlation
among parameters.

###### 1.2.1.3 Joint posterior distribution (wrong latent period)

This posterior distribution was obtained from fitting the \(M^{11}\) candidate to one \(D^{31}\) incidence report.

Fig 7. Posterior distribution. The diagonal shows the posterior marginal
distributions (histograms). In the lower triangular part, the joint
posterior distribution of each possible combination of two parameters is
displayed (heatmaps). The upper triangular part shows the correlation
among parameters.

###### 1.2.1.4 Marginal posterior distributions

In this section, we present a comparison between estimated marginal
posterior distributions (error bars) and actual values (vertical lines).
Such distributions are the result of fitting all eight candidates to
four \(D^{31}\) incidence reports. The
value in the middle of the error bars indicates the relative error
between the actual value and the mean of the marginal posterior
distribution.

###### 1.2.1.4.1 Basic reproduction number (\(\Re\_0\))

Fig 8. Estimates of the basic reproduction number by dataset and model
candidate. Error bars denote 95% credible intervals. The value in the
middle of the bars indicates the distance between the marginal
posterior’s mean and the true value (vertical line).

###### 1.2.1.4.2 Reporting rate (\(\rho\))

Fig 9. Estimates of the reporting rate by dataset and model candidate.
Error bars denote 95% credible intervals. The value in the middle of the
bars indicates the distance between the marginal posterior’s mean and
the true value (vertical line).

###### 1.2.1.4.3 Initial number of infectious individuals (\(I\_0\))

Fig 10. Estimates of the initial number of infectious individuals by
dataset and model candidate. Error bars denote 95% credible intervals.
The value in the middle of the bars indicates the distance between the
marginal posterior’s mean and the true value (vertical line).

##### 1.2.2 Fitting \(D^{32}\)

###### 1.2.2.1 Incidence fit

Fig 11. Posterior predictive checks against latent incidence. Dots
denote synthetic data while lines indicate simulations from candidates’
ODE structure. We configure these structures using samples from the
posterior distribution.

Fig 12. Fit scores by candidate model and type of data. \(x\) denotes latent incidence, whereas \(y\) indicates observed incidence. Vertical
line denotes the mean.

###### 1.2.2.2 Joint posterior distribution (right latent period)

This posterior distribution was derived from fitting the \(M^{32}\) candidate to one \(D^{32}\) incidence report.

Fig 13. Posterior distribution. The diagonal shows the posterior
marginal distributions (histograms). In the lower triangular part, the
joint posterior distribution of each possible combination of two
parameters is displayed (heatmaps). The upper triangular part shows the
correlation among parameters.

###### 1.2.2.3 Joint posterior distribution (wrong latent period)

This posterior distribution was obtained from fitting the \(M^{12}\) candidate to one \(D^{32}\) incidence report.

Fig 14. Posterior distribution. The diagonal shows the posterior
marginal distributions (histograms). In the lower triangular part, the
joint posterior distribution of each possible combination of two
parameters is displayed (heatmaps). The upper triangular part shows the
correlation among parameters.

###### 1.2.2.4 Marginal posterior distributions

###### 1.2.2.4.1 Basic reproduction number (\(\Re\_0\))

Fig 15. Estimates of the basic reproduction number by dataset and model
candidate. Error bars denote 95% credible intervals. The value in the
middle of the bars indicates the distance between the marginal
posterior’s mean and the true value (vertical line).

###### 1.2.2.4.2 Reporting rate (\(\rho\))

Fig 16. Estimates of the reporting rate by dataset and model candidate.
Error bars denote 95% credible intervals. The value in the middle of the
bars indicates the distance between the marginal posterior’s mean and
the true value (vertical line).

###### 1.2.2.4.3 Initial number of infectious individuals (\(I\_0\))

Fig 17. Estimates of the initial number of infectious individuals by
dataset and model candidate. Error bars denote 95% credible intervals.
The value in the middle of the bars indicates the distance between the
marginal posterior’s mean and the true value (vertical line).

##### 1.2.3 Fitting \(D^{33}\)

###### 1.2.3.1 Incidence fit

Fig 18. Posterior predictive checks against latent incidence. Dots
denote synthetic data while lines indicate simulations from candidates’
ODE structure. We configure these structures using samples from the
posterior distribution.

Fig 19. Fit scores by candidate model and type of data. \(x\) denotes latent incidence, whereas \(y\) indicates observed incidence. Vertical
line denotes the mean.

###### 1.2.3.2 Joint posterior distribution (right latent period)

This posterior distribution was derived from fitting the \(M^{33}\) candidate to one \(D^{33}\) incidence report.

Fig 20. Posterior distribution. The diagonal shows the posterior
marginal distributions (histograms). In the lower triangular part, the
joint posterior distribution of each possible combination of two
parameters is displayed (heatmaps). The upper triangular part shows the
correlation among parameters.

###### 1.2.3.3 Joint posterior distribution (wrong latent period)

This posterior distribution was obtained from fitting the \(M^{13}\) candidate to one \(D^{33}\) incidence report.

Fig 21. Posterior distribution. The diagonal shows the posterior
marginal distributions (histograms). In the lower triangular part, the
joint posterior distribution of each possible combination of two
parameters is displayed (heatmaps). The upper triangular part shows the
correlation among parameters.

###### 1.2.3.4 Marginal posterior distributions

###### 1.2.3.4.1 Basic reproduction number (\(\Re\_0\))

Fig 22. Estimates of the basic reproduction number by dataset and model
candidate. Error bars denote 95% credible intervals. The value in the
middle of the bars indicates the distance between the marginal
posterior’s mean and the true value (vertical line).

###### 1.2.3.4.2 Reporting rate (\(\rho\))

Fig 23. Estimates of the reporting rate by dataset and model candidate.
Error bars denote 95% credible intervals. The value in the middle of the
bars indicates the distance between the marginal posterior’s mean and
the true value (vertical line).

###### 1.2.3.4.3 Initial number of infectious individuals (\(I\_0\))

Fig 24. Estimates of the initial number of infectious individuals by
dataset and model candidate. Error bars denote 95% credible intervals.
The value in the middle of the bars indicates the distance between the
marginal posterior’s mean and the true value (vertical line).

##### 1.2.4 Fitting \(D^{34}\)

###### 1.2.4.1 Incidence fit

Fig 25. Posterior predictive checks against latent incidence. Dots
denote synthetic data while lines indicate simulations from candidates’
ODE structure. We configure these structures using samples from the
posterior distribution.

Fig 26. Fit scores by candidate model and type of data. \(x\) denotes latent incidence, whereas \(y\) indicates observed incidence. Vertical
line denotes the mean.

###### 1.2.4.2 Joint posterior distribution (right latent period)

This posterior distribution was derived from fitting the \(M^{34}\) candidate to one \(D^{34}\) incidence report.

Fig 27. Posterior distribution. The diagonal shows the posterior
marginal distributions (histograms). In the lower triangular part, the
joint posterior distribution of each possible combination of two
parameters is displayed (heatmaps). The upper triangular part shows the
correlation among parameters.

###### 1.2.4.3 Joint posterior distribution (wrong \(j\))

This posterior distribution was obtained from fitting the \(M^{14}\) candidate to one \(D^{34}\) incidence report.

Fig 28. Posterior distribution. The diagonal shows the posterior
marginal distributions (histograms). In the lower triangular part, the
joint posterior distribution of each possible combination of two
parameters is displayed (heatmaps). The upper triangular part shows the
correlation among parameters.

###### 1.2.4.4 Marginal posterior distributions

###### 1.2.4.4.1 Basic reproduction number (\(\Re\_0\))

Fig 29. Estimates of the basic reproduction number by dataset and model
candidate. Error bars denote 95% credible intervals. The value in the
middle of the bars indicates the distance between the marginal
posterior’s mean and the true value (vertical line).

###### 1.2.4.4.2 Reporting rate (\(\rho\))

Fig 30. Estimates of the reporting rate by dataset and model candidate.
Error bars denote 95% credible intervals. The value in the middle of the
bars indicates the distance between the marginal posterior’s mean and
the true value (vertical line).

###### 1.2.4.4.3 Initial number of infectious individuals (\(I\_0\))

Fig 31. Estimates of the initial number of infectious individuals by
dataset and model candidate. Error bars denote 95% credible intervals.
The value in the middle of the bars indicates the distance between the
marginal posterior’s mean and the true value (vertical line).

##### 1.2.5 Summary

###### 1.2.5.1 Coverage

Table 1. Coverage table

| D^ij^ | M^ij^ | ℜ_0_ | I_0_ | ρ |
| --- | --- | --- | --- | --- |
| 31 | 31 | 90% | 90% | 90% |
| 31 | 32 | 0% | 95% | 15% |
| 31 | 33 | 0% | 100% | 5% |
| 31 | 34 | 0% | 100% | 5% |
| 31 | 11 | 95% | 0% | 90% |
| 31 | 12 | 0% | 0% | 15% |
| 31 | 13 | 0% | 0% | 5% |
| 31 | 14 | 0% | 0% | 5% |
| 32 | 31 | 0% | 100% | 35% |
| 32 | 32 | 95% | 100% | 95% |
| 32 | 33 | 0% | 100% | 85% |
| 32 | 34 | 0% | 100% | 65% |
| 32 | 11 | 0% | 0% | 35% |
| 32 | 12 | 100% | 0% | 95% |
| 32 | 13 | 0% | 0% | 90% |
| 32 | 14 | 0% | 0% | 65% |
| 33 | 31 | 0% | 95% | 10% |
| 33 | 32 | 0% | 95% | 85% |
| 33 | 33 | 95% | 95% | 90% |
| 33 | 34 | 25% | 95% | 85% |
| 33 | 11 | 0% | 0% | 10% |
| 33 | 12 | 0% | 0% | 85% |
| 33 | 13 | 90% | 0% | 95% |
| 33 | 14 | 55% | 0% | 85% |
| 34 | 31 | 0% | 90% | 10% |
| 34 | 32 | 0% | 90% | 80% |
| 34 | 33 | 45% | 90% | 90% |
| 34 | 34 | 90% | 90% | 95% |
| 34 | 11 | 0% | 0% | 5% |
| 34 | 12 | 0% | 0% | 75% |
| 34 | 13 | 30% | 0% | 90% |
| 34 | 14 | 85% | 0% | 95% |

### 2 Four-unknown parameterisation

In this parameterisation, for each candidate, we assume one
additional unknown parameter: the recovery rate (\(\gamma\)). Furthermore, all candidates are
coupled with the Poisson distribution. In light of the equivalency found
in the previous section, we restrict model candidates to those with an
exponentially-distributed latent period (\(i =
1\)).

###### 2.2.1.3.4 Recovery rate (\(\gamma\))

For this parameter, we opt for a different visualisation.
Specifically, for each model candidate fitting a particular \(D^{31}\) incidence report, we compare the
the marginal posterior distribution against the prior distribution via
histograms.

#### 2.3 Summary

##### 2.3.1 Coverage

Table 2. Coverage table

| D^ij^ | M^ij^ | ℜ_0_ | I_0_ | γ^-1^ | ρ |
| --- | --- | --- | --- | --- | --- |
| 31 | 11 | 15% | 40% | 15% | 10% |
| 31 | 12 | 25% | 80% | 5% | 25% |
| 31 | 13 | 30% | 95% | 0% | 25% |
| 31 | 14 | 40% | 95% | 0% | 25% |
| 32 | 11 | 0% | 30% | 0% | 0% |
| 32 | 12 | 0% | 30% | 0% | 0% |
| 32 | 13 | 0% | 35% | 0% | 0% |
| 32 | 14 | 0% | 40% | 0% | 0% |
| 33 | 11 | 0% | 15% | 0% | 0% |
| 33 | 12 | 0% | 15% | 0% | 0% |
| 33 | 13 | 0% | 40% | 0% | 0% |
| 33 | 14 | 0% | 40% | 0% | 0% |
| 34 | 11 | 0% | 15% | 0% | 0% |
| 34 | 12 | 0% | 10% | 0% | 0% |
| 34 | 13 | 0% | 15% | 0% | 0% |
| 34 | 14 | 0% | 25% | 0% | 0% |

##### 2.3.2 \(\Re\_0\) vs \(\tau\)

Fig 61. Scatterplot of the interaction between the mean generation time
and the basic reproduction number.

###### 3.2.1.2 Marginal posterior distributions

###### 3.2.1.2.1 Basic reproduction number (\(\Re\_0\))

Fig 66. Estimates of the basic reproduction number by dataset and model
candidate. Error bars denote 95% credible intervals. The value in the
middle of the bars indicates the distance between the marginal
posterior’s mean and the true value (vertical line).

###### 3.2.1.2.2 Initial number of infectious individuals (\(I\_0\))

Fig 67. Estimates of the initial number of infectious individuals by
dataset and model candidate. Error bars denote 95% credible intervals.
The value in the middle of the bars indicates the distance between the
marginal posterior’s mean and the true value (vertical line).

##### 3.2.2 Fitting \(D^{32}\)

###### 3.2.2.1 Incidence fit

Fig 68. Posterior predictive checks against latent incidence. Dots
denote synthetic data while lines indicate simulations from candidates’
ODE structure. We configure these structures using samples from the
posterior distribution.

Fig 69. Fit scores by candidate model and type of data. \(x\) denotes latent incidence, whereas \(y\) indicates observed incidence. Vertical
line denotes the mean.

###### 3.2.2.1.1 Joint posterior distribution

Fig 70. Posterior distribution. The diagonal shows the posterior
marginal distributions (histograms). In the lower triangular part, the
joint posterior distribution of each possible combination of two
parameters is displayed (heatmaps). The upper triangular part shows the
correlation among parameters.

###### 3.2.2.2 Marginal posterior distributions

###### 3.2.2.2.1 Basic reproduction number (\(\Re\_0\))

Fig 71. Estimates of the basic reproduction number by dataset and model
candidate. Error bars denote 95% credible intervals. The value in the
middle of the bars indicates the distance between the marginal
posterior’s mean and the true value (vertical line).

###### 3.2.2.2.2 Initial number of infectious individuals (\(I\_0\))

Fig 72. Estimates of the initial number of infectious individuals by
dataset and model candidate. Error bars denote 95% credible intervals.
The value in the middle of the bars indicates the distance between the
marginal posterior’s mean and the true value (vertical line).

##### 3.2.3 Fitting \(D^{33}\)

###### 3.2.3.1 Incidence fit

Fig 73. Posterior predictive checks against latent incidence. Dots
denote synthetic data while lines indicate simulations from candidates’
ODE structure. We configure these structures using samples from the
posterior distribution.

Fig 74. Fit scores by candidate model and type of data. \(x\) denotes latent incidence, whereas \(y\) indicates observed incidence. Vertical
line denotes the mean.

###### 3.2.3.1.1 Joint posterior distribution

Fig 75. Posterior distribution. The diagonal shows the posterior
marginal distributions (histograms). In the lower triangular part, the
joint posterior distribution of each possible combination of two
parameters is displayed (heatmaps). The upper triangular part shows the
correlation among parameters.

###### 3.2.3.2 Marginal posterior distributions

###### 3.2.3.2.1 Basic reproduction number (\(\Re\_0\))

Fig 76. Estimates of the basic reproduction number by dataset and model
candidate. Error bars denote 95% credible intervals. The value in the
middle of the bars indicates the distance between the marginal
posterior’s mean and the true value (vertical line).

###### 3.2.3.2.2 Initial number of infectious individuals (\(I\_0\))

Fig 77. Estimates of the initial number of infectious individuals by
dataset and model candidate. Error bars denote 95% credible intervals.
The value in the middle of the bars indicates the distance between the
marginal posterior’s mean and the true value (vertical line).

##### 3.2.4 Fitting \(D^{34}\)

###### 3.2.4.1 Incidence fit

Fig 78. Posterior predictive checks against latent incidence. Dots
denote synthetic data while lines indicate simulations from candidates’
ODE structure. We configure these structures using samples from the
posterior distribution.

Fig 79. Fit scores by candidate model and type of data. \(x\) denotes latent incidence, whereas \(y\) indicates observed incidence. Vertical
line denotes the mean.

###### 3.2.4.1.1 Joint posterior distribution

Fig 80. Posterior distribution. The diagonal shows the posterior
marginal distributions (histograms). In the lower triangular part, the
joint posterior distribution of each possible combination of two
parameters is displayed (heatmaps). The upper triangular part shows the
correlation among parameters.

###### 3.2.4.2 Marginal posterior distributions

###### 3.2.4.2.1 Basic reproduction number (\(\Re\_0\))

Fig 81. Estimates of the basic reproduction number by dataset and model
candidate. Error bars denote 95% credible intervals. The value in the
middle of the bars indicates the distance between the marginal
posterior’s mean and the true value (vertical line).

###### 3.2.4.2.2 Initial number of infectious individuals (\(I\_0\))

Fig 82. Estimates of the initial number of infectious individuals by
dataset and model candidate. Error bars denote 95% credible intervals.
The value in the middle of the bars indicates the distance between the
marginal posterior’s mean and the true value (vertical line).

#### 3.3 Summary

##### 3.3.1 Coverage

Table 3. Coverage table

| D^ij^ | M^ij^ | ℜ_0_ | I_0_ | ρ |
| --- | --- | --- | --- | --- |
| 31 | 11 | 95% | 0% | 90% |
| 31 | 12 | 90% | 0% | 90% |
| 31 | 13 | 70% | 0% | 90% |
| 31 | 14 | 55% | 0% | 95% |
| 32 | 11 | 85% | 0% | 90% |
| 32 | 12 | 100% | 0% | 90% |
| 32 | 13 | 85% | 0% | 95% |
| 32 | 14 | 85% | 0% | 95% |
| 33 | 11 | 95% | 0% | 90% |
| 33 | 12 | 90% | 0% | 95% |
| 33 | 13 | 90% | 0% | 95% |
| 33 | 14 | 85% | 0% | 95% |
| 34 | 11 | 90% | 0% | 90% |
| 34 | 12 | 95% | 0% | 95% |
| 34 | 13 | 90% | 0% | 95% |
| 34 | 14 | 85% | 0% | 95% |
