## Supplementary material for "Anchoring the mean generation time in the SEIR to mitigate biases in ℜ_0_ estimates due to uncertainty in the distribution of the epidemiological delays": S5.html

#### 1.3 Summary

##### 1.3.1 Coverage

Table 1. Coverage table

| D^ij^ | M^ij^ | ℜ_0_ | I_0_ | ρ | φ^-1^ |
| --- | --- | --- | --- | --- | --- |
| 31 | 31 | 95% | 95% | 95% | 95% |
| 31 | 32 | 0% | 95% | 95% | 95% |
| 31 | 33 | 0% | 95% | 90% | 95% |
| 31 | 34 | 0% | 95% | 90% | 95% |
| 31 | 11 | 90% | 65% | 95% | 95% |
| 31 | 12 | 0% | 60% | 95% | 95% |
| 31 | 13 | 0% | 55% | 95% | 95% |
| 31 | 14 | 0% | 55% | 95% | 95% |
| 32 | 31 | 5% | 95% | 100% | 100% |
| 32 | 32 | 100% | 100% | 100% | 100% |
| 32 | 33 | 80% | 100% | 100% | 100% |
| 32 | 34 | 45% | 100% | 100% | 100% |
| 32 | 11 | 10% | 45% | 95% | 95% |
| 32 | 12 | 100% | 40% | 100% | 95% |
| 32 | 13 | 85% | 30% | 100% | 95% |
| 32 | 14 | 60% | 30% | 100% | 95% |
| 33 | 31 | 0% | 80% | 95% | 85% |
| 33 | 32 | 80% | 90% | 95% | 85% |
| 33 | 33 | 85% | 90% | 95% | 85% |
| 33 | 34 | 75% | 90% | 95% | 85% |
| 33 | 11 | 0% | 70% | 85% | 90% |
| 33 | 12 | 80% | 60% | 90% | 90% |
| 33 | 13 | 90% | 60% | 95% | 90% |
| 33 | 14 | 85% | 60% | 95% | 85% |
| 34 | 31 | 0% | 90% | 95% | 95% |
| 34 | 32 | 55% | 90% | 95% | 95% |
| 34 | 33 | 90% | 85% | 100% | 90% |
| 34 | 34 | 90% | 85% | 100% | 90% |
| 34 | 11 | 0% | 60% | 90% | 95% |
| 34 | 12 | 55% | 45% | 95% | 100% |
| 34 | 13 | 90% | 45% | 95% | 100% |
| 34 | 14 | 95% | 45% | 100% | 95% |

###### 2.2.3.3.5 Recovery rate (\(\gamma\))

Fig 61. Comparison between the recovery rate’s prior distribution (grey
histograms) against four marginal posterior distributions obtained from
each candidate (coloured histograms).

###### 2.2.4.3.5 Recovery rate (\(\gamma\))

Fig 69. Comparison between the recovery rate’s prior distribution (grey
histograms) against four marginal posterior distributions obtained from
each candidate (coloured histograms).

#### 2.3 Summary

##### 2.3.1 Coverage

Table 2. Coverage table

| D^ij^ | M^ij^ | ℜ_0_ | I_0_ | γ^-1^ | ρ | φ^-1^ |
| --- | --- | --- | --- | --- | --- | --- |
| 31 | 11 | 95% | 95% | 95% | 95% | 95% |
| 31 | 12 | 95% | 100% | 95% | 100% | 95% |
| 31 | 13 | 100% | 100% | 90% | 100% | 95% |
| 31 | 14 | 100% | 100% | 90% | 100% | 95% |
| 32 | 11 | 100% | 100% | 100% | 95% | 100% |
| 32 | 12 | 100% | 100% | 100% | 100% | 100% |
| 32 | 13 | 100% | 100% | 100% | 95% | 100% |
| 32 | 14 | 100% | 100% | 100% | 100% | 100% |
| 33 | 11 | 65% | 95% | 90% | 90% | 80% |
| 33 | 12 | 85% | 95% | 90% | 90% | 80% |
| 33 | 13 | 85% | 95% | 90% | 90% | 80% |
| 33 | 14 | 90% | 95% | 80% | 95% | 85% |
| 34 | 11 | 65% | 90% | 95% | 90% | 95% |
| 34 | 12 | 80% | 90% | 95% | 90% | 100% |
| 34 | 13 | 80% | 90% | 80% | 90% | 95% |
| 34 | 14 | 80% | 90% | 80% | 90% | 95% |

###### 2.3.1.1 \(\Re\_0\) vs \(\tau\)

Fig 70. Scatterplot of the interaction between the mean generation time
and the basic reproduction number.

#### 3.3 Summary

##### 3.3.1 Coverage

Table 3. Coverage table

| D^ij^ | M^ij^ | ℜ_0_ | I_0_ | ρ | φ^-1^ |
| --- | --- | --- | --- | --- | --- |
| 31 | 11 | 95% | 70% | 95% | 95% |
| 31 | 12 | 95% | 80% | 95% | 95% |
| 31 | 13 | 95% | 85% | 95% | 95% |
| 31 | 14 | 95% | 90% | 95% | 95% |
| 32 | 11 | 100% | 25% | 100% | 95% |
| 32 | 12 | 100% | 35% | 95% | 95% |
| 32 | 13 | 100% | 45% | 95% | 95% |
| 32 | 14 | 100% | 45% | 100% | 95% |
| 33 | 11 | 85% | 60% | 95% | 90% |
| 33 | 12 | 90% | 60% | 95% | 90% |
| 33 | 13 | 85% | 60% | 95% | 85% |
| 33 | 14 | 90% | 70% | 95% | 90% |
| 34 | 11 | 85% | 40% | 95% | 85% |
| 34 | 12 | 95% | 45% | 95% | 95% |
| 34 | 13 | 100% | 50% | 95% | 100% |
| 34 | 14 | 95% | 45% | 95% | 95% |

### 4 Three-unknown (Alternative) - Poisson

In this section, we test the implications of amalgamating the
alternative parameterisation with a Poisson measurement model
(likelihood function) to fit overdispersed incidence data. The prior
distribution for these fitting candidates correspond to those described
in the previous section.

#### 4.1 Posterior distributions

##### 4.1.1 Fitting \(D^{31}\)

###### 4.1.1.1 Incidence fit

Fig 92. Posterior predictive checks against latent incidence. Dots
denote synthetic data while lines indicate simulations from candidates’
ODE structure. We configure these structures using samples from the
posterior distribution.

Fig 93. Fit scores by candidate model and type of data. \(x\) denotes latent incidence, whereas \(y\) indicates observed incidence. Vertical
line denotes the mean.

###### 4.1.1.2 Joint posterior distribution

Fig 94. Posterior distribution. The diagonal shows the posterior
marginal distributions (histograms). In the lower triangular part, the
joint posterior distribution of each possible combination of two
parameters is displayed (heatmaps). The upper triangular part shows the
correlation among parameters.

###### 4.1.1.3 Marginal posterior distributions

###### 4.1.1.3.1 Basic reproduction number (\(\Re\_0\))

Fig 95. Estimates of the basic reproduction number by dataset and model
candidate. Error bars denote 95% credible intervals. The value in the
middle of the bars indicates the distance between the marginal
posterior’s mean and the true value (vertical line).

##### 4.1.2 Fitting \(D^{32}\)

###### 4.1.2.1 Incidence fit

Fig 96. Posterior predictive checks against latent incidence. Dots
denote synthetic data while lines indicate simulations from candidates’
ODE structure. We configure these structures using samples from the
posterior distribution.

Fig 97. Fit scores by candidate model and type of data. \(x\) denotes latent incidence, whereas \(y\) indicates observed incidence. Vertical
line denotes the mean.

###### 4.1.2.2 Joint posterior distribution

Fig 98. Posterior distribution. The diagonal shows the posterior
marginal distributions (histograms). In the lower triangular part, the
joint posterior distribution of each possible combination of two
parameters is displayed (heatmaps). The upper triangular part shows the
correlation among parameters.

###### 4.1.2.3 Marginal posterior distributions

###### 4.1.2.3.1 Basic reproduction number (\(\Re\_0\))

Fig 99. Estimates of the basic reproduction number by dataset and model
candidate. Error bars denote 95% credible intervals. The value in the
middle of the bars indicates the distance between the marginal
posterior’s mean and the true value (vertical line).

##### 4.1.3 Fitting \(D^{33}\)

###### 4.1.3.1 Incidence fit

Fig 100. Posterior predictive checks against latent incidence. Dots
denote synthetic data while lines indicate simulations from candidates’
ODE structure. We configure these structures using samples from the
posterior distribution.

###### 4.1.3.2 Joint posterior distribution

Fig 101. Posterior distribution. The diagonal shows the posterior
marginal distributions (histograms). In the lower triangular part, the
joint posterior distribution of each possible combination of two
parameters is displayed (heatmaps). The upper triangular part shows the
correlation among parameters.

###### 4.1.3.3 Marginal posterior distributions

###### 4.1.3.3.1 Basic reproduction number (\(\Re\_0\))

Fig 102. Estimates of the basic reproduction number by dataset and model
candidate. Error bars denote 95% credible intervals. The value in the
middle of the bars indicates the distance between the marginal
posterior’s mean and the true value (vertical line).

##### 4.1.4 Fitting \(D^{34}\)

###### 4.1.4.1 Incidence fit

Fig 103. Posterior predictive checks against latent incidence. Dots
denote synthetic data while lines indicate simulations from candidates’
ODE structure. We configure these structures using samples from the
posterior distribution.

Fig 104. Fit scores by candidate model and type of data. \(x\) denotes latent incidence, whereas \(y\) indicates observed incidence. Vertical
line denotes the mean.

###### 4.1.4.2 Joint posterior distribution

Fig 105. Posterior distribution. The diagonal shows the posterior
marginal distributions (histograms). In the lower triangular part, the
joint posterior distribution of each possible combination of two
parameters is displayed (heatmaps). The upper triangular part shows the
correlation among parameters.

###### 4.1.4.3 Marginal posterior distributions

###### 4.1.4.3.1 Basic reproduction number (\(\Re\_0\))

Fig 106. Estimates of the basic reproduction number by dataset and model
candidate. Error bars denote 95% credible intervals. The value in the
middle of the bars indicates the distance between the marginal
posterior’s mean and the true value (vertical line).

#### 4.2 Summary

##### 4.2.1 Coverage

Table 4. Coverage table

| D^ij^ | M^ij^ | ℜ_0_ | I_0_ | ρ |
| --- | --- | --- | --- | --- |
| 31 | 11 | 15% | 20% | 30% |
| 31 | 12 | 15% | 15% | 30% |
| 31 | 13 | 15% | 15% | 30% |
| 31 | 14 | 15% | 10% | 25% |
| 32 | 11 | 25% | 5% | 20% |
| 32 | 12 | 30% | 0% | 20% |
| 32 | 13 | 30% | 0% | 20% |
| 32 | 14 | 30% | 0% | 20% |
| 33 | 11 | 5% | 15% | 10% |
| 33 | 12 | 5% | 20% | 10% |
| 33 | 13 | 10% | 25% | 10% |
| 33 | 14 | 10% | 25% | 10% |
| 34 | 11 | 5% | 5% | 0% |
| 34 | 12 | 10% | 5% | 0% |
| 34 | 13 | 10% | 5% | 0% |
| 34 | 14 | 10% | 5% | 5% |
