## Supplementary material for "Anchoring the mean generation time in the SEIR to mitigate biases in ℜ_0_ estimates due to uncertainty in the distribution of the epidemiological delays": S6.html


# S6

This appendix aims to illustrate the performance of the alternative
parameterisation under various transmissibility levels (\(\Re\_0\)) and mean generation times. For
simplicity, we restrict the \(SE^iI^jR\) to four instances (\(i = 1\), and \(j
= \{1,2,3,4\}\)) for data generation and fitting.

### 1 Overview

This sensitivity analysis consists of testing the alternative
parameterisation under **four** additional scenarios. The
first scenario corresponds to that presented in **S3**. By
*scenario*, we refer to the particular configuration of \(\Re\_0\) and the mean generation time (\(\tau\)) in the \(SE^iI^jR\) framework to produce synthetic
data. Since, \(\tau\) varies as the
distribution of the infectious period (\(j\)) changes, we identify each scenario by
the mean generation time obtained from an exponentially-distributed
infectious period (denoted by \(\tau\_e\)). For each instance and
data-fidelity level, we generate 20 synthetic incidence reports. Recall
that fidelity levels correspond to two configurations of the Negative
Binomial measurement model: high fidelity (\(\phi^{-1} = 0\)) & low fidelity (\(\phi^{-1} = \frac{1}{3}\)). As a result, we
produce *160* synthetic incidence reports per scenario. Then, we
fit four candidates per incidence report, where we assume the
appropriate measurement model.

Table 1. Scenarios for sensitivity analysis

| Scenario | ℜ_0_ | τ_e_ |
| --- | --- | --- |
| 1 | 2.5 | 4 |
| 2 | 2.5 | 8 |
| 3 | 2.5 | 13 |
| 4 | 9.0 | 4 |
| 5 | 17.0 | 4 |

### 2 Scenario 2

In comparison to Scenario 1, we increase the reference mean
generation time (\(\tau\_e\)) from
**4** to **8**.

#### 2.1 Synthetic data

Fig 1. Scenario 2’s simulated incidence reports. Measurement noise from
the Poisson (no overdispersion) & Negative Binomial distributions
was added to the smooth trajectories obtained from SEIR instances with
an exponential-distributed latent period.

#### 2.2 Inference

##### 2.2.1 Poisson data

###### 2.2.1.1 Fitting \(D^{11}\)

###### 2.2.1.1.1 Incidence fit

Fig 2. Posterior predictive checks against latent incidence. Dots denote
synthetic data while lines indicate simulations from candidates’ ODE
structure. We configure these structures using samples from the
posterior distribution.

Fig 3. Fit scores by candidate model and type of data. \(x\) denotes latent incidence, whereas \(y\) indicates observed incidence. Vertical
line denotes the mean.

###### 2.2.1.1.2 \(\Re\_0\) marginal distribution

Fig 4. Estimates of the basic reproduction number by dataset and model
candidate. Error bars denote 95% credible intervals. The value in the
middle of the bars indicates the distance between the marginal
posterior’s mean and the true value (vertical line).

In the figure below, we show the results of fitting the
**traditional** three-unknown parameter model. Here, we can
see that the bias caused by the misspecification of the infectious
period is larger than the one obtained from the alternative
parameterisation. In other words, it is costlier to misspecify the mean
generation time than the mean infectious period.

Fig 5. Estimates of the basic reproduction number by dataset and model
candidate. Error bars denote 95% credible intervals. The value in the
middle of the bars indicates the distance between the marginal
posterior’s mean and the true value (vertical line).

###### 2.2.1.1.3 \(I\_0\) marginal distribution

Fig 6. Estimates of the initial number of infectious individuals by
dataset and model candidate. Error bars denote 95% credible intervals.
The value in the middle of the bars indicates the distance between the
marginal posterior’s mean and the true value (vertical line).

###### 2.2.1.2 Fitting \(D^{12}\)

###### 2.2.1.2.1 Incidence fit

Fig 7. Posterior predictive checks against latent incidence. Dots denote
synthetic data while lines indicate simulations from candidates’ ODE
structure. We configure these structures using samples from the
posterior distribution.

Fig 8. Fit scores by candidate model and type of data. \(x\) denotes latent incidence, whereas \(y\) indicates observed incidence. Vertical
line denotes the mean.

###### 2.2.1.2.2 \(\Re\_0\) marginal distribution

Fig 9. Estimates of the basic reproduction number by dataset and model
candidate. Error bars denote 95% credible intervals. The value in the
middle of the bars indicates the distance between the marginal
posterior’s mean and the true value (vertical line).

###### 2.2.1.2.3 \(I\_0\) marginal distribution

Fig 10. Estimates of the initial number of infectious individuals by
dataset and model candidate. Error bars denote 95% credible intervals.
The value in the middle of the bars indicates the distance between the
marginal posterior’s mean and the true value (vertical line).

###### 2.2.1.3 Fitting \(D^{13}\)

###### 2.2.1.3.1 Incidence fit

Fig 11. Posterior predictive checks against latent incidence. Dots
denote synthetic data while lines indicate simulations from candidates’
ODE structure. We configure these structures using samples from the
posterior distribution.

Fig 12. Fit scores by candidate model and type of data. \(x\) denotes latent incidence, whereas \(y\) indicates observed incidence. Vertical
line denotes the mean.

###### 2.2.1.3.2 \(\Re\_0\) marginal distribution

Fig 13. Estimates of the basic reproduction number by dataset and model
candidate. Error bars denote 95% credible intervals. The value in the
middle of the bars indicates the distance between the marginal
posterior’s mean and the true value (vertical line).

###### 2.2.1.3.3 \(I\_0\) marginal distribution

Fig 14. Estimates of the initial number of infectious individuals by
dataset and model candidate. Error bars denote 95% credible intervals.
The value in the middle of the bars indicates the distance between the
marginal posterior’s mean and the true value (vertical line).

###### 2.2.1.4 Fitting \(D^{14}\)

###### 2.2.1.4.1 Incidence fit

Fig 16. Fit scores by candidate model and type of data. \(x\) denotes latent incidence, whereas \(y\) indicates observed incidence. Vertical
line denotes the mean.

###### 2.2.1.4.2 \(\Re\_0\) marginal distribution

Fig 17. Estimates of the basic reproduction number by dataset and model
candidate. Error bars denote 95% credible intervals. The value in the
middle of the bars indicates the distance between the marginal
posterior’s mean and the true value (vertical line).

###### 2.2.1.4.3 \(I\_0\) marginal distribution

Fig 18. Estimates of the initial number of infectious individuals by
dataset and model candidate. Error bars denote 95% credible intervals.
The value in the middle of the bars indicates the distance between the
marginal posterior’s mean and the true value (vertical line).

###### 2.2.1.5 Summary

###### 2.2.1.5.1 Coverage

Table 2. Scenario 2's coverage table. Poisson noise

| D^ij^ | M^ij^ | ℜ_0_ | I_0_ | ρ |
| --- | --- | --- | --- | --- |
| 11 | 11 | 95% | 95% | 90% |
| 11 | 12 | 80% | 35% | 95% |
| 11 | 13 | 35% | 10% | 95% |
| 11 | 14 | 30% | 0% | 90% |
| 12 | 11 | 80% | 45% | 80% |
| 12 | 12 | 95% | 95% | 80% |
| 12 | 13 | 90% | 80% | 80% |
| 12 | 14 | 75% | 65% | 80% |
| 13 | 11 | 60% | 25% | 85% |
| 13 | 12 | 95% | 85% | 85% |
| 13 | 13 | 95% | 95% | 90% |
| 13 | 14 | 100% | 85% | 95% |
| 14 | 11 | 60% | 25% | 100% |
| 14 | 12 | 75% | 70% | 100% |
| 14 | 13 | 75% | 85% | 95% |
| 14 | 14 | 90% | 85% | 95% |

##### 2.2.2 Overdispersed data

###### 2.2.2.1 Fitting \(D^{11}\)

###### 2.2.2.1.1 Incidence fit

Fig 19. Posterior predictive checks against latent incidence. Dots
denote synthetic data while lines indicate simulations from candidates’
ODE structure. We configure these structures using samples from the
posterior distribution.

Fig 20. Fit scores by candidate model and type of data. \(x\) denotes latent incidence, whereas \(y\) indicates observed incidence. Vertical
line denotes the mean.

###### 2.2.2.1.2 \(\Re\_0\) marginal distribution

Fig 21. Estimates of the basic reproduction number by dataset and model
candidate. Error bars denote 95% credible intervals. The value in the
middle of the bars indicates the distance between the marginal
posterior’s mean and the true value (vertical line).

###### 2.2.2.1.3 \(I\_0\) marginal distribution

Fig 22. Estimates of the initial number of infectious individuals by
dataset and model candidate. Error bars denote 95% credible intervals.
The value in the middle of the bars indicates the distance between the
marginal posterior’s mean and the true value (vertical line).

###### 2.2.2.2 Fitting \(D^{12}\)

Fig 24. Fit scores by candidate model and type of data. \(x\) denotes latent incidence, whereas \(y\) indicates observed incidence. Vertical
line denotes the mean.

###### 2.2.2.2.1 \(\Re\_0\) marginal distribution

Fig 25. Estimates of the basic reproduction number by dataset and model
candidate. Error bars denote 95% credible intervals. The value in the
middle of the bars indicates the distance between the marginal
posterior’s mean and the true value (vertical line).

###### 2.2.2.2.2 \(I\_0\) marginal distribution

Fig 26. Estimates of the initial number of infectious individuals by
dataset and model candidate. Error bars denote 95% credible intervals.
The value in the middle of the bars indicates the distance between the
marginal posterior’s mean and the true value (vertical line).

###### 2.2.2.3 Fitting \(D^{13}\)

Fig 28. Fit scores by candidate model and type of data. \(x\) denotes latent incidence, whereas \(y\) indicates observed incidence. Vertical
line denotes the mean.

###### 2.2.2.3.1 \(\Re\_0\) marginal distribution

Fig 29. Estimates of the basic reproduction number by dataset and model
candidate. Error bars denote 95% credible intervals. The value in the
middle of the bars indicates the distance between the marginal
posterior’s mean and the true value (vertical line).

###### 2.2.2.3.2 \(I\_0\) marginal distribution

Fig 30. Estimates of the initial number of infectious individuals by
dataset and model candidate. Error bars denote 95% credible intervals.
The value in the middle of the bars indicates the distance between the
marginal posterior’s mean and the true value (vertical line).

###### 2.2.2.4 Fitting \(D^{14}\)

Fig 32. Fit scores by candidate model and type of data. \(x\) denotes latent incidence, whereas \(y\) indicates observed incidence. Vertical
line denotes the mean.

###### 2.2.2.4.1 \(\Re\_0\) marginal distribution

Fig 33. Estimates of the basic reproduction number by dataset and model
candidate. Error bars denote 95% credible intervals. The value in the
middle of the bars indicates the distance between the marginal
posterior’s mean and the true value (vertical line).

###### 2.2.2.4.2 \(I\_0\) marginal distribution

Fig 34. Estimates of the initial number of infectious individuals by
dataset and model candidate. Error bars denote 95% credible intervals.
The value in the middle of the bars indicates the distance between the
marginal posterior’s mean and the true value (vertical line).

###### 2.2.2.5 Summary

###### 2.2.2.5.1 Coverage

Table 3. Scenario 2's coverage table. Overdispersion

| D^ij^ | M^ij^ | ℜ_0_ | I_0_ | ρ | φ^-1^ |
| --- | --- | --- | --- | --- | --- |
| 11 | 11 | 95% | 95% | 95% | 95% |
| 11 | 12 | 90% | 80% | 90% | 95% |
| 11 | 13 | 85% | 75% | 90% | 95% |
| 11 | 14 | 85% | 75% | 95% | 95% |
| 12 | 11 | 100% | 95% | 90% | 85% |
| 12 | 12 | 100% | 95% | 90% | 90% |
| 12 | 13 | 100% | 100% | 90% | 90% |
| 12 | 14 | 100% | 100% | 90% | 90% |
| 13 | 11 | 85% | 95% | 95% | 95% |
| 13 | 12 | 90% | 95% | 100% | 95% |
| 13 | 13 | 90% | 90% | 95% | 95% |
| 13 | 14 | 90% | 85% | 100% | 95% |
| 14 | 11 | 90% | 80% | 95% | 95% |
| 14 | 12 | 90% | 90% | 95% | 95% |
| 14 | 13 | 95% | 95% | 95% | 95% |
| 14 | 14 | 95% | 95% | 95% | 95% |

### 3 Scenario 3

In comparison to Scenario 2, we increase the reference mean
generation time (\(\tau\_e\)) from
**8** to **13**.

#### 3.1 Synthetic data

Fig 35. Scenario 3’s simulated incidence reports. Measurement noise from
the Poisson (no overdispersion) & Negative Binomial distributions
was added to the smooth trajectories obtained from SEIR instances with
an exponential-distributed latent period.

#### 3.2 Inference

##### 3.2.1 Poisson data

###### 3.2.1.1 Fitting \(D^{11}\)

###### 3.2.1.1.1 Incidence fit

Fig 36. Posterior predictive checks against latent incidence. Dots
denote synthetic data while lines indicate simulations from candidates’
ODE structure. We configure these structures using samples from the
posterior distribution.

Fig 37. Fit scores by candidate model and type of data. \(x\) denotes latent incidence, whereas \(y\) indicates observed incidence. Vertical
line denotes the mean.

###### 3.2.1.1.2 \(\Re\_0\) marginal distribution

Fig 38. Estimates of the basic reproduction number by dataset and model
candidate. Error bars denote 95% credible intervals. The value in the
middle of the bars indicates the distance between the marginal
posterior’s mean and the true value (vertical line).

###### 3.2.1.1.3 \(I\_0\) marginal distribution

Fig 39. Estimates of the initial number of infectious individuals by
dataset and model candidate. Error bars denote 95% credible intervals.
The value in the middle of the bars indicates the distance between the
marginal posterior’s mean and the true value (vertical line).

###### 3.2.1.2 Fitting \(D^{12}\)

###### 3.2.1.2.1 Incidence fit

Fig 40. Posterior predictive checks against latent incidence. Dots
denote synthetic data while lines indicate simulations from candidates’
ODE structure. We configure these structures using samples from the
posterior distribution.

Fig 41. Fit scores by candidate model and type of data. \(x\) denotes latent incidence, whereas \(y\) indicates observed incidence. Vertical
line denotes the mean.

###### 3.2.1.2.2 \(\Re\_0\) marginal distribution

Fig 42. Estimates of the basic reproduction number by dataset and model
candidate. Error bars denote 95% credible intervals. The value in the
middle of the bars indicates the distance between the marginal
posterior’s mean and the true value (vertical line).

###### 3.2.1.2.3 \(I\_0\) marginal distribution

Fig 43. Estimates of the initial number of infectious individuals by
dataset and model candidate. Error bars denote 95% credible intervals.
The value in the middle of the bars indicates the distance between the
marginal posterior’s mean and the true value (vertical line).

###### 3.2.1.3 Fitting \(D^{13}\)

###### 3.2.1.3.1 Incidence fit

Fig 45. Fit scores by candidate model and type of data. \(x\) denotes latent incidence, whereas \(y\) indicates observed incidence. Vertical
line denotes the mean.

###### 3.2.1.3.2 \(\Re\_0\) marginal distribution

Fig 46. Estimates of the basic reproduction number by dataset and model
candidate. Error bars denote 95% credible intervals. The value in the
middle of the bars indicates the distance between the marginal
posterior’s mean and the true value (vertical line).

###### 3.2.1.3.3 \(I\_0\) marginal distribution

Fig 47. Estimates of the initial number of infectious individuals by
dataset and model candidate. Error bars denote 95% credible intervals.
The value in the middle of the bars indicates the distance between the
marginal posterior’s mean and the true value (vertical line).

###### 3.2.1.4 Fitting \(D^{14}\)

###### 3.2.1.4.1 Incidence fit

Fig 49. Fit scores by candidate model and type of data. \(x\) denotes latent incidence, whereas \(y\) indicates observed incidence. Vertical
line denotes the mean.

###### 3.2.1.4.2 \(\Re\_0\) marginal distribution

Fig 50. Estimates of the basic reproduction number by dataset and model
candidate. Error bars denote 95% credible intervals. The value in the
middle of the bars indicates the distance between the marginal
posterior’s mean and the true value (vertical line).

###### 3.2.1.4.3 \(I\_0\) marginal distribution

Fig 51. Estimates of the initial number of infectious individuals by
dataset and model candidate. Error bars denote 95% credible intervals.
The value in the middle of the bars indicates the distance between the
marginal posterior’s mean and the true value (vertical line).

###### 3.2.1.5 Summary

###### 3.2.1.5.1 Coverage

Table 4. Scenario 3's coverage table. Poisson noise

| D^ij^ | M^ij^ | ℜ_0_ | I_0_ | ρ |
| --- | --- | --- | --- | --- |
| 11 | 11 | 100% | 100% | 100% |
| 11 | 12 | 95% | 75% | 100% |
| 11 | 13 | 90% | 55% | 100% |
| 11 | 14 | 90% | 45% | 100% |
| 12 | 11 | 95% | 85% | 90% |
| 12 | 12 | 90% | 100% | 90% |
| 12 | 13 | 90% | 90% | 95% |
| 12 | 14 | 90% | 85% | 95% |
| 13 | 11 | 95% | 65% | 100% |
| 13 | 12 | 100% | 95% | 95% |
| 13 | 13 | 100% | 95% | 90% |
| 13 | 14 | 95% | 100% | 90% |
| 14 | 11 | 90% | 75% | 100% |
| 14 | 12 | 95% | 90% | 100% |
| 14 | 13 | 90% | 90% | 100% |
| 14 | 14 | 90% | 90% | 100% |

##### 3.2.2 Overdispersed data

###### 3.2.2.1 Fitting \(D^{11}\)

###### 3.2.2.1.1 Incidence fit

Fig 53. Fit scores by candidate model and type of data. \(x\) denotes latent incidence, whereas \(y\) indicates observed incidence. Vertical
line denotes the mean.

###### 3.2.2.1.2 \(\Re\_0\) marginal distribution

Fig 54. Estimates of the basic reproduction number by dataset and model
candidate. Error bars denote 95% credible intervals. The value in the
middle of the bars indicates the distance between the marginal
posterior’s mean and the true value (vertical line).

###### 3.2.2.1.3 \(I\_0\) marginal distribution

Fig 55. Estimates of the initial number of infectious individuals by
dataset and model candidate. Error bars denote 95% credible intervals.
The value in the middle of the bars indicates the distance between the
marginal posterior’s mean and the true value (vertical line).

###### 3.2.2.2 Fitting \(D^{12}\)

###### 3.2.2.2.1 Incidence fit

Fig 56. Posterior predictive checks against latent incidence. Dots
denote synthetic data while lines indicate simulations from candidates’
ODE structure. We configure these structures using samples from the
posterior distribution.

Fig 57. Fit scores by candidate model and type of data. \(x\) denotes latent incidence, whereas \(y\) indicates observed incidence. Vertical
line denotes the mean.

###### 3.2.2.2.2 \(\Re\_0\) marginal distribution

Fig 58. Estimates of the basic reproduction number by dataset and model
candidate. Error bars denote 95% credible intervals. The value in the
middle of the bars indicates the distance between the marginal
posterior’s mean and the true value (vertical line).

###### 3.2.2.2.3 \(I\_0\) marginal distribution

Fig 59. Estimates of the initial number of infectious individuals by
dataset and model candidate. Error bars denote 95% credible intervals.
The value in the middle of the bars indicates the distance between the
marginal posterior’s mean and the true value (vertical line).

###### 3.2.2.3 Fitting \(D^{13}\)

###### 3.2.2.3.1 Incidence fit

Fig 61. Fit scores by candidate model and type of data. \(x\) denotes latent incidence, whereas \(y\) indicates observed incidence. Vertical
line denotes the mean.

###### 3.2.2.3.2 \(\Re\_0\) marginal distribution

Fig 62. Estimates of the basic reproduction number by dataset and model
candidate. Error bars denote 95% credible intervals. The value in the
middle of the bars indicates the distance between the marginal
posterior’s mean and the true value (vertical line).

###### 3.2.2.3.3 \(I\_0\) marginal distribution

Fig 63. Estimates of the initial number of infectious individuals by
dataset and model candidate. Error bars denote 95% credible intervals.
The value in the middle of the bars indicates the distance between the
marginal posterior’s mean and the true value (vertical line).

###### 3.2.2.4 Fitting \(D^{14}\)

###### 3.2.2.4.1 Incidence fit

Fig 64. Posterior predictive checks against latent incidence. Dots
denote synthetic data while lines indicate simulations from candidates’
ODE structure. We configure these structures using samples from the
posterior distribution.

Fig 65. Fit scores by candidate model and type of data. \(x\) denotes latent incidence, whereas \(y\) indicates observed incidence. Vertical
line denotes the mean.

###### 3.2.2.4.2 \(\Re\_0\) marginal distribution

Fig 66. Estimates of the basic reproduction number by dataset and model
candidate. Error bars denote 95% credible intervals. The value in the
middle of the bars indicates the distance between the marginal
posterior’s mean and the true value (vertical line).

###### 3.2.2.4.3 \(I\_0\) marginal distribution

Fig 67. Estimates of the initial number of infectious individuals by
dataset and model candidate. Error bars denote 95% credible intervals.
The value in the middle of the bars indicates the distance between the
marginal posterior’s mean and the true value (vertical line).

###### 3.2.2.5 Summary

###### 3.2.2.5.1 Coverage

Table 5. Scenario 3's coverage table. Overdispersion

| D^ij^ | M^ij^ | ℜ_0_ | I_0_ | ρ | φ^-1^ |
| --- | --- | --- | --- | --- | --- |
| 11 | 11 | 95% | 95% | 100% | 90% |
| 11 | 12 | 100% | 95% | 95% | 95% |
| 11 | 13 | 90% | 95% | 100% | 95% |
| 11 | 14 | 90% | 90% | 100% | 95% |
| 12 | 11 | 90% | 100% | 100% | 85% |
| 12 | 12 | 95% | 100% | 100% | 90% |
| 12 | 13 | 95% | 90% | 100% | 80% |
| 12 | 14 | 95% | 90% | 100% | 80% |
| 13 | 11 | 100% | 95% | 95% | 90% |
| 13 | 12 | 100% | 100% | 95% | 90% |
| 13 | 13 | 100% | 100% | 95% | 90% |
| 13 | 14 | 100% | 100% | 95% | 85% |
| 14 | 11 | 85% | 90% | 95% | 85% |
| 14 | 12 | 90% | 90% | 95% | 85% |
| 14 | 13 | 90% | 90% | 90% | 85% |
| 14 | 14 | 90% | 90% | 95% | 85% |

### 4 Scenario 4

In comparison to Scenario 1, we increase \(\Re\_0\) from **2.5** to
**9**.

#### 4.1 Synthetic data

Fig 68. Scenario 4’s simulated incidence reports. Measurement noise from
the Poisson (no overdispersion) & Negative Binomial distributions
was added to the smooth trajectories obtained from SEIR instances with
an exponential-distributed latent period.

#### 4.2 Inference

##### 4.2.1 Poisson data

###### 4.2.1.1 Fitting \(D^{11}\)

###### 4.2.1.1.1 Incidence fit

Fig 69. Posterior predictive checks against latent incidence. Dots
denote synthetic data while lines indicate simulations from candidates’
ODE structure. We configure these structures using samples from the
posterior distribution.

Fig 70. Fit scores by candidate model and type of data. \(x\) denotes latent incidence, whereas \(y\) indicates observed incidence. Vertical
line denotes the mean.

###### 4.2.1.1.2 \(\Re\_0\) marginal distribution

Fig 71. Estimates of the basic reproduction number by dataset and model
candidate. Error bars denote 95% credible intervals. The value in the
middle of the bars indicates the distance between the marginal
posterior’s mean and the true value (vertical line).

###### 4.2.1.1.3 \(I\_0\) marginal distribution

Fig 72. Estimates of the initial number of infectious individuals by
dataset and model candidate. Error bars denote 95% credible intervals.
The value in the middle of the bars indicates the distance between the
marginal posterior’s mean and the true value (vertical line).

###### 4.2.1.2 Fitting \(D^{12}\)

###### 4.2.1.2.1 Incidence fit

Fig 73. Posterior predictive checks against latent incidence. Dots
denote synthetic data while lines indicate simulations from candidates’
ODE structure. We configure these structures using samples from the
posterior distribution.

Fig 74. Fit scores by candidate model and type of data. \(x\) denotes latent incidence, whereas \(y\) indicates observed incidence. Vertical
line denotes the mean.

###### 4.2.1.2.2 \(\Re\_0\) marginal distribution

Fig 75. Estimates of the basic reproduction number by dataset and model
candidate. Error bars denote 95% credible intervals. The value in the
middle of the bars indicates the distance between the marginal
posterior’s mean and the true value (vertical line).

###### 4.2.1.2.3 \(I\_0\) marginal distribution

Fig 76. Estimates of the initial number of infectious individuals by
dataset and model candidate. Error bars denote 95% credible intervals.
The value in the middle of the bars indicates the distance between the
marginal posterior’s mean and the true value (vertical line).

###### 4.2.1.3 Fitting \(D^{13}\)

###### 4.2.1.3.1 Incidence fit

Fig 77. Posterior predictive checks against latent incidence. Dots
denote synthetic data while lines indicate simulations from candidates’
ODE structure. We configure these structures using samples from the
posterior distribution.

Fig 78. Fit scores by candidate model and type of data. \(x\) denotes latent incidence, whereas \(y\) indicates observed incidence. Vertical
line denotes the mean.

###### 4.2.1.3.2 \(\Re\_0\) marginal distribution

Fig 79. Estimates of the basic reproduction number by dataset and model
candidate. Error bars denote 95% credible intervals. The value in the
middle of the bars indicates the distance between the marginal
posterior’s mean and the true value (vertical line).

###### 4.2.1.3.3 \(I\_0\) marginal distribution

Fig 80. Estimates of the initial number of infectious individuals by
dataset and model candidate. Error bars denote 95% credible intervals.
The value in the middle of the bars indicates the distance between the
marginal posterior’s mean and the true value (vertical line).

###### 4.2.1.4 Fitting \(D^{14}\)

###### 4.2.1.4.1 Incidence fit

Fig 81. Posterior predictive checks against latent incidence. Dots
denote synthetic data while lines indicate simulations from candidates’
ODE structure. We configure these structures using samples from the
posterior distribution.

Fig 82. Fit scores by candidate model and type of data. \(x\) denotes latent incidence, whereas \(y\) indicates observed incidence. Vertical
line denotes the mean.

###### 4.2.1.4.2 \(\Re\_0\) marginal distribution

Fig 83. Estimates of the basic reproduction number by dataset and model
candidate. Error bars denote 95% credible intervals. The value in the
middle of the bars indicates the distance between the marginal
posterior’s mean and the true value (vertical line).

###### 4.2.1.4.3 \(I\_0\) marginal distribution

Fig 84. Estimates of the initial number of infectious individuals by
dataset and model candidate. Error bars denote 95% credible intervals.
The value in the middle of the bars indicates the distance between the
marginal posterior’s mean and the true value (vertical line).

###### 4.2.1.5 Summary

###### 4.2.1.5.1 Coverage

Table 6. Scenario 4's coverage table. Poisson noise

| D^ij^ | M^ij^ | ℜ_0_ | I_0_ | ρ |
| --- | --- | --- | --- | --- |
| 11 | 11 | 95% | 95% | 80% |
| 11 | 12 | 10% | 90% | 80% |
| 11 | 13 | 0% | 80% | 80% |
| 11 | 14 | 0% | 65% | 80% |
| 12 | 11 | 45% | 70% | 95% |
| 12 | 12 | 90% | 85% | 95% |
| 12 | 13 | 55% | 90% | 95% |
| 12 | 14 | 35% | 85% | 95% |
| 13 | 11 | 10% | 75% | 100% |
| 13 | 12 | 80% | 90% | 100% |
| 13 | 13 | 95% | 90% | 100% |
| 13 | 14 | 90% | 95% | 100% |
| 14 | 11 | 0% | 50% | 85% |
| 14 | 12 | 60% | 85% | 90% |
| 14 | 13 | 100% | 95% | 90% |
| 14 | 14 | 100% | 100% | 80% |

##### 4.2.2 Overdispersed data

###### 4.2.2.1 Fitting \(D^{11}\)

###### 4.2.2.2 Incidence fit

Fig 85. Posterior predictive checks against latent incidence. Dots
denote synthetic data while lines indicate simulations from candidates’
ODE structure. We configure these structures using samples from the
posterior distribution.

Fig 86. Fit scores by candidate model and type of data. \(x\) denotes latent incidence, whereas \(y\) indicates observed incidence. Vertical
line denotes the mean.

###### 4.2.2.2.1 \(\Re\_0\) marginal distribution

Fig 87. Estimates of the basic reproduction number by dataset and model
candidate. Error bars denote 95% credible intervals. The value in the
middle of the bars indicates the distance between the marginal
posterior’s mean and the true value (vertical line).

###### 4.2.2.2.2 \(I\_0\) marginal distribution

Fig 88. Estimates of the initial number of infectious individuals by
dataset and model candidate. Error bars denote 95% credible intervals.
The value in the middle of the bars indicates the distance between the
marginal posterior’s mean and the true value (vertical line).

###### 4.2.2.3 Fitting \(D^{12}\)

###### 4.2.2.3.1 Incidence fit

Fig 89. Posterior predictive checks against latent incidence. Dots
denote synthetic data while lines indicate simulations from candidates’
ODE structure. We configure these structures using samples from the
posterior distribution.

Fig 90. Fit scores by candidate model and type of data. \(x\) denotes latent incidence, whereas \(y\) indicates observed incidence. Vertical
line denotes the mean.

###### 4.2.2.3.2 \(\Re\_0\) marginal distribution

Fig 91. Estimates of the basic reproduction number by dataset and model
candidate. Error bars denote 95% credible intervals. The value in the
middle of the bars indicates the distance between the marginal
posterior’s mean and the true value (vertical line).

###### 4.2.2.3.3 \(I\_0\) marginal distribution

Fig 92. Estimates of the initial number of infectious individuals by
dataset and model candidate. Error bars denote 95% credible intervals.
The value in the middle of the bars indicates the distance between the
marginal posterior’s mean and the true value (vertical line).

###### 4.2.2.4 Fitting \(D^{13}\)

###### 4.2.2.4.1 Incidence fit

Fig 93. Posterior predictive checks against latent incidence. Dots
denote synthetic data while lines indicate simulations from candidates’
ODE structure. We configure these structures using samples from the
posterior distribution.

Fig 94. Fit scores by candidate model and type of data. \(x\) denotes latent incidence, whereas \(y\) indicates observed incidence. Vertical
line denotes the mean.

###### 4.2.2.4.2 \(\Re\_0\) marginal distribution

Fig 95. Estimates of the basic reproduction number by dataset and model
candidate. Error bars denote 95% credible intervals. The value in the
middle of the bars indicates the distance between the marginal
posterior’s mean and the true value (vertical line).

###### 4.2.2.4.3 \(I\_0\) marginal distribution

Fig 96. Estimates of the initial number of infectious individuals by
dataset and model candidate. Error bars denote 95% credible intervals.
The value in the middle of the bars indicates the distance between the
marginal posterior’s mean and the true value (vertical line).

###### 4.2.2.5 Fitting \(D^{14}\)

###### 4.2.2.5.1 Incidece fit

Fig 97. Posterior predictive checks against latent incidence. Dots
denote synthetic data while lines indicate simulations from candidates’
ODE structure. We configure these structures using samples from the
posterior distribution.

Fig 98. Fit scores by candidate model and type of data. \(x\) denotes latent incidence, whereas \(y\) indicates observed incidence. Vertical
line denotes the mean.

###### 4.2.2.5.2 \(\Re\_0\) marginal distribution

Fig 99. Estimates of the basic reproduction number by dataset and model
candidate. Error bars denote 95% credible intervals. The value in the
middle of the bars indicates the distance between the marginal
posterior’s mean and the true value (vertical line).

###### 4.2.2.5.3 \(I\_0\) marginal distribution

Fig 100. Estimates of the initial number of infectious individuals by
dataset and model candidate. Error bars denote 95% credible intervals.
The value in the middle of the bars indicates the distance between the
marginal posterior’s mean and the true value (vertical line).

###### 4.2.2.6 Summary

###### 4.2.2.6.1 Coverage

Table 7. Scenario 4's coverage table. Overdispersion

| D^ij^ | M^ij^ | ℜ_0_ | I_0_ | ρ | φ^-1^ |
| --- | --- | --- | --- | --- | --- |
| 11 | 11 | 95% | 95% | 100% | 80% |
| 11 | 12 | 95% | 95% | 100% | 90% |
| 11 | 13 | 90% | 95% | 100% | 80% |
| 11 | 14 | 80% | 95% | 100% | 90% |
| 12 | 11 | 85% | 100% | 85% | 90% |
| 12 | 12 | 90% | 95% | 95% | 90% |
| 12 | 13 | 95% | 95% | 90% | 90% |
| 12 | 14 | 95% | 95% | 90% | 90% |
| 13 | 11 | 75% | 100% | 100% | 95% |
| 13 | 12 | 90% | 100% | 100% | 95% |
| 13 | 13 | 95% | 100% | 100% | 95% |
| 13 | 14 | 100% | 100% | 100% | 95% |
| 14 | 11 | 85% | 100% | 100% | 100% |
| 14 | 12 | 100% | 100% | 100% | 100% |
| 14 | 13 | 100% | 100% | 100% | 100% |
| 14 | 14 | 100% | 100% | 100% | 95% |

### 5 Scenario 5

In comparison to Scenario 4, we increase \(\Re\_0\) from **9** to
**17**.

#### 5.1 Synthetic data

Fig 101. Scenario 5’s simulated incidence reports. Measurement noise
from the Poisson (no overdispersion) & Negative Binomial
distributions was added to the smooth trajectories obtained from SEIR
instances with an exponential-distributed latent period.

#### 5.2 Inference

##### 5.2.1 Poisson data

###### 5.2.1.1 Fitting \(D^{11}\)

###### 5.2.1.1.1 Incidence fit

Fig 102. Posterior predictive checks against latent incidence. Dots
denote synthetic data while lines indicate simulations from candidates’
ODE structure. We configure these structures using samples from the
posterior distribution.

Fig 103. Fit scores by candidate model and type of data. \(x\) denotes latent incidence, whereas \(y\) indicates observed incidence. Vertical
line denotes the mean.

###### 5.2.1.1.2 \(\Re\_0\) marginal distribution

Fig 104. Estimates of the basic reproduction number by dataset and model
candidate. Error bars denote 95% credible intervals. The value in the
middle of the bars indicates the distance between the marginal
posterior’s mean and the true value (vertical line).

###### 5.2.1.1.3 \(I\_0\) marginal distribution

Fig 105. Estimates of the initial number of infectious individuals by
dataset and model candidate. Error bars denote 95% credible intervals.
The value in the middle of the bars indicates the distance between the
marginal posterior’s mean and the true value (vertical line).

###### 5.2.1.2 Fitting \(D^{12}\)

###### 5.2.1.2.1 Incidence fit

Fig 107. Fit scores by candidate model and type of data. \(x\) denotes latent incidence, whereas \(y\) indicates observed incidence. Vertical
line denotes the mean.

###### 5.2.1.2.2 \(\Re\_0\) marginal distribution

Fig 108. Estimates of the basic reproduction number by dataset and model
candidate. Error bars denote 95% credible intervals. The value in the
middle of the bars indicates the distance between the marginal
posterior’s mean and the true value (vertical line).

###### 5.2.1.2.3 \(I\_0\) marginal distribution

Fig 109. Estimates of the initial number of infectious individuals by
dataset and model candidate. Error bars denote 95% credible intervals.
The value in the middle of the bars indicates the distance between the
marginal posterior’s mean and the true value (vertical line).

###### 5.2.1.3 Fitting \(D^{13}\)

###### 5.2.1.3.1 Incidence fit

Fig 110. Posterior predictive checks against latent incidence. Dots
denote synthetic data while lines indicate simulations from candidates’
ODE structure. We configure these structures using samples from the
posterior distribution.

Fig 111. Fit scores by candidate model and type of data. \(x\) denotes latent incidence, whereas \(y\) indicates observed incidence. Vertical
line denotes the mean.

###### 5.2.1.3.2 \(\Re\_0\) marginal distribution

Fig 112. Estimates of the basic reproduction number by dataset and model
candidate. Error bars denote 95% credible intervals. The value in the
middle of the bars indicates the distance between the marginal
posterior’s mean and the true value (vertical line).

###### 5.2.1.3.3 \(I\_0\) marginal distribution

Fig 113. Estimates of the initial number of infectious individuals by
dataset and model candidate. Error bars denote 95% credible intervals.
The value in the middle of the bars indicates the distance between the
marginal posterior’s mean and the true value (vertical line).

###### 5.2.1.4 Fitting \(D^{14}\)

###### 5.2.1.4.1 Incidence fit

Fig 114. Posterior predictive checks against latent incidence. Dots
denote synthetic data while lines indicate simulations from candidates’
ODE structure. We configure these structures using samples from the
posterior distribution.

Fig 115. Fit scores by candidate model and type of data. \(x\) denotes latent incidence, whereas \(y\) indicates observed incidence. Vertical
line denotes the mean.

###### 5.2.1.4.2 \(\Re\_0\) marginal distribution

Fig 116. Estimates of the basic reproduction number by dataset and model
candidate. Error bars denote 95% credible intervals. The value in the
middle of the bars indicates the distance between the marginal
posterior’s mean and the true value (vertical line).

###### 5.2.1.4.3 \(I\_0\) marginal distribution

Fig 117. Estimates of the initial number of infectious individuals by
dataset and model candidate. Error bars denote 95% credible intervals.
The value in the middle of the bars indicates the distance between the
marginal posterior’s mean and the true value (vertical line).

###### 5.2.1.5 Summary

###### 5.2.1.5.1 Coverage

Table 8. Scenario 5's coverage table. Poisson

| D^ij^ | M^ij^ | ℜ_0_ | I_0_ | ρ |
| --- | --- | --- | --- | --- |
| 11 | 11 | 100% | 100% | 95% |
| 11 | 12 | 0% | 100% | 95% |
| 11 | 13 | 0% | 100% | 95% |
| 11 | 14 | 0% | 100% | 95% |
| 12 | 11 | 25% | 90% | 95% |
| 12 | 12 | 95% | 95% | 95% |
| 12 | 13 | 60% | 100% | 95% |
| 12 | 14 | 30% | 95% | 95% |
| 13 | 11 | 0% | 95% | 100% |
| 13 | 12 | 60% | 100% | 100% |
| 13 | 13 | 95% | 100% | 100% |
| 13 | 14 | 95% | 90% | 100% |
| 14 | 11 | 0% | 100% | 100% |
| 14 | 12 | 25% | 100% | 100% |
| 14 | 13 | 85% | 95% | 100% |
| 14 | 14 | 95% | 95% | 100% |

##### 5.2.2 Overdispersed data

#### 5.2.2.1 \(D^{11}\)

###### 5.2.2.1.1 Incidence fit

Fig 118. Posterior predictive checks against latent incidence. Dots
denote synthetic data while lines indicate simulations from candidates’
ODE structure. We configure these structures using samples from the
posterior distribution.

Fig 119. Fit scores by candidate model and type of data. \(x\) denotes latent incidence, whereas \(y\) indicates observed incidence. Vertical
line denotes the mean.

###### 5.2.2.1.2 \(\Re\_0\) marginal distribution

Fig 120. Estimates of the basic reproduction number by dataset and model
candidate. Error bars denote 95% credible intervals. The value in the
middle of the bars indicates the distance between the marginal
posterior’s mean and the true value (vertical line).

###### 5.2.2.1.3 \(I\_0\) marginal distribution

Fig 121. Estimates of the initial number of infectious individuals by
dataset and model candidate. Error bars denote 95% credible intervals.
The value in the middle of the bars indicates the distance between the
marginal posterior’s mean and the true value (vertical line).

###### 5.2.2.2 Fitting \(D^{12}\)

###### 5.2.2.2.1 Incidence fit

Fig 122. Posterior predictive checks against latent incidence. Dots
denote synthetic data while lines indicate simulations from candidates’
ODE structure. We configure these structures using samples from the
posterior distribution.

Fig 123. Fit scores by candidate model and type of data. \(x\) denotes latent incidence, whereas \(y\) indicates observed incidence. Vertical
line denotes the mean.

###### 5.2.2.2.2 \(\Re\_0\) marginal distribution

Fig 124. Estimates of the basic reproduction number by dataset and model
candidate. Error bars denote 95% credible intervals. The value in the
middle of the bars indicates the distance between the marginal
posterior’s mean and the true value (vertical line).

###### 5.2.2.2.3 \(I\_0\) marginal distribution

Fig 125. Estimates of the initial number of infectious individuals by
dataset and model candidate. Error bars denote 95% credible intervals.
The value in the middle of the bars indicates the distance between the
marginal posterior’s mean and the true value (vertical line).

###### 5.2.2.3 Fitting \(D^{13}\)

###### 5.2.2.3.1 Incidence fit

Fig 126. Posterior predictive checks against latent incidence. Dots
denote synthetic data while lines indicate simulations from candidates’
ODE structure. We configure these structures using samples from the
posterior distribution.

Fig 127. Fit scores by candidate model and type of data. \(x\) denotes latent incidence, whereas \(y\) indicates observed incidence. Vertical
line denotes the mean.

###### 5.2.2.3.2 \(\Re\_0\) marginal distribution

Fig 128. Estimates of the basic reproduction number by dataset and model
candidate. Error bars denote 95% credible intervals. The value in the
middle of the bars indicates the distance between the marginal
posterior’s mean and the true value (vertical line).

###### 5.2.2.3.3 \(I\_0\) marginal distribution

Fig 129. Estimates of the initial number of infectious individuals by
dataset and model candidate. Error bars denote 95% credible intervals.
The value in the middle of the bars indicates the distance between the
marginal posterior’s mean and the true value (vertical line).

###### 5.2.2.4 Fitting \(D^{14}\)

###### 5.2.2.4.1 Incidence fit

Fig 130. Posterior predictive checks against latent incidence. Dots
denote synthetic data while lines indicate simulations from candidates’
ODE structure. We configure these structures using samples from the
posterior distribution.

Fig 131. Fit scores by candidate model and type of data. \(x\) denotes latent incidence, whereas \(y\) indicates observed incidence. Vertical
line denotes the mean.

###### 5.2.2.4.2 \(\Re\_0\) marginal distribution

Fig 132. Estimates of the basic reproduction number by dataset and model
candidate. Error bars denote 95% credible intervals. The value in the
middle of the bars indicates the distance between the marginal
posterior’s mean and the true value (vertical line).

###### 5.2.2.4.3 \(I\_0\) marginal distribution

Fig 133. Estimates of the initial number of infectious individuals by
dataset and model candidate. Error bars denote 95% credible intervals.
The value in the middle of the bars indicates the distance between the
marginal posterior’s mean and the true value (vertical line).

###### 5.2.2.5 Summary

###### 5.2.2.5.1 Coverage

Table 9. Scenario 5's coverage table. Overdispersion

| D^ij^ | M^ij^ | ℜ_0_ | I_0_ | ρ | φ^-1^ |
| --- | --- | --- | --- | --- | --- |
| 11 | 11 | 100% | 100% | 100% | 100% |
| 11 | 12 | 100% | 100% | 100% | 100% |
| 11 | 13 | 95% | 100% | 100% | 100% |
| 11 | 14 | 80% | 100% | 100% | 100% |
| 12 | 11 | 80% | 95% | 100% | 95% |
| 12 | 12 | 100% | 95% | 100% | 100% |
| 12 | 13 | 100% | 90% | 100% | 95% |
| 12 | 14 | 95% | 90% | 100% | 95% |
| 13 | 11 | 70% | 100% | 100% | 90% |
| 13 | 12 | 95% | 100% | 95% | 90% |
| 13 | 13 | 95% | 100% | 100% | 90% |
| 13 | 14 | 100% | 100% | 95% | 90% |
| 14 | 11 | 65% | 95% | 100% | 100% |
| 14 | 12 | 90% | 95% | 100% | 100% |
| 14 | 13 | 90% | 95% | 100% | 100% |
| 14 | 14 | 95% | 95% | 100% | 100% |
