## Supplementary material for "Anchoring the mean generation time in the SEIR to mitigate biases in ℜ_0_ estimates due to uncertainty in the distribution of the epidemiological delays": S7.html


# S7

In this appendix, we illustrate the procedure to infer the basic
reproduction number (\(\Re\_0\)) of
influenza during the second wave of the **1918 pandemic**
in Cumberland (Maryland). Specifically, we fit four model candidates
from the **alternative parameterisation** to incidence
data. The inference process yields almost identical \(\Re\_0\) estimates, regardless of the
infectious period distribution. Furthermore, we compare the results of
the alternative parameterisation to those of the *traditional*
one.

### 1 Incidence data

After enduring a wave of influenza infections during the spring of
1918, the U.S. Public Health Service organised special surveys in
several localities to determine as accurately as possible the proportion
of the population infected during the second wave of infections in the
autumn of 1918. In the figure below, we show the report of new cases
detected in the city of Cumberland (Maryland) over that period.

Fig 1. Cumberland’s incidence data

### 2 Inference

We employ four candidates per parameterisation (traditional and
alternative). On the one hand, the traditional parameterisation refers
to the approach of fixing the mean of the epidemiological delays (latent
and infectious periods) to values obtained from the literature,
irrespective of their particular distribution. On the other hand, the
proposed alternative parameterisation refers to the special emphasis
placed on mean generation time of the **SEIR**, while
flexibilising the mean and distribution of the epidemiological delays.
Namely, the epidemiological delays can take any mean or shape provided
that as a whole conform to the observed mean generation time.
Furthermore, following the results shown in **S4** and
**S5**, for all candidates, we assume an
exponentially-distributed latent period (\(SE^1I^jR\)), where \(j = \{1,2,3,4\}\).

#### 2.1 Prior distribution

Prior distributions for both parameterisations correspond to those
employed in *S3*.

#### 2.2 Posterior distribution

##### 2.2.1 Incidence fit

Fig 2. Posterior predictive checks

##### 2.2.2 Joint posterior distribution

##### 2.2.3 Marginal distributions

###### 2.2.3.1 Basic reproduction number (\(\Re\_0\))

Fig 3. Estimation of the basic reproduction number by model candidate
and parameterisation

###### 2.2.3.2 Reporting rate (\(\rho\))

Fig 4. Estimation of the reporting rate by model candidate and
parameterisation

###### 2.2.3.3 Initial number of infectious individuals (\(I\_0\))

Fig 5. Estimation of the initial number of infectious individuals by
model candidate and parameterisation

###### 2.2.3.4 Overdispersion parameter (\(\phi^{-1}\))

Fig 6. Estimation of overdispersion by model candidate and
parameterisation
